# Predicting the antigenic evolution of seasonal influenza viruses using phylogenetic convergence

**DOI:** 10.64898/2026.04.10.717627

**Authors:** Samuel A. Turner, David J. Pattinson, Ron A. M. Fouchier, Derek J. Smith

## Abstract

The antigenic evolution of human seasonal influenza viruses is primarily driven by single amino acid substitutions immediately adjacent to the receptor binding site in the hemagglutinin (HA) protein. The ability to predict these substitutions would allow vaccine strains to be selected with an understanding of likely future antigenic variation. Here, we estimate the effect of HA substitutions on viral fitness using measurements of convergent evolution in a large phylogeny. We show that the substitutions which have historically caused major antigenic changes in H3N2 influenza viruses were nearly always one of few substitutions near the HA receptor binding site estimated to be under positive selection in sequences collected before the antigenic transition, based on convergent acquisition of the substitution in multiple independent lineages. Furthermore, this signal predates the establishment of the major clade containing the antigenic substitution by more than one year, so is highly informative for prospective prediction.

## Main Text

Vaccination is the primary tool used to counter the substantial economic and public health burden caused by seasonal influenza viruses. Protection is conferred mainly by antibodies against hemagglutinin (HA), one of the influenza virus surface proteins. However, influenza virus evolves to escape antibody recognition by acquiring mutations in HA in a process of punctuated antigenic evolution (*1*). It is therefore important that the HA contained in the vaccine is well matched to the predominant circulating HA each year, as mismatches are associated with reduced vaccine effectiveness (*2*–*4*). Achieving this is challenging because vaccine strains must be selected ∼9 months before the influenza season, to allow time to manufacture, distribute, and administer vaccines (*5*).

There is therefore substantial interest in predicting the antigenic evolution of seasonal influenza viruses, which would allow vaccine strains to be selected with an understanding of the variants likely to circulate in the future. Existing methods aim to predict the outcome of competition between circulating variants, using fitness estimates based on changes in variant frequencies (*6*), the shape of the phylogenetic tree (*7*), and the number of mutations at amino acid positions which have historically been subject to positive selection as measured by dN/dS (*8*), or which are in epitope and non-epitope positions (*9*). These methods share the limitation that they do not attempt to predict the emergence of future antigenic variants (*10*).

Here, we test whether the substitutions which cause major antigenic changes in H3N2 viruses can be predicted *before* the major clade containing the substitution emerges in nature, by estimating the effects of individual HA substitutions on viral fitness. This is made more tractable by the observation that the major antigenic transitions of H3N2 viruses between 1968 and 2003 were primarily caused by amino acid substitutions at seven positions surrounding the HA receptor binding site (*11*), in contrast to the 131 positions traditionally considered to determine antigenic phenotype (*12*–*14*). The ability to predict amino acid substitutions at these seven key antigenic positions would provide an earlier indication of the substitutions most likely to cause major antigenic changes in the future, which could be used to inform preemptive vaccine updates and avoid the out-of-date vaccine mismatches that reduce vaccine effectiveness.

Previous work has shown that adaptive amino acid substitutions arise multiple times during periods of adaptive evolution of H3N2 virus (*15*), and that substitutions which arise multiple times on the global phylogeny (*16*) or in chronically infected patients (*17*) are more likely to subsequently become fixed, suggesting that convergence holds information about the evolutionary fate of substitutions.

Here, we systematically measure this convergent evolution to infer the direction and strength of selection for each amino acid substitution, using an independently derived method related to that previously applied to SARS-CoV-2 (*18*, *19*). This extends existing site-specific dN/dS methods, which have long been applied to influenza viruses and other organisms (*20*–*25*), providing measurements of selection at the level of individual amino acid substitutions, rather than positions—critical for predicting the evolution of specific antigenic variants.

## Results

### Estimating effects of amino acid substitutions on viral fitness

We first constructed a maximum-likelihood phylogenetic tree of all HA1 nucleotide sequences from GISAID (*26*) using IQTREE (*27*) and CMAPLE (*28*, *29*), and annotated each branch with the nucleotide mutations estimated to have occurred on it using UShER (*30*). To estimate selection acting on an amino acid substitution, we count the number of independent phylogenetic occurrences of each nucleotide mutation which produces that amino acid substitution (Fig. 1A). However, different types of nucleotide mutation occur at substantially different rates, even when synonymous (*31*–*33*): for example, we find that GA occurs >50x more often than CG on average (Fig. 1B). To control for these rate differences, we compare the observed number of occurrences of an amino acid substitution against a neutral expectation derived from mutation rates at four-fold-synonymous positions (where the same amino acid is encoded regardless of the nucleotide present at the position, meaning every nucleotide mutation is synonymous) (Fig. 1C). We divide the observed by the neutral expected number of occurrences to give a “convergence ratio”, and take log_2_ of this value to produce the “log convergence ratio” (LCR).

**Fig. 1.**
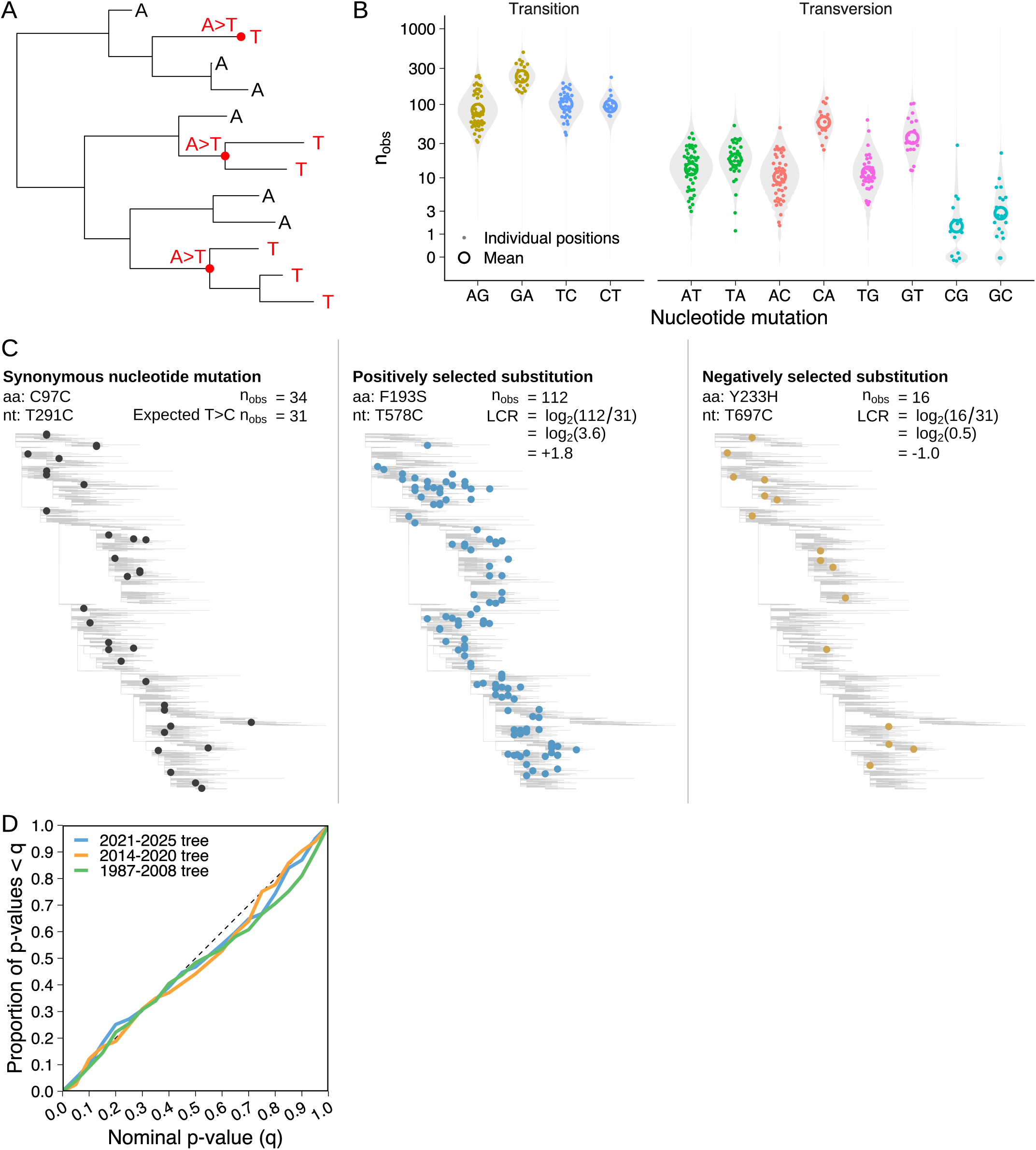
Effects of HA substitutions on viral fitness are estimated by measuring convergent evolution. (**A**) An example phylogenetic tree containing three independent occurrences of an AT mutation. (**B**) The number of independent occurrences of each type of nucleotide mutation at four-fold-synonymous HA1 positions in the predominant group of H3N2 viruses since 2021 (i.e. the Darwin 2021 antigenic cluster). Grey violin plots show the fitted distribution of synonymous mutation counts. (**C**) Example calculation of the log convergence ratio (LCR) for a positively selected amino acid substitution (F193S) and a negatively selected substitution (H233Y) in the predominant group of H3N2 viruses between 2014 and 2020 (i.e. the Hong Kong 2014 antigenic cluster). In practice, LCR estimates for an amino acid substitution sum over all possible causative nucleotide mutations, and consider only the branches of the tree where each mutation would cause the amino acid substitution in question. See Methods section *Calculating the expected number of occurrences* for details. (**D**) Distribution of empirical p-values for synonymous mutations in three independent phylogenetic trees, using model parameters fit to the 2021-2025 tree for all three trees. See Methods section *Calculating empirical p-values* for details.

A positive LCR therefore indicates that an amino acid substitution occurs more often than expected under neutral evolution, so positive selection is inferred; and vice versa for a negative value. To check whether LCR estimates were sensitive to the method used to construct the tree, we constructed a second tree by maximum-parsimony using IQTREE, UShER and matOptimize (*34*), and find that estimates are highly concordant (Fig. S1).

### Estimating statistical significance of observed mutation counts

To determine whether a particular observed number of occurrences represents a statistically significant deviation from neutral evolution, we use a null model of evolution containing two sources of variation: Poisson distributed sampling variation, caused by stochasticity in the evolutionary and sequence sampling processes (see Methods section *Testing the Poisson model of sampling variation* and Fig. S2); and normally distributed variation in the synonymous mutation rate between positions (see Methods section *Calculating the expected number of occurrences*, Supplementary text section *Synonymous mutation rate variation,* and Figs. S3 & S4). To test the suitability of the model, we fitted it to a tree containing sequences from 2021-2025, and calculated empirical p-values for synonymous mutations in both the 2021-2025 tree and in two out-of-sample trees, containing sequences from 2014-2020 and 1987-2008 respectively. Despite using parameter values fit to the 2021-2025 tree, the empirical p-values were approximately uniformly distributed for all three trees, indicating they are well-calibrated (Fig. 1D, Methods section *Calculating empirical p-values*).

### Substitutions contributing to the long-term sequence evolution of HA are positively selected

We find that most amino acid substitutions are negatively selected, indicating they are deleterious (Fig. 2A), mirroring findings from experimental work on HA (*35*) and on the SARS-CoV-2 spike protein (*36*), and similar estimates of the fitness effect of substitutions to SARS-CoV-2 proteins (*18*). Accordingly, many of these substitutions have never been observed on the tree. Of those which are observed, we find that those present only on minor branches of the tree (i.e. not on the trunk or leading to an antigenic cluster, described below) are on average approximately neutral, and that many are slightly deleterious, particularly outside the antigenic sites—consistent with previous findings that influenza virus carries a deleterious mutation load which drives the extinction of non-trunk lineages (*9*, *37*, *38*). By contrast, substitutions on the trunk and major branches (i.e. branches leading to an antigenic cluster) are typically positively selected (Figs. 2A & 2B), particularly in the antigenic sites nearest to the receptor binding site, A, B, and D (*9*, *39*). We find that a small number of trunk and major branch substitutions are neutral or slightly deleterious, suggesting that while linkage (*40*) or genetic drift during transmission bottlenecks and the seeding of epidemics (*41*) can allow non-adaptive substitutions to reach high frequency, this occurs relatively rarely.

**Fig. 2.**
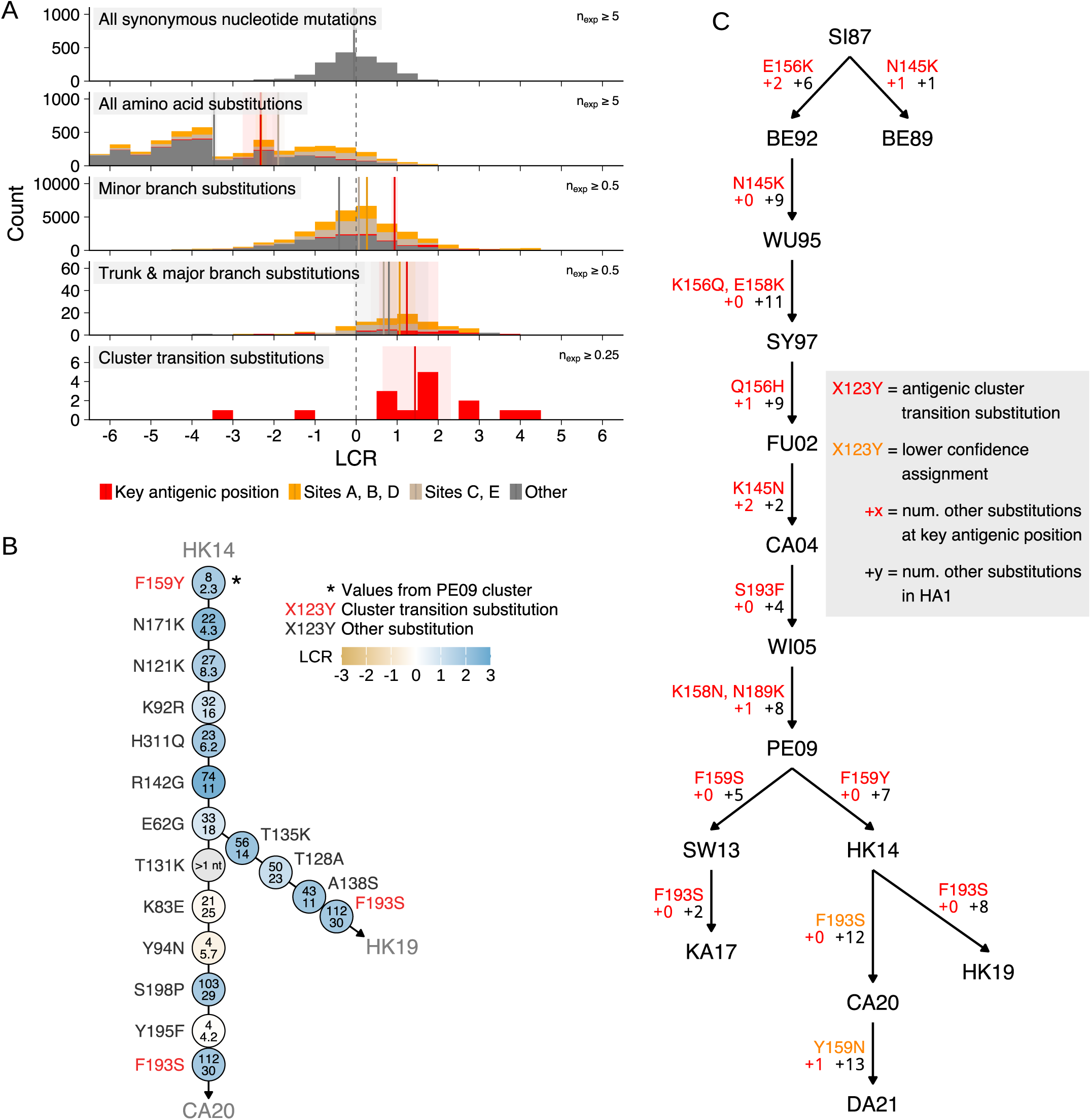
Substitutions on the trunk or leading to an antigenic cluster are typically positively selected. (**A**) Distribution of log convergence ratios (LCR) for synonymous nucleotide mutations, all amino acid substitutions accessible with a single nucleotide change, substitutions occurring on minor branches (i.e. not on the trunk or leading to an antigenic cluster), substitutions occurring on the trunk & major branches (i.e. leading to an antigenic cluster), and antigenic cluster transition substitutions. Coloring corresponds to the antigenic sites or key antigenic positions from Koel et al. 2013 (*11*). Solid vertical lines indicate the mean, surrounded by shaded 95% bootstrap confidence intervals. n_obs_ and n_exp_ are the observed and neutral expected number of occurrences respectively. (**B**) LCRs of substitutions occurring on the trunk & major branches between 2014 and 2020 (i.e. in the Hong Kong 2014 antigenic cluster). The observed (upper number) and neutral expected (lower number) number of occurrences are given for each substitution. (**C**) Substitutions estimated to be responsible for antigenic cluster transitions between 1987 and 2024. See Supplementary text section *Assignment of antigenic clusters and causative cluster transition substitutions* and Figs. S6 & S7 for more details, and Fig. S5 for antigenic cluster transitions predating 1987.

### Convergent evolution predicts substitutions which cause antigenic cluster transitions

When forecasting the antigenic evolution of influenza virus, some substitutions are more important to predict than others. The antigenic evolution of influenza virus is organized into clusters of cross-reactive, antigenically similar viruses, with large antigenic changes seeding a new antigenic cluster (an “antigenic cluster transition”) every three years on average for H3N2 (*1*). Koel et al. 2013 showed that the major antigenic change at such cluster transitions is caused by substitutions at seven key amino acid positions near to the receptor binding site (“antigenic cluster transition substitutions”)—H3 HA amino acid positions 145, 155, 156, 158, 159, 189, and 193 (*11*). Subsequent work has shown that the antigenic evolution of other subtypes of influenza virus in humans (*42*, *43*) and other species (*44*–*47*) is also largely determined by changes at the same or proximal positions near the HA receptor binding site.

Here, we attempt to prospectively identify antigenic cluster transition substitutions by measuring the LCR of substitutions at the seven key antigenic positions identified in Koel et al. 2013 (Figs. 2C & S5; a summary of the studies from which these estimates of antigenic cluster transition substitutions were obtained (*48*–*60*) is provided in Supplementary text section *Assignment of antigenic clusters and causative cluster transition substitutions* and Figs. S6 & S7). Importantly, we make LCR measurements without using sequences from the descendant cluster, via which the cluster transition substitution ultimately reaches high frequency.

For example, HA substitution F159Y is responsible for the antigenic cluster transition from the Perth 2009 (PE09) cluster to the Hong Kong 2014 (HK14) cluster (antigenic clusters are named after the first WHO recommended vaccine strain in the cluster, and abbreviated by location and year; so the cluster containing the Perth/16/2009 vaccine strain is Perth 2009, or PE09) (*53*, *54*). F159Y occurs multiple times in the ancestral PE09 cluster before the occurrence which founds the HK14 cluster (Fig. 3A, Fig. S8 for other cluster transition substitutions). Using sequences from the ancestral PE09 cluster only, we measured the LCR of all amino acid substitutions accessible by a single nucleotide change at the seven key antigenic positions (Fig. 3B): F159Y placed 2^nd^ out of 39 when the substitutions were ranked by LCR (Fig. 3C, Fig. S9 for other antigenic clusters).

**Fig. 3.**
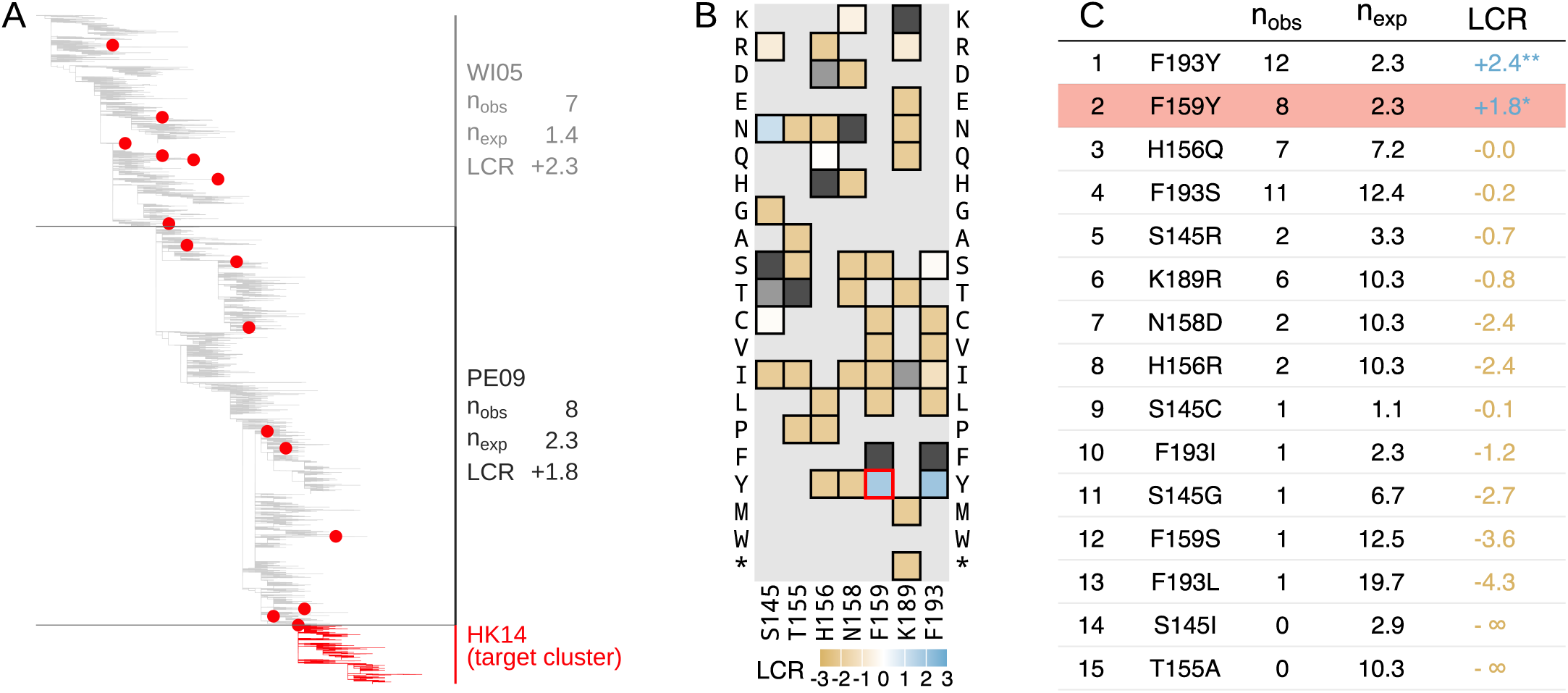
The F159Y substitution was highly convergent prior to causing an antigenic cluster transition. (**A**) Phylogenetic tree showing WI05, PE09, and start of HK14 antigenic clusters, marked with occurrences of the F159Y substitution, which caused the antigenic cluster transition from PE09 to HK14. n_obs_ and n_exp_ are the observed and neutral expected number of occurrences respectively. Fig. S8 shows equivalent trees for other cluster transition substitutions. (**B**) Log convergence ratios (LCR) of substitutions at key antigenic positions in PE09. Dark grey squares identify the ancestral amino acid at the position, and light grey squares show substitutions where neither the observed nor neutral expected number of occurrences exceed one. (**C**) Ranking of substitutions from **B** (see Methods section *Calculating ranks* for details). Fig. S9 shows equivalent tables for other antigenic cluster transitions. * = p<0.05, **= p<0.01, *** = p<0.001. F159Y is highlighted in **B** with a red outline and in **C** with red shading.

We repeated this analysis for each antigenic cluster transition since 1987 (Fig. 4A, Fig. S10 for earlier clusters back to 1968, excluded from the main analysis due to the small number of available sequences). Out of 14 cluster transition substitutions caused by single nucleotide changes since 1987, only one had a negative LCR in the ancestral cluster (F159S, causing PE09 to SW13), indicating negative selection. In 12 out of 14 cases, the antigenic cluster transition substitution was ranked as one of the top six substitutions by the LCR, with a median ranking of 3^rd^. The rankings of cluster transition substitutions were highly consistent between the trees produced using CMAPLE and UShER + matOptimize, differing by at most one ranking position (Fig. S11).

**Fig. 4.**
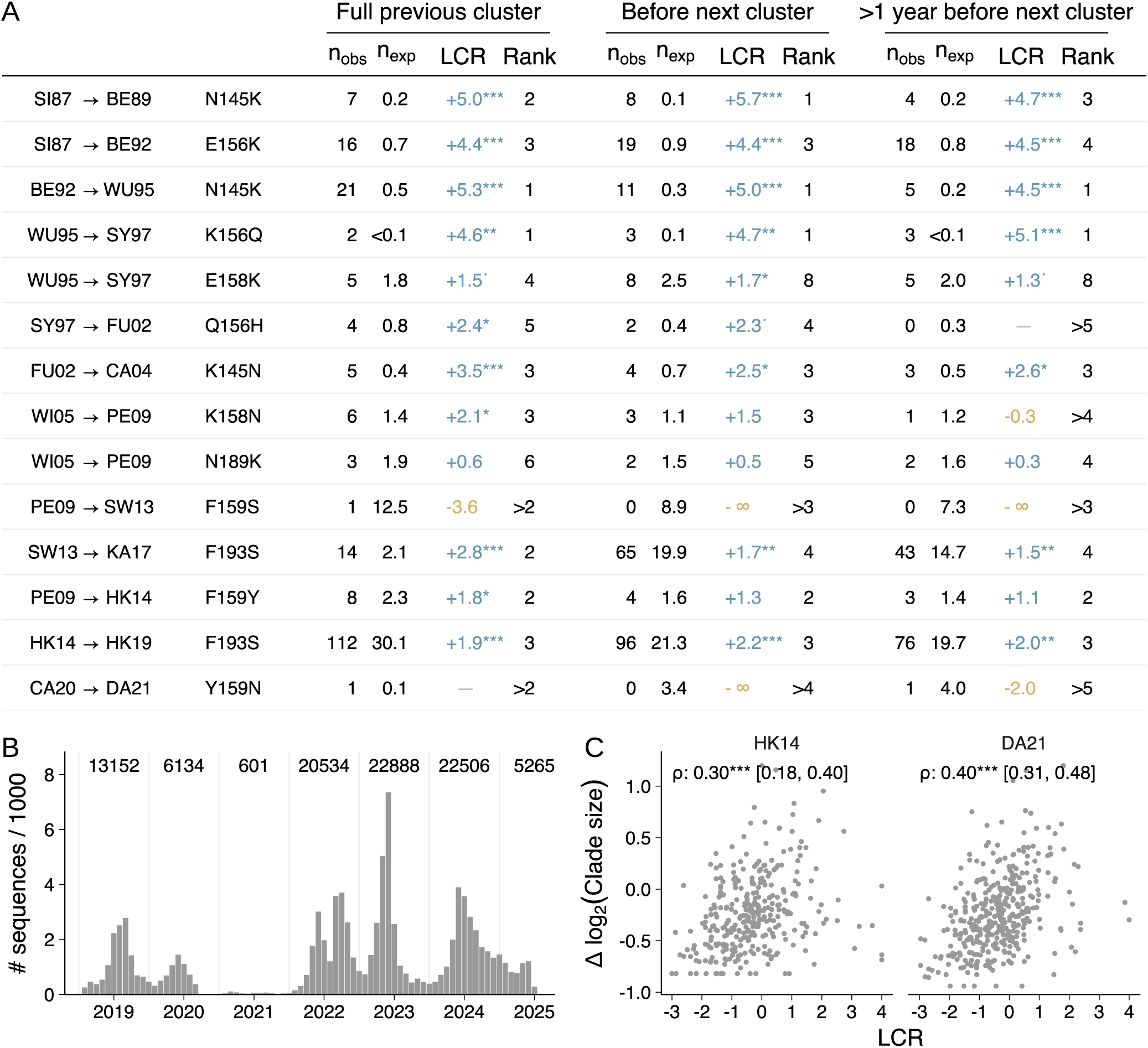
Antigenic cluster transition substitutions are typically identified by convergent evolution before the establishment of the next antigenic cluster. (**A**) Log convergence ratio (LCR) and rankings for cluster transition substitutions from 1987 to 2024. The “Full previous cluster” column shows measurements using all sequences in the ancestral cluster; the “Before next cluster” and “>1 year before next cluster” columns use all sequences detected 0-36 months or 12-48 months before the descendant cluster reached 5% frequency, respectively. n_obs_ and n_exp_ are the observed and neutral expected number of occurrences respectively. LCR values and ranks are only calculated for substitutions where either n_obs_ or n_exp_ exceed one (see Methods section *Calculating ranks* for details). Two cluster transitions are not shown: CA04 to WI05 (S193F), as S193F required two nucleotide changes, so an LCR cannot be estimated; and HK14 to CA20 (F193S), as F193S previously caused the transition from HK14 to HK19. See Fig. S10 for clusters predating 1987. (**B**) Number of sequences per year (June to May) in the full GISAID dataset after processing. (**C**) Correlation between LCR and average clade size relative to synonymous mutations, for substitutions in the HK14 and DA21 clusters with at least eight observed occurrences. Spearman’s rank correlation is shown with a 95% bootstrap confidence interval. ^.^ = p<0.1, * = p<0.05, ** = p<0.01, *** = p<0.001.

### Predictive sequences are placed reliably in the tree

We wanted to check that “early occurrences” of cluster transition substitutions are truly phylogenetically independent, and not incorrectly placed instances of a larger clade (for example, the next antigenic cluster) which falsely led us to infer additional convergence. To test this, we removed all early occurrences of antigenic cluster transition substitutions from each cluster, and identified all “near-optimal” placements of these occurrences (defined as those which increase the parsimony score by less than two compared to their optimal placement).

Across the cluster transition substitutions, an average of 95% of the early occurrences had all near-optimal placements within the ancestral cluster, indicating they were accurately placed in the original tree as an early occurrence. To check whether the few early occurrences with ambiguous placements substantially affect the prediction results, we calculated “minimized” counts of early occurrences for each cluster transition substitution, by discarding early occurrences which had any near-optimal placements outside the ancestral cluster, and combining those which shared near-optimal placements (see Methods section *Minimized counts of early occurrences*). Using these minimized counts, the rank of a cluster transition substitution never increased by more than one position, and the median rank remained 3^rd^—despite using non-minimized LCRs for the other substitutions in the rankings (Fig. S12). We therefore conclude that the predictive power of the LCR for major antigenic substitutions is not caused by misplacement of sequences in the tree.

### Convergent evolution identifies cluster transition substitutions more than one year before the establishment of the next antigenic cluster

Successive antigenic clusters typically cocirculate for a period (Fig. S13). For prospective prediction, positive selection for the antigenic cluster transition substitution must be detectable before the establishment of the new cluster. To test whether this is possible, we calculated LCRs using sequences detected before the descendant cluster reached 5% frequency (see Methods section *Constructing temporally early trees* and Fig. S14). For all 12 cluster transition substitutions with a positive LCR in the whole ancestral cluster, the LCR remained positive when calculated using only the temporally early sequences, with all 12 ranking in the top eight positions, and 11 of 12 ranking in the top five (Fig. 4A). We repeated the analysis using sequences collected more than one year before the establishment of the descendant cluster: the LCR remained positive and the ranking within the top eight positions for 10 of the 12 substitutions, with 9 of 12 ranking in the top four. Together, these results suggest that the LCR can identify likely future antigenic substitutions substantially in advance of their circulation.

### Cases where cluster transition substitutions are not predicted by log convergence ratios

There have been two cluster transition substitutions since 1987 that the LCR did not identify as being positively selected in the ancestral cluster.

One case, Y159N (CA20 to DA21), is likely due to an unusually small number of sequences being available from the ancestral cluster: the observed number of occurrences of Y159N (one) did in fact exceed its neutral expected number of occurrences (0.1)—but we do not rank substitutions with only one observed occurrence, because positive selection cannot be confidently inferred based on only a single occurrence, which may be only a single isolate. It is unlikely that so few sequences will be available for future clusters: the period of circulation for CA20 coincided with the SARS-CoV-2 pandemic, so was sparsely sequenced with ∼30-fold fewer sequences collected in 2020/21 than in any of the subsequent years (Fig. 4B). There were therefore only 909 CA20 sequences available, compared to at least 1400 for every other cluster since 1997. Y159N is also notable for being a potential example of epistasis: prior to Y159N’s noisy but positive LCR in CA20, and its fixation in DA21, it had an LCR of -2.5 in HK14 (the ancestor of CA20). This change is likely caused by an epistatic interaction with the Y195F substitution, which occurred at the start of the CA20 cluster, without which Y159N substantially reduces receptor binding strength (*61*).

The second case, F159S (PE09 to SW13), is the only cluster transition substitution under negative selection in its ancestral cluster, with an LCR of -3.6 in PE09 and -2.9 in PE09’s ancestor, WI05. This is unusual not only among cluster transition substitutions, but also among trunk and major branch substitutions in general: out of 92 substitutions with sufficient data to assess selection, only two others have an LCR below -1.8 among sequences collected in the previous two years (Fig. 2A). Notably, SW13 and its descendant KA17 reached a maximum frequency of only ∼37% (Fig. S14) before being outcompeted by HK14, another descendant of PE09 caused by a different substitution at position 159 (F159Y). Consistent with the fitness advantage of HK14 over SW13, the LCR identifies tyrosine (Y) as fitter than serine (S) at position 159 before, during, and after the circulation of SW13 and HK14: the LCR is positive for F159Y in PE09, and for S159Y in both SW13 and its descendant KA17 (Fig. S9). The negative LCR for F159S in PE09 is unlikely to be explained by epistasis: the LCR for S159Y in SW13 is even greater than that for F159Y in PE09, suggesting 159S is unfit even in the SW13 sequence context.

### Convergence measurements may identify unrealized adaptive paths for evolution

It is informative to consider the evolutionary fate of substitutions ranked above the antigenic cluster transition substitutions (“outranking substitutions”). 25% of outranking substitutions (10 of 40) are themselves cluster transition substitutions, either for another cluster transition from the current cluster, or for a cluster transition from the *next* antigenic cluster (Figs. S9 & S15). For example, N189K (which caused the WI05 to PE09 cluster transition) was ranked below F159Y in WI05, with F159Y later causing the PE09 to HK14 cluster transition (Fig. 3A). Such cluster transition substitutions may have been viable alternative paths for antigenic evolution from earlier clusters, before later being taken in nature (Fig. S16). In two further cases (S145N, EN72 to TX77; and Y159F, FU02 to CA04), the outranking substitution became fixed in the descendant cluster, but was not responsible for the antigenic effect at the cluster transition (*11*)—with positive selection potentially caused by an effect on a non-antigenic aspect of phenotype.

Among the remaining 70% of outranking substitutions (28 of 40) which do not get fixed, the LCR remains positive in the descendant cluster 75% of the time (21 of 28), suggesting they too may have been fit alternative paths for evolution, which could have resulted in antigenic change. Indeed, this is true of substitutions which have a positive LCR but which are ranked below the cluster transition substitution, of which there are two on average in each cluster.

There are some occasions where positive selection does not persist in descendant clusters: for example, F193Y is the highest ranked substitution in PE09, with an LCR of 2.4, but is negatively selected in HK14 with an LCR of -0.9 (Fig. S9), potentially due to changes in the selective environment or genetic context between the clusters.

### Average clade size correlates with convergence, but is less predictive of cluster transition substitutions

In addition to beneficial substitutions occurring more frequently on the tree than neutral or deleterious substitutions, their occurrences are expected to produce larger descendant clades—and indeed we find a moderate but statistically significant correlation between the LCR and average descendant clade size (r = 0.30 in HK14, r = 0.40 in DA21; both p < 0.001; Fig. 4C). To test whether clade size or convergence ratio is a better predictor, we checked whether cluster transition substitutions typically had larger average clade sizes than synonymous mutations. We found seven instances where the LCR for a cluster transition substitution was positive, suggesting positive selection, but the average clade size was smaller than that of synonymous mutations, suggesting negative selection—and zero instances of the opposite, where clade size suggests positive selection but the LCR is negative (Fig. S17). Consequently, we conclude that average clade size less accurately identifies future cluster transition substitutions than the LCR.

### Sampling rate partially determines which substitutions contribute to HA evolution

When calculating the LCR, we control for the >50x difference in mutation rate between nucleotide mutations (Fig. 1B). These mutation rate differences also affect the sampling rate of amino acid substitutions, with some substitutions having more opportunities to occur and potentially become fixed. Population genetic theory predicts that such differences in mutational supply can affect evolutionary outcomes (*62*–*65*). If only fitness, and not sampling rate, determined which substitutions become fixed, we would expect each nucleotide mutation to contribute a similar proportion of fixed substitutions, after accounting for the number of ways each mutation can cause an amino acid substitution, and disregarding correlations between the genetic code and properties of amino acids (*66*–*68*).

Indeed, we find that nucleotide mutations with higher neutral sampling rates cause a higher proportion of amino acid substitutions on the tree as a whole, on the trunk and major branches of the tree, and of antigenic cluster transition substitutions (Fig. 5A). The two lowest rate nucleotide mutations (CG and GC) have caused zero out of the 175 single nucleotide amino acid substitutions on the trunk and major branches, and therefore zero out of 21 single nucleotide cluster transition substitutions—while the six (out of 12) nucleotide mutations with the highest rate have caused 79% of trunk and major branch substitutions, and 67% of cluster transition substitutions. Differences in mutational supply therefore partially determine the identity of the substitutions which contribute to the long-term sequence evolution and antigenic evolution of the virus.

**Fig. 5.**
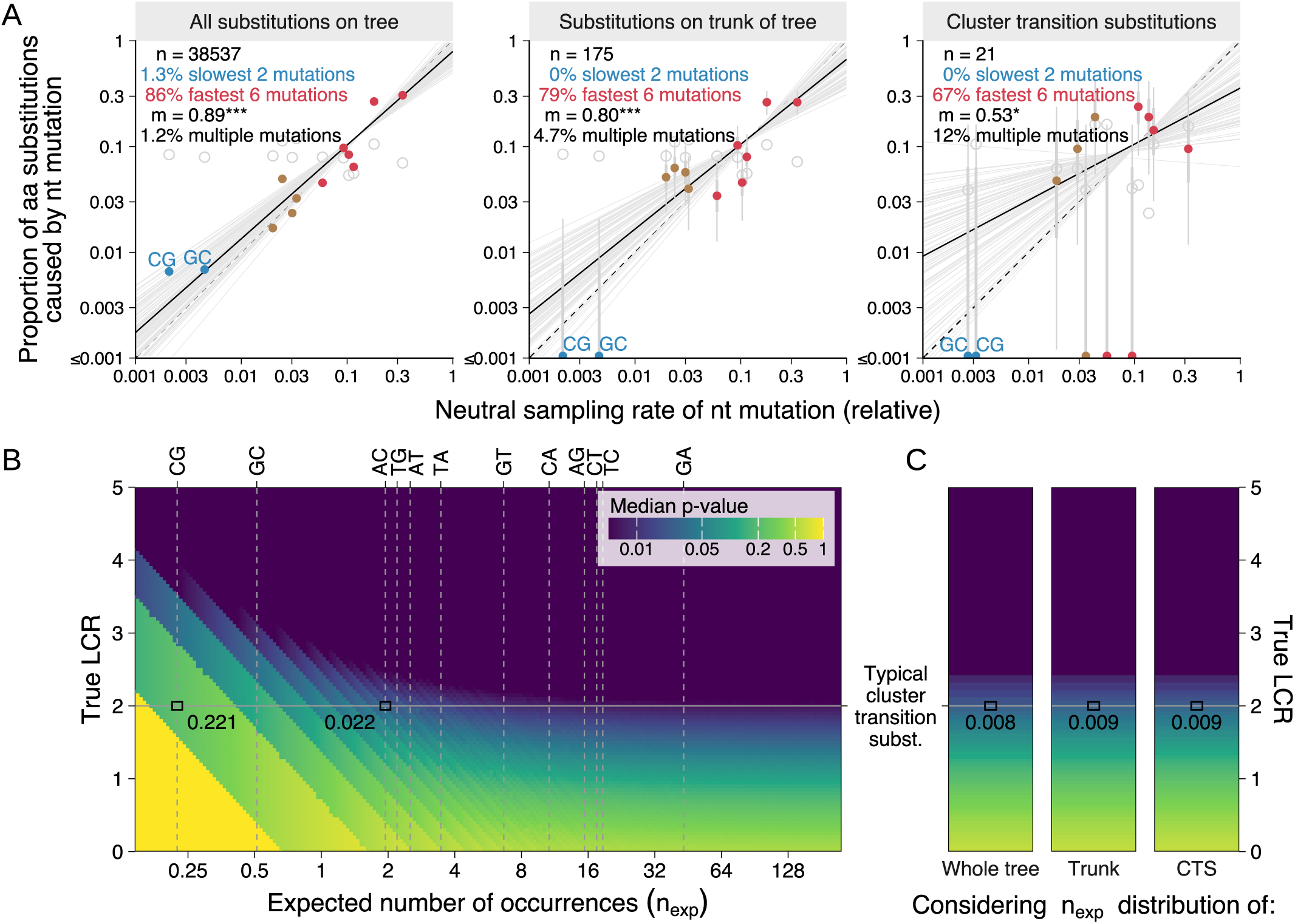
Nucleotide mutation rate affects evolutionary trajectory and the detectability of positive selection. (**A**) Nucleotide (nt) mutations with faster neutral sampling rates cause a disproportionately large fraction of observed amino acid (aa) substitutions. The proportion of single-nucleotide aa substitutions caused by each nt mutation is plotted against its neutral sampling rate (95% CI is thin, 50% CI is bold). Results are shown for aa substitutions occurring anywhere on the tree, those occurring on the trunk & major branches, and cluster transition substitutions (*n* indicates the number of aa substitutions in each group). The proportions of aa substitutions caused by the slowest two nt mutations (colored blue) and fastest six nt mutations (red) are given. The maximum-likelihood regression line is shown in black; 100 parameter samples are shown in grey; *m* is the regression line gradient (see Methods section *Regression of substitution proportions*). Hollow points show the expected proportions for each nt mutation ignoring mutation rate, based on the number of opportunities for each nt mutation to cause an aa substitution. The proportion of aa substitutions caused by multiple nt mutations, excluded from other analyses, is shown. (**B**) Heatmap showing how the detectability of positive selection (median empirical p-value; represented by coloring) varies based on the true log convergence ratio (LCR; y-axis) and the neutral expected number of occurrences (n_exp_; x-axis) for a substitution. Darker colors indicate lower p-values, and therefore stronger evidence for positive selection. Vertical dashed lines show n_exp_ for each nt mutation in 40,000 sequences. The horizontal line indicates an LCR value typical of cluster transition substitutions. Empirical p-values referenced in the text are highlighted. See Methods section *Simulating LCRs for a hypothetical cluster transition substitution* for details. (**C**) As **B**, except n_exp_ values are sampled from empirical distributions of observed substitutions (Fig. S19). Results therefore represent average detectability across substitutions in each of the three groups: substitutions observed anywhere on the tree, those observed on the trunk, and cluster transition substitutions.

This effect extends to amino acid substitutions requiring multiple nucleotide mutations in a codon, which are sampled at a lower rate than those requiring only one mutation. Such multi-nucleotide substitutions comprise 1.2% of all amino acid substitutions in the tree, 4.7% on the trunk and major branches, and 12% of antigenic cluster transition substitutions (Fig. 5A)—substantially lower than the ∼70% expected if sampling rate had no effect (∼13 of the 19 amino acid substitutions at a position require >1 nucleotide mutation on average for naturally occurring H3 HA sequences).

### Current sequencing levels are sufficient to estimate the effect of most substitutions likely to contribute to HA evolution

We investigated how statistical uncertainty in LCR estimates affects its ability to identify positive selection for an amino acid substitution. Two key quantities determine this: first, the magnitude of positive selection for the substitution; and second, the neutral expected number of occurrences of the substitution, with larger values yielding more precise LCR estimates. Fig. 5B shows that stronger positive selection and larger neutral expected occurrence counts both result in greater statistical significance (lower p-value) for positive selection.

The expected number of occurrences of an amino acid substitution depends crucially on the nucleotide mutations which can cause it: in Fig. 5B, each vertical dashed line shows the neutral expected number of occurrences for one nucleotide mutation in an exemplar cluster containing 40,000 sequences, representing ∼2 years of circulation at current sequencing levels. If a hypothetical cluster transition substitution were caused by the lowest rate nucleotide mutation, CG, it would have a neutral expected number of occurrences of 0.22 in this exemplar cluster. Even if it were under strong positive selection, with a “true” LCR of 2.0 (typical of cluster transition substitutions), the median p-value would be 0.221 (Fig. 5B), and 41% of the time the substitution would not be observed even once in the 40,000 sequence cluster (Fig. S18A).

However, while ∼12% of amino acid substitutions accessible by a single nucleotide change are produced only by a CG or GC mutation (the two lowest rate mutations) (Fig. S19), these mutations have caused none of the 175 trunk and major branch substitutions or the 21 cluster transition substitutions since 1968 (Fig. 5A). Low confidence estimates of the LCR for substitutions caused by CG and GC are therefore unlikely to be problematic for predicting the evolution of the virus. A substitution caused by the third lowest rate nucleotide mutation, AC, would have a neutral expected number of occurrences of 1.95: for the hypothetical strongly selected cluster transition substitution with a true LCR of 2.0, the median p-value would be 0.022 (Fig. 5B), and the measured LCR would be positive 97% of the time in the 40,000 sequence cluster (Fig. S18A). If we consider the full set of substitutions which have historically occurred on the trunk & major branches, and the nucleotide mutations which could have caused them, the measured LCR for the hypothetical strongly selected cluster transition substitution would be positive 98% of the time (Fig. S18B), with a median p-value of 0.009 (Fig. 5C)—with similar results obtained when considering substitutions which occurred anywhere on the tree or considering cluster transition substitutions.

The temporal resolution of LCR estimates is likely to be important for prediction, because the substitutions under selection may change after a cluster transition or over time within a cluster. While it takes ∼2 years to achieve a neutral expected number of occurrences above 1.95 for substitutions caused by the 10 highest rate nucleotide mutations, this is achieved in ∼7 months for the 79% of trunk and major branch substitutions caused by the six highest rate nucleotide mutations. For these substitutions, convergence signals are likely to be detectable over time windows shorter than the two years considered here, with this time scale decreasing further if increases in sequencing rate continue.

However, the benefit of increasing sequencing rate is limited in two ways. First, increasing sequencing intensity gives diminishing returns on the neutral expected number of occurrences, because newly sampled sequences are on average increasingly closely related to previously collected sequences. We estimate that to double the expected number of occurrences, or halve the length of time required to achieve a particular expected number of occurrences, sequencing intensity must be increased by ∼2.7x (see Methods section *Modelling the effect of sequencing intensity on n_exp_* and Figs. S20 & S21).

Second, as sequencing rate increases, the magnitude of LCR estimates decreases (i.e. they get closer to zero). This is because lineages are sampled sooner after they arise, meaning there is less time for selection to affect the frequency of a lineage and therefore the probability it is sampled before going extinct. This is analogous to the sensitivity of dN/dS to the time since divergence for within-population samples (*69*, *70*).

In practice, we find that this effect is relatively small: an order of magnitude change in sequencing rate from current levels causes a ∼20% change in the LCR (Fig. S22), meaning sequencing rate changes are unlikely to change the interpretation of the LCR in the short to medium term.

### Substitutions at positions 145, 158, and 189 are under positive selection in recently circulating viruses

Fig. 6 shows cluster-wide and year-by-year convergence ratios for the DA21 antigenic cluster, which has circulated continuously since its emergence from the CA20 cluster in 2021 through to 2025. Six substitutions at the key antigenic positions have a positive LCR in sequences since April 2024: in order of their LCR, S145N, N158H, K189R, N158K, S145R, and N159S (Fig. 6A). Notably, the LCR for these substitutions is typically higher in more recent years, potentially reflecting increasing selection for antigenic escape as population immunity to DA21-like viruses increases. For the four substitutions with a sufficient number of occurrences (i.e. excluding N158H and S145R), we calculated year-by-year average clade sizes compared to synonymous mutations. Among these, S145N, N158K, and K189R showed a pattern of increasing average clade size over time in addition to their increasing LCR, while N159S did not (Fig. 6B). Notably, 158K and 189R both circulated at high frequency in late 2025 in clades designated J.2.3, J.2.4, and K (Fig. S23), the emergence and spread of which are considered in more detail below.

**Fig. 6.**
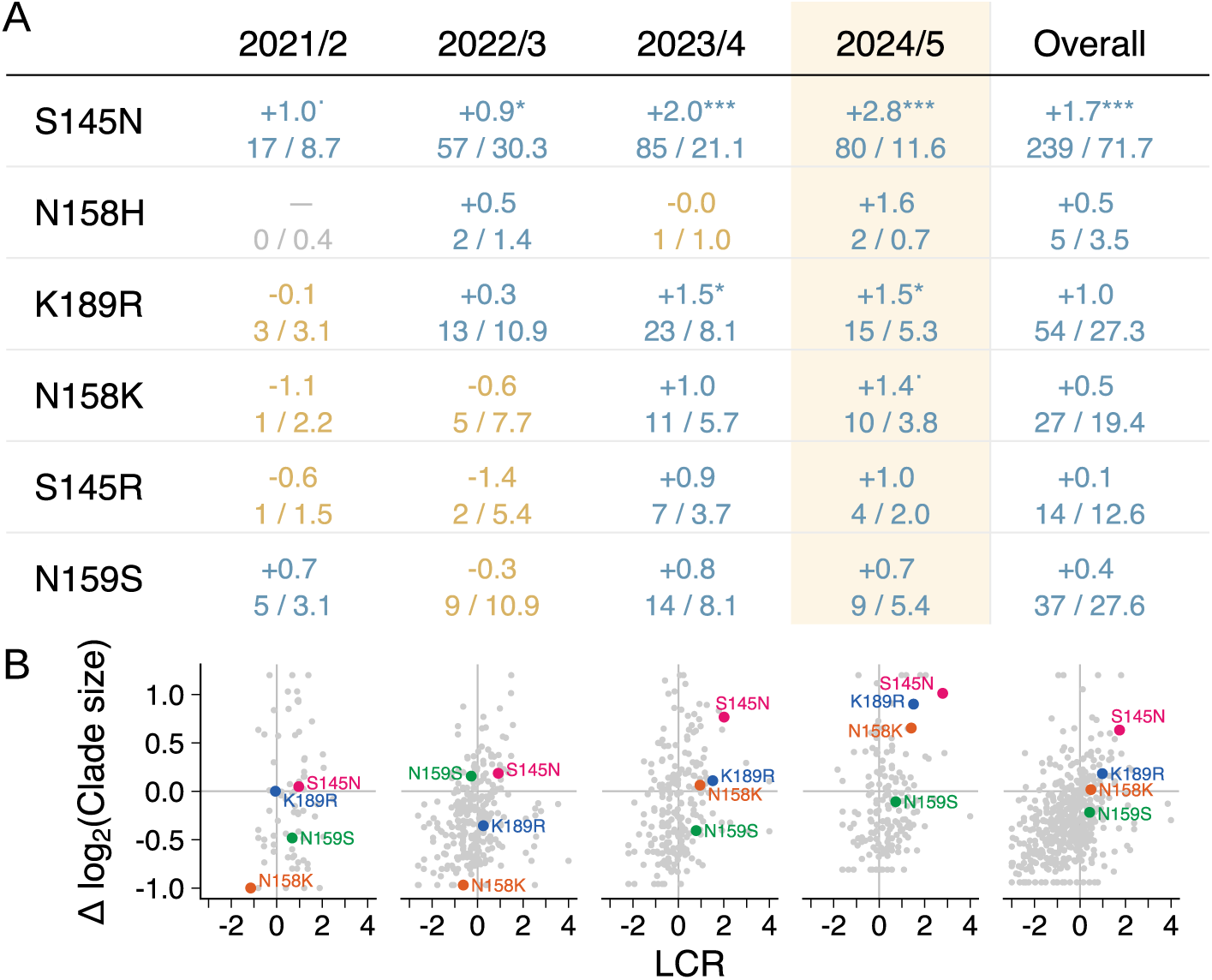
S145N, N158K, and K189R substitutions are highly convergent and produce large clades in recently circulating viruses. Year-by-year and cluster-wide log convergence ratios (LCR) (**A**) and average clade sizes relative to synonymous mutations (**B**) for substitutions with a positive LCR in 2024/25 in DA21. In **A**, the LCR is shown in large font, and the observed and expected number of occurrences in smaller font below. Light grey points in **B** show data for other substitutions in HA1. Years are April to March. ^.^ = p<0.1, * = p<0.05, ** = p<0.01, *** = p<0.001.

## Discussion

We systematically measured convergent evolution in a global phylogeny of influenza H3N2 viruses. We find that substitutions occurring on the trunk or leading to antigenic clusters were typically under positive selection, in contrast with substitutions on other branches of the tree. We find that the next antigenic cluster transition is nearly always caused by one of few substitutions under positive selection at the key antigenic positions in the currently circulating viruses—and further, that these signals were typically present at least one year before the establishment of the next antigenic cluster, making them highly informative for prospective prediction of the antigenic evolution of the virus.

Multiple aspects of this work corroborate and build upon previous findings at a more fine-grained level. Previous work has shown that adaptive substitutions often occur multiple times in the evolution of influenza H3N2 viruses (*15*–*17*); we show that systematic measurement of these convergence signals for all substitutions at antigenic positions can be used prospectively to predict antigenic evolution.

The log convergence ratio measured in this study is conceptually related to the dN/dS ratio, long used in evolutionary biology to infer the nature of selection. Existing dN/dS methods typically estimate selection at the level of amino acid sites (*20*, *22*, *24*, *25*); the log convergence ratio additionally identifies selection for particular amino acids at sites. This is preferable to the approach of first identifying sites under positive selection, before checking which substitutions are most common at those positions. First, because of mutational biases, the most frequent substitution at a position may not be the most strongly selected. Second, if different substitutions at a site are under a mixture of positive and negative selection, a site-level method may not identify the site as being under positive selection in the first place.

Our work is also more finely resolved in the time scale over which we measure selection. dN/dS has often been calculated using sequences covering longer periods of influenza virus’ evolutionary history, aiming to identify long-term temporal and structural patterns in selection (*39*, *71*)—including a previous use of dN/dS for evolutionary prediction, which produced a list of the most critical positions for monitoring by identifying sites with positive dN/dS in a phylogeny containing sequences collected over the preceding ∼15 years (*8*). By contrast, we estimate recent selection acting on each substitution over a time scale of a few years, to identify those that are beneficial to viral fitness in the current sequence context and selective environment.

Our estimates of selection, made for individual substitutions over short time windows, corroborate the evolutionary patterns found in previous work using either dN/dS measurements aggregated over long periods of evolution and/or groups of multiple amino acid positions, or using frequency-based methods: we find that the trunk lineage is characterized by positive selection, particularly for mutations in antigenic sites and near to the receptor binding site (*9*, *15*, *21*, *39*, *71*, *72*); that mutations on side branches of the tree are often deleterious (*9*, *37*, *38*); and that deleterious substitutions can reach high frequency or fix, but do so only rarely (*40*, *41*).

There are three primary limitations to the use of convergence to measure selection. First, it relies on global surveillance and sequencing of a substantial number of viruses. We find that current, 2026, surveillance levels are sufficient to detect the direction of selection at least 97% of the time for a typical antigenic cluster transition substitution with two years of sequence data, if it is caused by the top 10 (of 12) highest rate nucleotide mutations. To achieve this for the two lowest rate nucleotide mutations would require a further ∼25-fold increase in sequencing rate. However, for practical purposes this is unlikely to be an issue, with the low rate of these nucleotide mutations meaning they have never caused an antigenic cluster transition substitution, or indeed any other substitution on the trunk or leading towards an antigenic cluster.

Second, this requirement for a large quantity of sequences means they must be aggregated—across geography, time, and genetic background—with the convergence measurement representing a weighted (by the quantity of available sequences) average over these aspects of context. If a key aspect of evolution is occurring in an under-surveilled (or non-surveilled) region or population, or a key substitution is fit in some sequence contexts but not others, the signal will be underrepresented in the aggregated sequence dataset—though this has not prevented prediction in historical cases, as the LCR was positive for all except one cluster transition substitution since 1987. The flip side is that convergence ratios can be used to identify instances of epistasis and spatiotemporal variation in selection, by partitioning sequences based on geography, time, and genetic background.

The third limitation is that approximately 10% of cluster transition substitutions required multiple mutations in the same codon. Because we calculate the neutral expected number of occurrences using the rate of synonymous single nucleotide changes, we do not calculate a log convergence ratio for multi-nucleotide substitutions. While theoretically possible to do so, sequencing rate would need to be increased by roughly two orders of magnitude to achieve similar resolution to that which is currently possible for single nucleotide substitutions, given that only ∼1% of observed amino acid substitutions are multi-nucleotide substitutions.

Measurements of convergence can form an integral part of a protocol for prospectively predicting the substitutions most likely to drive antigenic novelty of future influenza virus strains, which can inform the design of vaccine antigens which are protective against upcoming antigenic variation. In such a protocol, a shortlist of antigenic substitutions under positive selection in nature could be produced by measuring convergence: historically, there have been an average of 5.3 such substitutions at key antigenic positions in each antigenic cluster, and the next antigenic cluster transition substitution has nearly always been one of them.

Indeed, we have produced such shortlists for WHO Vaccine Composition Meetings (Data S1-S3) since the Vaccine Composition Meeting for the Southern Hemisphere in September 2024. At that time, N158K and K189R had the strongest convergence signals among low frequency substitutions, circulating at 0.8% and 1.6% frequency respectively in the preceding 3 months (Fig. S23, Data S1). We incrementally updated the substitution shortlist for the two teleconferences prior to the Northern Hemisphere VCM (in December 2024 and January 2025), and tracked the ongoing emergence of new lineages containing 158K or 189R (Data S2 & S3). This led to the recognition in January 2025 of the importance of a clade of 12 “double mutant” viruses containing both substitutions, resulting in rapid epidemiological and experimental characterization which confirmed their substantial antigenic divergence ahead of the Northern Hemisphere VCM (*73*, *74*). The clade subsequently spread globally, reaching peak frequencies of ∼95% in South America and ∼40% in North America as of December 2025. Meanwhile, another clade containing 189R (subsequently designated clade K) began to spread rapidly in July 2025, leading to an updated WHO vaccine recommendation in September 2025 to a virus carrying 189R (*75*).

Clade K viruses also contain 158D, obtained via an N158D substitution, which has an LCR of -3.6 among sequences collected between April 2022 and April 2025. If this clade reaches fixation, N158D would join only three other trunk or major branch substitutions with an LCR below -1.8: first, F159S, responsible for the transition to the SW13 cluster, which went extinct without becoming the majority cluster; second, D225N, which occurred in CA04 and reduced receptor binding avidity (*76*), and reverted to 225D after a period of strong positive selection for the reversion in the WI05 and PE09 clusters (LCR of +3.3 in both clusters); and third, G186D, which occurred in DA21 and which severely reduces receptor avidity in the absence of 190N, which was obtained concomitantly with 186D in nature (*77*). Notably, an I160K substitution co-occurred with N158D on the phylogenetic branch ancestral to the clade K viruses. The rarity of a substitution with such a low LCR measurement reaching high frequency, and its proximity to the I160K substitution, suggests a possible epistatic interaction between the N158D and I160K substitutions.

We envisage two key refinements to a convergence-based prediction protocol for antigenic substitutions. First, predictions will likely be improved by accounting for the sampling rate of amino acid substitutions, by deprioritizing substitutions which are beneficial but unlikely to arise. Consistent with theoretical expectations (*62*–*65*), we find that the >50x differences in sampling rate in H3N2 influenza substantially affect the identity of substitutions which are evolutionarily successful and cause antigenic change. This corroborates observations in experimentally and naturally evolving populations of other species, where mutational supply has been shown to affect which substitutions contribute to adaptive evolution (*78*–*82*).

Second, measurements of convergent evolution reflect the total effect of a substitution on viral fitness in nature, which is determined by the interaction between many components of phenotype. The approach therefore avoids a universal difficulty associated with characterizing the effect of substitutions in the laboratory—that such approaches necessarily test an incomplete set of phenotypic characteristics, with assay conditions only partially reflecting those faced by the virus during circulation, and no clear way to combine measurements of different aspects of phenotype into an estimate of total fitness.

However, for selecting and designing vaccine antigens, it is important to understand whether each predicted substitution would cause antigenic escape. Laboratory-based phenotyping of the shortlisted substitutions is therefore essential, allowing the convergence-based measurement of total fitness to be decomposed into individual phenotypic components, including an estimate of the antigenic effect size for each substitution. In this study, our rankings include all substitutions at seven historically key antigenic positions identified in Koel et al. 2013 (*11*). However, substitutions at these positions often do not result in antigenic change (*11*, *54*), while substitutions at other positions can contribute to antigenic change, albeit typically with smaller effect sizes (*54*). Phenotypic characterization of positively selected substitutions across a broader set of positions—including potentially the entire HA (Fig. S24) and NA proteins—would refine the rankings by the exclusion of non-antigenic substitutions at positions from Koel et al. 2013, and inclusion of antigenic substitutions at other positions.

Beyond its value for prospectively evaluating the threat posed by strains carrying each substitution, joint information on the circulation probability and antigenicity of each substitution could be produced and used retrospectively to help build an understanding of how different aspects of phenotype contribute to the fitness of viral strains.

## Acknowledgments

We gratefully acknowledge all data contributors, i.e., the Authors and their Originating laboratories responsible for obtaining the specimens, and their Submitting laboratories for generating the genetic sequence and metadata and sharing via the GISAID Initiative, on which this research is based.

## Author contributions

Conceptualization: SAT, DJS, RAMF

Methodology: SAT, DJS

Software: SAT, DJP

Investigation: SAT, DJP, DJS

Resources: DJS

Data curation: SAT

Writing - Original Draft: SAT, DJS

Writing - Review and Editing: All

Visualization: SAT, DJS Supervision: DJS

## Data and materials availability

Code and data can be accessed at https://github.com/acorg/predicting_influenza_using_convergence_ms_public

## Supplementary Materials

### Materials and Methods

#### Sequence data

For the main analysis, all HA nucleotide sequences for human H3N2 viruses were downloaded from GISAID up to 28^th^ January 2025. Sequences were aligned to A/Alaska/232/2015 (EPI_ISL_202884) using mafft (*83*) with options --6merpair --keeplength --addfragments, and trimmed to HA1 (amino acid positions 1 to 328 in H3 numbering). Sequences were discarded if they contained >10% gaps or Ns between nucleotide positions 100 and 900 (older sequences often have incomplete coverage at the start and end of HA1), or if there were >100 mismatches to a consensus of all sequences isolated in the same year. Egg passaged sequences were filtered out by removing those with passage information matching the regular expression “AM[1–9]|E[1–7]|AMNIOTIC|EGG|EX|AM_[1–9]”. Isolate names were standardized by (a) removing the ‘A/’ prefix when present; (b) replacing all characters except alphanumeric characters and ‘_’, ‘-‘, ‘.’, ‘|’, or ‘/’ in the isolate name with ‘_’; (c) putting the year in four-digit format; (d) discarding all parts of the isolate name following the year; (e) capitalizing the isolate name.

When multiple sequences were present with the same isolate name (after standardization), the sequences were deduplicated by taking the consensus. In these cases, the sequence collection date was set to the earliest collection date excluding YYYY-01-01 dates (except if only YYYY-01-01 dates were available). An additional alignment containing sequences up to 16^th^ October 2025 was produced for Fig. S23, by additionally downloading sequences from GISAID on 16^th^ October 2025, selecting those with submission dates since 1^st^ January 2025, and processing as above. A table of GISAID accession IDs used in this study is available at: https://github.com/acorg/predicting_influenza_using_convergence_ms_public/blob/main/make_alignment/results/gisaid_acknowledgements.csv.

#### Tree construction

A starting tree containing a temporally evenly distributed subset of the sequences was built using IQTREE 2 (*27*). Two whole-alignment trees were then produced by expanding the starting tree with the remaining sequences either using CMAPLE (*29*) or using UShER (*30*) and matOptimize (*34*). To produce the starting tree, a reduced alignment which was approximately evenly sampled over time was produced by randomly selecting up to 50 sequences per year. The starting tree was constructed from this alignment with IQTREE 2 using the GTR substitution model. To produce the CMAPLE tree, the remaining sequences in the full alignment were iteratively added to the starting tree with CMAPLE using the GTR substitution model with the --search option set to “NORMAL”, producing intermediate trees with up to 200, 300, 400, … sequences per year, until all sequences were included. To produce the UShER+matOptimize tree, the remaining sequences were added using UShER with the --write-uncondensed-final-tree option, and further optimized using matOptimize with the options --min-improvement 0.0000000000001 --drift_iteration 100 --max-iterations 1000000000 --max-hours 10. Both trees were rooted using A/HONG KONG/1-10-MA21-2/1968 (EPI_ISL_25014) as the outgroup. Sequences descending from long branches were manually inspected and removed from the tree. Names and GISAID accession IDs for the 161 removed sequences are available at https://github.com/acorg/predicting_influenza_using_convergence_ms_public/blob/main/make_figures/data/sequence_removals.txt.

Nucleotide mutations on branches of the phylogenetic trees were estimated by producing a mutation-annotated tree using UShER, and extracted from the mutation-annotated tree using the matUtils program included with UShER. The mutation annotations were used to infer nucleotide sequences at the internal branches.

#### Antigenic clusters and causative cluster transition substitutions

We aggregated results from numerous sources to assign antigenic clusters and causative antigenic cluster transition substitutions. To assign antigenic clusters, for evolution up to approximately 2002 we used Smith et al. 2004 (*1*); for evolution between approximately 2002 and 2012 we used Russell et al. 2008 (*48*), Bedford et al. 2014 (*49*), and Fonville et al. 2014 (*50*); and for subsequent evolution we used Chambers et al. 2015 (*51*), Li et al. 2016 (*52*), and HI data from reports produced by the Francis Crick Institute for WHO Vaccine Composition Meetings (*60*). For the assignment of causative cluster transition substitutions, we used Koel et al. 2013 (*11*) for clusters up to FU02; and for subsequent clusters, we used (primarily) Chambers et al. 2015 (*51*), Neher et al. 2016 (*54*), Pattinson 2019 (*53*), and Jorquera et al. 2019 (*59*). A detailed description of this data is available in Supplementary text section *Assignment of antigenic clusters and causative cluster transition substitutions*.

#### Assignment of clusters on phylogenetic tree

To identify sections of the phylogenetic tree corresponding to each antigenic cluster, we first identified cluster ancestor branches by choosing a known representative virus from the cluster, and finding the nearest ancestral branch annotated with the corresponding cluster transition substitution for that cluster. Each branch of the tree was assigned to a cluster by identifying the nearest ancestral branch which was a cluster ancestor branch. For clusters with multiple causative cluster transition substitutions which occurred on different branches, the cluster ancestor was defined as the branch where the final cluster transition substitution was obtained. The earlier branch(es) where other cluster transition substitution(s) occurred were also marked so appropriate sections of the tree could be removed for prediction tests of those cluster transition substitutions.

#### Defining the trunk and major branches

The “trunk and major branches” of the tree were defined to include all branches on the path between the root of the tree and the ancestral branch of the J clade (*84*), or on the path between the root and any cluster ancestor branch. There are some instances where a cluster went extinct without producing any descendant clusters (e.g. BE89). We include branches ancestral to these clusters because they represent major groups of antigenically divergent viruses which circulated widely enough that a vaccine strain was selected from them.

#### Calculating the LCR

To calculate the LCR for an amino acid substitution in a set of branches of the phylogenetic tree we calculate:

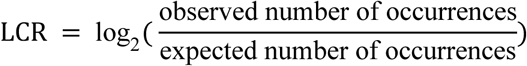

The observed number of occurrences is the number of branches annotated with a nucleotide mutation which causes a particular amino acid substitution. Because the same nucleotide mutation can result in different amino acid substitutions in different codon contexts, it is necessary to count occurrences of each nucleotide mutation separately for each codon context. The calculation of the expected number of occurrences for each [nucleotide mutation, parental codon] pair is more complex, and is described in the next section.

LCR estimates for amino acid substitutions are produced by summing the observed and expected numbers of occurrences for [nucleotide mutation, parental codon] pairs which result in that amino acid substitution.

In Fig. 2B, we include low fitness substitutions with zero observed occurrences (which would have an LCR of negative infinity using the definition above) by adding an arbitrary pseudo-count of 0.5 to the observed and expected occurrence counts.

#### Calculating the expected number of occurrences

The neutral expected number of occurrences for a particular nucleotide mutation (e.g. C100T) is calculated based on the observed rate of synonymous mutation of the same type (e.g. CT) at four-fold synonymous positions (where the same amino acid is encoded regardless of which of the four nucleotides is present).

We identify nucleotide positions which are four-fold synonymous at more than 95% of the branches of the tree, and where more than 90% of sequences share the same nucleotide (the “parental” nucleotide). At these positions, we count the number of occurrences of each of the three nucleotide mutations away from the parental nucleotide. To account for mutation rate variation, which is needed for significance testing, we model the distribution of these mutation counts (rather than simply averaging them) by fitting a hierarchical model of the form:

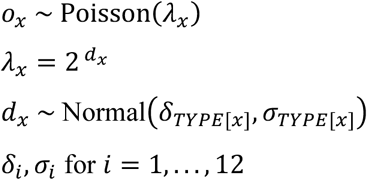

Where:

*o*_*x*_ is the observed number of occurrences of nucleotide mutation *x*
TYPE[*x*] ∈ {1, . . . , 12} is the mutation type (C to G, C to T, etc.) of the nucleotide mutation *x*
*δ*_*i*_, *σ*_*i*_ are the mean and standard deviation of the log_2_-rate distribution for mutation type *i*

We estimate each *δ*_*i*_ and *σ*_*i*_ by maximum likelihood with the ‘optim’ function in R, using the L-BFGS-B method, to obtain *δ̂*_*i*_ and *σ̂*_*i*_. We use the median as the point estimate for the expected number of occurrences for mutations of type *i*, *n*^*i*^_*exp*_ = 2^*δ̂_*i*_*^.

We require *n*^*i*_exp_^ values for many distinct sets of branches: first, we calculate LCR estimates for partitions of the tree which correspond to antigenic clusters and/or time intervals; and second, within each partition, we require as many *n*^*i*_exp_^ values as there are [nucleotide mutation, parental codon] pairs in the partition (because a mutation may only be accessible at a subset of the branches in the partition, and the resultant amino acid substitution depends on the codon context). However, it is computationally expensive to fit the model described above, and estimates may be noisy when calculated for sets of branches containing few occurrences of mutations. Rather than fitting the model for each branch set of interest, we fit the model once using the whole tree. We then scale the *n*^*i*_exp_^ values according to the size (defined below) of the branch set, and use the same *σ*_*i*_ as estimated for the whole tree. This has the advantage that the size of a branch set can be computed quickly by counting mutations, and can be defined using all mutational occurrences at four-fold synonymous positions, rather than the occurrences being partitioned between the 12 mutation types when fitting the synonymous mutation rate model.

The size of a branch set *t*, *S*_*t*_, is the total number of synonymous mutations per four-fold synonymous site, adjusted to account for differences between branch sets in the number of four-fold synonymous sites and in the number of four-fold synonymous positions with each parental nucleotide. Considering positions which are four-fold synonymous at greater than 95% of the branches in the set, and with greater than 90% identity for a particular parental nucleotide:

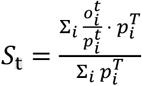

Where:

*t* is the branch set of interest
*T* is the branch set to which the *δ*_*i*_ and *σ*_*i*_ were fit (typically the entire phylogenetic tree)
*i* indexes the 12 nucleotide mutation types
*o*^*t*^_*i*_ is the total number of observed occurrences of all nucleotide mutations of type *i* at four-fold synonymous positions in branch set *t*
*p*^*t*^_*i*_ is the number of four-fold synonymous positions with parental nucleotide matching that of mutation type *i* in a branch set *t*

To obtain the expected number of occurrences of a nucleotide mutation of type *i* in a branch set *t*, we scale the *n*^*i*_exp_^ value obtained for the whole tree *T* according to the relative size of the branch set *t* compared to the whole tree *T*:

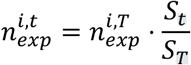

To summarize, we first estimate *n*^*i*,*T*^_*exp*_ for each mutation type using the full phylogenetic tree *T*. Then, for a given set of branches *t*, we calculate the size, *S*_*t*_. We use *S*_*t*_ to calculate an expected number of occurrences by scaling *n*^*i*,*T*^_*exp*_ by the relative sizes of the branch set and the full tree, 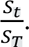

To obtain the expected number of occurrences of an amino acid substitution in a partition of the tree, we sum all of the *n*^*i*,*T*^_*exp*_ values for [nucleotide mutation, parental codon] pairs which result in the substitution (obtained by considering the branch sets *t* within the partition where each [nucleotide mutation, parental codon] pair can occur). To calculate the LCR, we compare this sum against the sum of the observed numbers of occurrences of those pairs in the branch sets.

LCR values are computed and reported per antigenic cluster, except for the “Trunk and major branches” panel of Fig. 2A, where LCR is calculated using the tree section corresponding to the 2 years of circulation prior to each trunk or major branch substitution. This is to ensure sufficient data is available for substitutions which become fixed shortly after the emergence of a new antigenic cluster.

#### Calculating empirical p-values

The above procedure produces a point estimate of the expected number of occurrences, against which we compare the observed count. We assess statistical significance by generating a null distribution as follows. For each [nucleotide mutation, parental codon] pair, we generate 10^4^ simulated occurrence counts by sampling *d*_*i*_^*t*^ ∼ Normal(log_2_ *n*^*i,t*^_*exp*_, σ_*i*_), and then *n*^*sim*^_*obs*_ ∼ Poisson (2*d^t^_i_*). For each amino acid substitution, we sum the *n*_*obs*_^*sim*^ values of [nucleotide mutation, parental codon] pairs causing the substitution (in the same way as the *n*^*i*,*T*^_*exp*_ values and *n*_*obs*_), to produce a null distribution containing 10^4^ counts. To calculate an empirical p-value for the alternative hypothesis that a substitution is subject to positive selection, we calculate the proportion of simulated occurrence counts which are greater than or equal to the real observed count.

We tested that these values were well-calibrated by checking that the distribution of empirical p-values for synonymous mutations was approximately uniform, using trees containing sequences from approximately 2021-2025, 2014-2020, and 1987-2008 (Fig. 1D). However, rather than fitting the model for *δ*_*i*_ and *σ*_*i*_ to each tree, we fit it only once to the 2021-2025 tree to obtain *δ*_*i*_^2021−2025^ (from which we obtain *n*_*exp*_^*i*,2021−2025^) and *σ*_*i*_^2021−2025^. We then scale the *n*_*exp*_^*i*,2021−2025^ according to the ratio of the tree sizes, and use the standard deviation for each mutation type directly. For example, for the 1987-2008 tree: 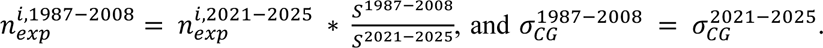 calculated empirical p-values for synonymous mutations in each tree, and show mutations with at least five neutral expected occurrences in Fig. 1D.

#### Testing the Poisson model of sampling variation

To test whether sampling variation in the expected number of occurrences is well described by a Poisson process, we took the subtree representing the DA21 antigenic cluster, and performed three independent splits of this tree, each into a pair of two approximately evenly sized subtrees. For each pair, we identified positions which were four-fold synonymous in both of the DA21 subtrees, and in a third subtree representing the HK14 antigenic cluster. We counted the number of occurrences of each mutation at these positions in each of the three subtrees (for a mutation *x*, *n*_*x*_^*DA*21−*A*^, *n*_*x*_^*DA*21−*B*^, and *n*_*x*_^*HK*14^). For each mutation, we used the observed count in HK14 as a baseline and scaled it to obtain expected counts for the DA21 subtrees: 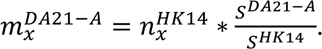 For each mutation, we then sample 100 values from each of Poisson(*m*_*x*_^*DA*21−*A*^) and Poisson(*m*_*x*_^*DA*21−*B*^). The lower panels of Fig. S2A show one of the sampled values for each mutation in each subtree. The frequency polygon in Fig. S2B is calculated using all 100 sampled values for each mutation in each subtree. Variation in the ratio in observed counts, 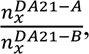 was similar to that obtained using Poisson-simulated values, indicating the Poisson model adequately captures sampling variation.

#### Minimized counts of early occurrences

To determine whether “early occurrences” of cluster transition substitutions in ancestral clusters were placed reliably, we estimated the smallest number of branches which could reasonably accommodate all of the occurrences. First, we identified and removed all branches (and descendants of branches) annotated with such occurrences, producing a “reduced tree”. We then determined the set of plausible placements in the tree for each early occurrence, by taking the first descendant sequence from each early occurrence and measuring the parsimony cost of placing it at each branch of the reduced tree using UShER with the option ‘--write-parsimony-scores-per-nodè. We then identified a set of near-optimal placements for each occurrence, defined as the branches where the occurrence could be added at a parsimony cost within one of the optimal value.

We used a greedy algorithm to estimate the smallest set of branches such that every early occurrence had at least one near-optimal placement within the set. First, we initialize an empty “minimized branch set”. At each step, we identify which branch is within the set of near-optimal placements for the largest number of early occurrences (excluding early occurrences which already have a near-optimal placement in the minimized branch set). We add this branch to the minimized branch set, and repeat until every early occurrence has a near-optimal placement in the minimized branch set.

#### Constructing temporally early trees

To construct trees representing the information available prior to the establishment of a new antigenic cluster for Fig. 4A, we constructed “temporally early” trees for each cluster transition. For each cluster, we produced a tree in which sequences had been removed if they had collection dates on or after the “establishment date” of the descendant cluster, or if they descended from the cluster-ancestor branch. The establishment date was set as the beginning of the first sequence of two year-quarters (if <30% of collection dates are only specified to the nearest year) or of two years (if ≥30% of collection dates are only specified to the nearest year) where the descendant cluster was at ≥5% frequency (Fig. S14). Collection dates where only the year is known are often recorded as YYYY-01-01. We conservatively reassign these to YYYY-12-31 when identifying sequences to remove, to avoid overestimating how early a convergence signal appeared. To construct trees representing the information available >N months prior to the establishment of a new antigenic cluster, we carried out the same procedure with N months subtracted from the establishment date of the cluster.

We remove all branches with no remaining descendant sequences. This is necessary because some branches may have estimated dates before the establishment date, but could not have been observed at that time because none of the descendant sequences had yet been isolated.

When a cluster transition involved multiple cluster transition substitutions occurring successively on different branches, we analyzed each substitution independently. For each substitution, we find an establishment date using all sequences descending from the occurrence of the substitution ancestral to the descendant cluster; and when producing the temporally early tree, we remove all sequences which descend from that occurrence of the substitution.

#### Calculating ranks

When presenting rankings of amino acid substitutions, we consider substitutions whose ancestral amino acid matches the consensus sequence of the ancestral cluster and that satisfy the following criteria:

1. The substitution has more than one observed or expected occurrence.
2. The substitution has a positive LCR.
3. The substitution is not a reversion to the ancestral cluster consensus sequence. Immediate reversions to the ancestral cluster are excluded as they are unlikely to confer escape from population immunity and therefore cause an antigenic cluster transition.

For substitutions not meeting these criteria, the rank is reported as “>n”, where *n* is the number of substitutions which fulfill the criteria. We do not include the occurrence of the cluster transition substitution which is ancestral to the descendant cluster (such occurrences are placed within the ancestral cluster when a descendant cluster was caused by multiple cluster transition substitutions which occurred on different branches).

#### Regression of substitution proportions

In Fig. 5A, we show: (i) the proportion of single nucleotide amino acid substitutions caused by each type of nucleotide mutation (y axis, black points), (ii) their relative neutral sampling rates (x axis), and (iii) the mutation-rate independent proportions (y axis, pale grey points). Respectively, we calculate these by summing (over mutations of each of the 12 types): (i) the observed number of occurrences, (ii) the expected numbers of occurrences, and (iii) the branch set sizes; and then normalizing the values to sum to one. For each nucleotide mutation, we only consider codon contexts in which it is non-synonymous. For the panel showing cluster transition substitutions, we consider only the seven key amino acid positions from Koel et al. 2013.

The relationship between the relative sampling rates and the proportion of non-synonymous mutations caused by each type of nucleotide mutation was modelled as:

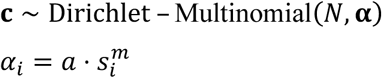

Where:

*i* indexes the 12 nucleotide mutation types
*s*_*i*_ is the relative neutral sampling rate, as described above
*m* is the exponent parameter, determining the strength of the relationship between the sampling rates (*s*_*i*_) and the proportion of non-synonymous mutations caused by each mutation type
*a* scales the concentration parameters, with larger *a* resulting in less overdispersion.
**α** = (*α*_1_, *I*, *α*_12_) are the Dirichlet concentration parameters
**c** = (*c*_1_, *c*_2_, *I*, *c*_12_) are the observed non-synonymous mutation counts (i.e. the unnormalized counts as described above);
*N* = ∑_*i*_ *c*_*i*_ is the total number of non-synonymous mutations.

We estimated the parameters *m* and *a* by maximum likelihood with the ‘optim’ function in R, using the L-BFGS-B method. Standard errors were obtained from the inverse of the observed-information matrix, computed by numerical differentiation. These standard errors were used to construct confidence intervals, compute p-values, and to generate samples from the asymptotic normal distribution of the parameter estimates.

Fig. 5A plots regression lines *y* ∝ *s*^*m*^, normalized by 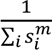 so the 12 *s*_*i*_^*m*^ sum to one.

#### Simulating LCRs for a hypothetical cluster transition substitution

In Fig. 5B, we report the median p-value as a function of the true LCR (LCR*) and the neutral expected number of occurrences (n_exp_). For each combination of LCR* and n_exp_ in a grid of values, we simulate 10^6^ values of the observed number of occurrences (n_obs_):

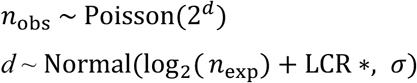

We use *σ* = 0.8, the median of the values estimated across the 12 mutation types. For each simulated n_obs_ value, we calculate the corresponding estimated LCR and p-value using the n_exp_ and *σ* = 0.8 used in the simulation. We display the median p-value in Fig. 5B, and the proportion of LCR estimates above zero in Fig. S18A.

#### n_exp_ distributions for different substitution types

In Fig. S19, we show empirical cumulative distribution functions for n_exp_ in a hypothetical cluster containing 40,000 sequences, for four scenarios: (a) all amino acid substitutions accessible by a single nucleotide mutation; (b) all substitutions observed on the tree; (c) substitutions observed on the trunk of the tree; and (d) observed cluster transition substitutions. To calculate the CDF for all accessible substitutions, for each cluster we consider all amino acid substitutions away from the cluster consensus sequence. We obtain the n_exp_ for each substitution by summing n_exp_ values for all nucleotide mutations which could have caused the substitution from the cluster consensus sequence. For each mutation, we obtain n_exp_ by scaling the n_exp_ of the corresponding mutation type in DA21 (*n*_*exp*_^*i*,*DA*21^) by the ratio 40,000 / number of sequences in DA21.

To produce the remaining three CDFs for the sets of observed substitutions, we use the same procedure, except considering amino acid substitutions which (respectively) occurred anywhere on the tree, on the trunk of the tree, or causing a cluster transition; again we consider all single nucleotide mutations which could cause each substitution, now from the codon present in the parental reconstructed sequence of the branch where each substitution occurred.

The empirical CDFs for the 40,000 sequence cluster therefore assume the same sequencing intensity as DA21. We present empirical CDFs using the DA21 sequencing intensity, rather than another cluster, as the DA21 sequencing intensity is more applicable to the present and near future than the lower sequencing intensities of earlier clusters, which would result in (inappropriately) higher estimates of n_exp_ for the same number of sequences.

In Figs. 5C & S18B, we estimate the marginal distribution of p-values and LCR estimates, integrated over the n_exp_ distributions described above. For each value of LCR*, we draw 10^4^ samples with replacement from the empirical n_exp_ distribution. For each sampled n_exp_, we simulate one n_obs_ as in Fig. 5B, and calculate the resulting LCR and p-value. The median p-value and proportion of LCR estimates above zero across the 10^4^ samples are reported in Figs. 5C & S18B, respectively.

#### Estimating dates of internal branches

To calculate per-year log convergence ratios for the DA21 cluster, internal branches were assigned dates equal to the date of the nearest descendant tip (by branch length) with a non-missing collection date. Branches were then separated into April-to-March time bins, and LCRs were calculated separately for each set of branches.

The time-resolved phylogeny in Fig. S23 was produced using Chronumental (*85*).

#### Modelling the effect of sequencing intensity on n_exp_

The expected number of occurrences does not scale linearly with sequencing intensity, because as more sequences are collected from the population, each additional sequence is increasingly closely related to those already sampled and is therefore expected to contain fewer new mutations. The exact relationship depends on the distribution of branch lengths in the tree, which is influenced by factors such as selection, population dynamics, and sequence sampling strategy. To characterize this relationship for our set of influenza H3 sequences, we therefore used a subsampling approach.

Sequencing intensity was varied by randomly discarding sequences from the tree, ranging from a minimum intensity (removing 90% of sequences) to a maximum intensity (no sequences removed). For each subsampled tree, the total branch length was calculated as a proxy for the expected number of occurrences. We fitted a power-law model to the resulting data:

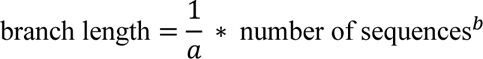

The exponent *b* determines the relative rate at which branch length, and therefore the expected number of occurrences, changes as the number of sequences is increased. We repeated this process for each cluster in the tree independently, and found an average value of *b* = ∼0.69 across clusters with >200 sequences, which was consistent across clusters spanning a range of ∼200 to ∼70,000 sequences (Figs. S20 & S21). This implies that to double the branch length, the number of sequences needs to be increased by 2^1/0.69^ = 2.7x.

We further validated the value obtained from this approach by comparing the expected occurrence counts in the full tree to the 10% subsampled trees used in Fig. S22, obtaining a similar value for *b* of 0.65.

#### Supplementary Text

##### Synonymous mutation rate variation

There are two sources of variation which contribute to uncertainty in LCR estimates. The first is sampling variation, which results from the fact that the observed number of occurrences represents a single instance of a stochastic evolutionary and sequence collection process. This is analogous to sampling variation in any statistic calculated from a finite sample, and therefore decreases asymptotically to zero with increased sampling. We determined that this variation should be modelled as a Poisson process by comparing occurrence counts of individual synonymous nucleotide mutations between non-overlapping subtrees, which represent independent realizations of this sampling process (Fig. S2; see Methods section *Testing the Poisson model of sampling variation* for more details).

The second source of variation is variation in synonymous mutation rate. We find that the number of occurrences of each type (e.g. C to T) of nucleotide mutation differs substantially between four-fold synonymous positions—more than can be explained by sampling variation (Fig. S3). In other words, a synonymous C to T mutation at genome position X has a different underlying mutation rate to a synonymous C to T mutation at genome position Y. Existing site-wise dN/dS methods account for this type of variation by estimating a distinct synonymous mutation rate for each codon, either explicitly as a parameter which multiplies the instantaneous rate matrix (*86*–*89*), or implicitly by counting synonymous mutations independently in each codon (*22*, *25*, *89*). However, we find no correlation in the mutation rate between different synonymous mutations at the same four-fold synonymous position (Fig. S4). The rate of synonymous mutations in a codon is therefore not likely to be informative about the “latent” synonymous mutation rate for non-synonymous mutations in the codon (i.e. the rate of the non-synonymous mutations *if they were synonymous*). It is therefore not appropriate to use codon-specific synonymous mutation rates to calculate or adjust the expected number of occurrences of non-synonymous mutations.

As such, we include uncertainty about the latent synonymous mutation rate in our null model of evolution. To achieve statistical significance, the LCR must therefore be large enough that it cannot be explained by synonymous rate variation: for example, the smallest LCR which can be significant at the 10% level is ∼1.0 when using a standard deviation of 0.8 for the synonymous mutation rate (the median among the estimates for the 12 nucleotide mutation types). This is the case regardless of the quantity of sampled sequences—because collecting more sequences does not give additional information on the latent synonymous mutation rate for a particular nucleotide mutation—meaning synonymous rate variation is the main barrier to achieving statistical significance when the expected number of occurrences is large.

#### Assignment of antigenic clusters and causative cluster transition substitutions

The antigenic evolution of influenza H3N2 viruses is organized into clusters of antigenically similar viruses, with substitutions at a small number of key positions near to the receptor binding site largely responsible for differences between these clusters (*11*). In this manuscript, the targets of prediction are these antigenic substitutions, which have been retrospectively identified by numerous methods. In this section of the Supplementary text, we describe the evidence supporting the assignment of antigenic clusters, and the identification of the causative antigenic cluster transition substitutions in each case.

The subsection *Assignment of antigenic clusters* provides an overview of the evidence relating to the assignment of antigenic clusters, summarised in Fig. S6, with the supporting data described in subsection *Detailed description of data supporting the assignment of antigenic clusters*.

The subsection *Estimation of causative antigenic cluster transition substitutions* provides an overview of evidence relating to the identification of causative substitutions, summarised in Fig. S7, with the supporting data described in subsection *Detailed description of data supporting the assignment of causative cluster transition substitutions*.

#### Assignment of antigenic clusters

For H3N2 viruses between 1968 and 2003, we follow the assignment of antigenic clusters in Smith et al. 2004, which is also used in Bedford et al. 2014, and Fonville et al. 2014 (*1*, *49*, *50*). For H3N2 viruses between 2003 and 2011, we used Russell et al. 2008 (*48*), Bedford et al. 2014, Fonville et al. 2014 to assign antigenic clusters. There was one instance where clusters were assigned differently between these publications: Bedford et al. 2014 assigns a new antigenic cluster for the Brisbane/10/2007 vaccine strain, while Fonville et al. 2014 includes it in the WI05 cluster. To resolve this, we used HI data from the Francis Crick Institute reports for WHO Vaccine Composition Meetings (subsequently “FCI VCM reports”) (*60*) to check whether ferret sera raised against Wisconsin/67/2005 had titers to Brisbane/10/2007 which were substantially lower than homologous. We found that the mean fold reduction was small (0.3 log_2_ units), so include Brisbane/10/2007 in the WI05 cluster.

The Bedford et al. 2014 and Fonville et al. 2014 publications characterise H3N2 influenza viruses up to approximately 2011. For the H3N2 evolution since 2011, we therefore used the list of WHO recommended vaccine strains, and checked whether each successive vaccine strain represented a large antigenic change from the previous vaccine strain. We grouped together vaccine strains separated only by small antigenic changes, and assigned a new cluster when there was a large antigenic change. The size of each antigenic change was assessed using both research articles, which each individually cover short periods of the post-2011 evolution (*51*, *52*), and using HI data from the FCI VCM reports, similarly to the assessment of whether Brisbane/10/2007 should be included in the same cluster as Wisconsin/67/2005.

This method resulted in the assignment of the SW13, HK14, KA17, HK19, CA20, and DA21 antigenic clusters. In every case where one of these clusters contained more than one WHO recommended vaccine strain, the fold reduction in titer from the first vaccine strain to each of the later vaccine strains was ≤2-fold for at least 83% of the titer tables in the FCI VCM reports. For every vaccine update where we assigned a new cluster, the fold reduction from the final vaccine strain in the previous cluster to the first vaccine strain of the new cluster was ≥4-fold for at least 60% of the titer tables in the FCI VCM reports.

A detailed description of the data pertaining to each antigenic cluster transition since 2003 is provided in the subsection *Detailed description of data supporting the assignment of antigenic clusters*, and an overview schematic is provided in Fig. S6.

### Estimation of causative antigenic cluster transition substitutions

For antigenic cluster transitions between 1968 and 2003, Koel et al. 2013 (*11*) identified the substitutions which caused the antigenic change using reverse genetics. For each cluster transition, amino acid substitutions which differed between the ancestral and descendant cluster were identified (“cluster-difference substitutions”) and introduced (individually and in combinations) into a virus from the ancestral cluster. The resultant mutant viruses were antigenically characterized using hemagglutination inhibition or neutralization assays against ferret antisera. An antigenic map was produced, allowing the mutant virus(es) whose antigenic phenotype was similar to the descendant cluster to be identified. For cluster transitions between 1968 and 2003, we use the substitutions identified by Koel et al. 2013.

For later antigenic cluster transitions, we use estimates of substitution-level antigenic effects from multiple sources (*51*, *53*–*57*, *59*). While one (Chambers et al. 2015, (*51*)) uses a similar reverse genetics approach to Koel et al. 2013, most of these sources infer the antigenic effect of substitutions from titrations performed with naturally circulating viruses.

Pattinson 2019 (*53*) searched for “natural experiments” in antigenic maps covering evolution between 2003 and 2016. Sets of viruses which differ only by a single substitution were identified, and the positioning of these viruses in antigenic maps was then examined, allowing the impact of each cluster-difference substitution to be individually determined—producing a comparison similar to the one achieved by the reverse genetics approaches of Koel et al. 2013 and Chambers et al. 2015. The effect of each cluster-difference substitution was further tested in a formal linear mixed model framework. This approach was used to identify the substitutions responsible for cluster transitions between FU02 and HK14.

For H3N2 viruses up to 2016, Neher et al. 2016 (*54*) estimate the effect of individual substitutions on HI titer using a linear model framework, similar to that in Pattinson 2019, except estimating the effect of substitutions on HI titer directly rather than on position in an antigenic map. We used the estimates in Neher et al. 2016 to cross-check the cluster transition substitutions identified in Pattinson 2019, by checking whether the cluster transition substitutions identified in Pattinson 2019 had the largest estimated effect on HI titer out of any of the cluster-difference substitutions in Neher et al. 2016. We found that this was always the case.

For the WI05 to PE09 cluster transition, multiple studies investigated the antigenic properties of two clades (represented by the Victoria/208/2009 and Perth/16/2009 viruses) which emerged in parallel and which both contained the K158N and N189K substitutions. These studies determined them to be antigenically similar (*55*–*57*), with a parsimonious explanation being that these two substitutions were responsible for the change in antigenic phenotype.

For H3N2 viruses between 2011 and 2018, Jorquera et al. 2019 (*59*) performed titrations of many viruses from the 3C.3a (SW13) and 3C.2a (HK14) subclades. Multiple of these independently acquired the F193S substitution, the presence or absence of which was the primary determinant of titer between 3C.3a and 3C.2a subclades and 3C or 3C.3a sera. They therefore identify it as causative of antigenic distancing of these viruses from these sera.

On numerous occasions, these analyses overlap in the time periods they cover. When they do, they are consistent in which substitution(s) they identify as causing antigenic change: every cluster transition substitution identified in Pattinson 2019 had a larger estimated effect size in Neher et al. 2016 than any of the other cluster-difference substitutions; for PE09 to SW13, the F159S substitution identified by Pattinson 2019 and Neher et al. 2016 is also identified by the reverse genetics approach of Chambers et al. 2015; and for WI05 to PE09, in addition to the parsimony argument based on the two parallel clades containing K158N and N189K, these substitutions have a larger effect size in Neher et al. 2016 than any other cluster-difference substitutions.

For the most recent cluster transitions (from HK14 to CA20 and CA20 to DA21), there is less data available to estimate the causative antigenic substitution, so our assignments of causative antigenic substitutions are made with lower confidence. For HK14 to CA20, we identify F193S as the causative substitution based on a similar parsimony argument to the WI05 to PE09 case—except that we neither have as much data to compare the antigenic phenotypes of HK19 and CA20, nor have substitution-level estimates from Neher et al. 2016 for this time period. Because of this lower confidence, and because F193S already caused a cluster transition from the HK14 cluster to the HK19 cluster, we do not include this cluster transition in our retrospective analysis—a prediction of F193S from the HK14 cluster is not particularly informative once it has already caused the cluster transition from the HK14 cluster. Nonetheless, we define both a CA20 and HK19 cluster, despite both being caused by an F193S substitution from HK14, because the DA21 cluster evolved from CA20-like viruses (rather than HK19-like viruses).

For CA20 to DA21, we identify Y159N as the causative substitution based on its presence in the Koel-7 group of positions, and the observation that it causes escape from neutralisation by human sera when added to a virus from the HK19 antigenic cluster (Welsh et al. 2024, (*58*) – though it should be noted that the transition to DA21 occurred from CA20 rather than HK19 as tested in Welsh et al. 2024). It is therefore the lowest confidence assignment of any of the cluster transition substitutions we use in the retrospective evaluation. However, we include it nonetheless, because it is one of two cases since 1987 where the cluster transition substitution was *not* successfully identified using convergence data, so is informative to consider from that perspective.

A detailed summary of the data pertaining to each antigenic cluster transition substitution since 2003 is provided in the subsection *Detailed description of data supporting the assignment of causative cluster transition substitutions*, and a schematic provided in Fig. S7.

### Detailed description of data supporting the assignment of antigenic clusters

For H3N2 viruses between 2003 and 2011, we follow the assignment of antigenic clusters from Fonville et al. 2014. We additionally checked that, when there was more than one WHO vaccine strain recommendation within a cluster, published data (*48*–*50*, *60*) supported their placement in the same antigenic cluster (i.e. that they were not substantially antigenically divergent):

#### Fujian/411/2002 and Wellington/1/2004 vaccine strains both placed in the FU02 cluster

This cluster assignment is consistent with Russell et al. 2008, and Fonville et al. 2014, and Bedford et al. 2014 (*48*–*50*).

#### Wisconsin/67/2005 and Brisbane/10/2007 both placed in the WI05 cluster

This cluster assignment is consistent with Russell et al. 2008, and Fonville et al. 2014; but is inconsistent with Bedford et al. 2014, which assigns a separate cluster containing Brisbane/10/2007.

We used HI data from reports prepared by the Francis Crick Institute for WHO Vaccine Composition Meetings (*60*) to check whether a ferret serum raised against the Wisconsin/67/2005 virus had substantially lower titers to the Brisbane/10/2007 virus than to the homologous Wisconsin/67/2005 virus. In FCI VCM reports between September 2007 and February 2010, the geometric mean fold-reduction in titer from Wisconsin/67/2005 to Brisbane/10/2007 against anti-Wisconsin/67/2005 sera was 0.3 log_2_ units, with 27% of fold-reductions greater than or equal to 2 log_2_ units across the titer tables (n = 11). We therefore concluded that Brisbane/10/2007 does not represent a large antigenic change from Wisconsin/67/2005, so we consider it within the same antigenic cluster.

#### Perth/16/2009 and Victoria/361/2011 vaccine strains both placed in the PE09 cluster

Bedford et al. 2014 and Fonville et al. 2014 both contain viruses from 2011, and do not identify a separate Victoria 2011 cluster.

For H3N2 virus evolution since 2011, we used the list of WHO recommended vaccine strains and checked in published data whether each successive vaccine strain represented a large or small antigenic change from the previous vaccine strain. When multiple passages of the same isolate were available in a titer table in an FCI VCM report, we selected the passage that most closely matched that of the virus it was being compared against (e.g., both egg-passaged or both cell-passaged where possible).

#### Texas/50/2012 vaccine strain is placed in the PE09 antigenic cluster

We do not assign a new antigenic cluster for the Texas/50/2012 antigen, based on the following data:

- In FCI VCM reports from September 2013 and February 2014, the geometric mean fold-reduction in titer from Perth/16/2009 to Texas/50/2012 against anti-Perth/16/2009 sera was 0.6 log_2_ units, with 7% of fold-reductions greater than or equal to 2 log_2_ units (n = 27).

#### A new SW13 antigenic cluster, containing Switzerland/9715293/2013-like viruses, is assigned as a descendant of the PE09 antigenic cluster

The Switzerland/9715293/2013 vaccine strain is a member of the 3C.3a clade, which is a descendant of the 3C clade containing the Texas/50/2012 vaccine strain. We assign a new antigenic cluster for Switzerland/9715293/2013-like viruses based on the following data:

- In the FCI VCM report from February 2015, the geometric mean fold-reduction in titer from Texas/50/2012 (the last vaccine strain in the PE09 cluster) to Switzerland/9715293/2013 against anti-Texas/50/2012 sera was 2.9 log_2_ units, with 100% of fold-reductions greater than or equal to 2 log_2_ units (n = 13).
- Table 2 of Chambers et al. 2015 (*51*) shows a fold-reduction in titer from Texas/50/2012 to Switzerland/9715293/2013 of 4 log_2_ units using an anti-Texas/50/2012 ferret serum, and 3 log_2_ units using an anti-Texas/50/2012 sheep serum.
- The antigenic map in Fig. 6 of Li et al. 2016 (*52*) shows substantial antigenic divergence between the Texas/50/2012 antigen and the Switzerland/9715293/2013 antigen.

#### A new KA17 antigenic cluster, containing Kansas/14/2017-like viruses, is assigned as a descendant of the SW13 antigenic cluster

The Kansas/14/2017 vaccine strain is a member of the 3C.3a1 clade, which is a descendant of the 3C.3a clade containing the Switzerland/9715293/2013 vaccine strain. We assign a new antigenic cluster for Kansas/14/2017-like viruses based on the following data:

- In the FCI VCM reports from September 2018 and September 2019, the geometric mean fold-reduction in titer from Stockholm/6/2014 (this is the representative 3C.3a virus used in these FCI VCM reports) to Kansas/14/2017 against anti-Stockholm/6/2014 sera was 3.3 log_2_ units, with 100% of fold-reductions greater than or equal to 2 log_2_ units (n = 3).

#### A new HK14 antigenic cluster, containing Hong Kong/4801/2014-like viruses, is assigned as a descendant of the PE09 antigenic cluster

The Hong Kong/4801/2014 vaccine strain is a member of the 3C.2a clade, which is a descendant of the 3C clade containing the Texas/50/2012 vaccine strain. We assign a new antigenic cluster for Hong Kong/4801/2014-like viruses based on the following data:

- In the FCI VCM report from February 2016, the geometric mean fold-reduction in titer from Texas/50/2012 (the last vaccine strain in the PE09 cluster) to Hong Kong/4801/2014 against anti-Texas/50/2012 sera was 5.9 log_2_ units, with 100% of fold-reductions greater than or equal to 2 log_2_ units (n = 7).
- The antigenic map in Fig. 6 of Li et al. 2016 shows substantial antigenic divergence between the Texas/50/2012 antigen and the Hong Kong/4801/2014 antigen.

#### Singapore/Infimh-16-0019/2016, Switzerland/8060/2017, and South Australia/34/2019 vaccine strains are placed in the HK14 antigenic cluster

The Singapore/Infimh-16-0019/2016 vaccine strain is a member of the 3C.2a1 clade, which is a descendant of the 3C.2a clade containing the Hong Kong/4801/2014 vaccine strain. We do not assign a new antigenic cluster for Singapore/Infimh-16-0019/2016-like viruses based on the following data:

- In the FCI VCM report from February 2020, the geometric mean fold-reduction in titer from Hong Kong/5738/2014 (this is the 3C.2a representative virus in this FCI VCM report) to Singapore/Infimh-16-0019/2016 against anti-Hong Kong/5738/2014 sera was 0.5 log_2_ units, with 0% of fold-reductions greater than or equal to 2 log_2_ units (n = 12).

The Switzerland/8060/2017 vaccine strain is a member of the 3C.2a2 clade, which is a descendant of the 3C.2a clade containing the Hong Kong/4801/2014 vaccine strain. We do not assign a new antigenic cluster for Switzerland/8060/2017-like viruses based on the following data:

- In the FCI VCM report from February 2019, the geometric mean fold-reduction in titer from Hong Kong/5738/2014 (this is the 3C.2a representative virus in this FCI VCM report) to Switzerland/8060/2017 against anti-Hong Kong/5738/2014 sera was 0.0 log_2_ units, with 0% of fold-reductions greater than or equal to 2 log_2_ units (n = 11).

The South Australia/34/2019 vaccine strain is a member of the 3C.2a1b.2 clade, which is a descendant of the 3C.2a clade containing the Hong Kong/4801/2014 vaccine strain, and the 3C.2a1 clade containing the Singapore/Infimh-16-0019/2016 vaccine strain. We do not assign a new antigenic cluster for South Australia/34/2019-like viruses based on the following data:

- In the FCI VCM report from February 2020, the geometric mean fold-reduction in titer from Hong Kong/5738/2014 (this is the 3C.2a representative virus in this FCI VCM report, and the earliest vaccine strain in the HK14 cluster is a 3C.2a virus) to South Australia/34/2019 against anti-Hong Kong/5738/2014 sera was -0.2 log_2_ units, with 0% of fold-reductions greater than or equal to 2 log_2_ units (n = 11).

#### A new HK19 antigenic cluster, containing Hong Kong/45/2019-like viruses, is assigned as a descendant of the HK14 antigenic cluster

The Hong Kong/45/2019 vaccine strain is a member of the 3C.2a1b.1b clade, which is a descendant of the 3C.2a clade containing the Hong Kong/4801/2014 vaccine strain, and the 3C.2a1 clade containing the Singapore/Infimh-16-0019/2016 vaccine strain. We assign a new antigenic cluster for Hong Kong/45/2019-like viruses based on the following data:

- In the FCI VCM report from September 2021, the geometric mean fold-reduction in titer from Singapore/Infimh-16-0019/2016 to Hong Kong/2671/2019 (this is the 3C.2a1b.1b representative virus used in this report) against anti-Singapore/Infimh-16-0019/2016 sera was 1.6 log_2_ units, with 60% of fold-reductions greater than or equal to 2 log_2_ units (n = 10).

#### A new CA20 antigenic cluster, containing Cambodia/e0826360/2020-like viruses, is assigned as a descendant of the HK14 antigenic cluster

The Cambodia/e0826360/2020 vaccine strain is a member of the 1a clade, which is a descendant of the 3C.2a clade containing the Hong Kong/4801/2014 vaccine strain, the 3C.2a1 clade containing the Singapore/Infimh-16-0019/2016 vaccine strain, and the 3C.2a1b.2 clade containing the South Australia/34/2019 vaccine strain. We assign a new antigenic cluster for Cambodia/e0826360/2020-like viruses based on the following data:

- In the FCI VCM report from September 2021, the geometric mean fold-reduction in titer from Singapore/Infimh-16-0019/2016 (the most proximate ancestral vaccine strain present in the same titer table as Cambodia/e0826360/2020) to Cambodia/e0826360/2020 against anti-Singapore/Infimh-16-0019/2016 sera was 1.9 log_2_ units, with 80% of fold-reductions greater than or equal to 2 log_2_ units (n = 10).

#### A new DA21 antigenic cluster, containing Darwin/9/2021-like viruses, is assigned as a descendant of the CA20 antigenic cluster

The Darwin/9/2021 vaccine strain is a member of the 2a clade. While not a direct phylogenetic ancestor, the most appropriate previous vaccine strain for comparison is Cambodia/e0826360/2020, as they share the antigenic F193S mutation, which is not present in the most proximate direct ancestor, South Australia/34/2019. Further, there are no FCI VCM report tables which contain both South Australia/34/2019 and Darwin/9/2021. We assign a new antigenic cluster for Darwin/9/2021-like viruses based on the following data:

- In the FCI VCM report from February 2022, the geometric mean fold-reduction in titer from Cambodia/e0826360/2020 to Darwin/9/2021 against anti-Cambodia/e0826360/2020 sera was 2.7 log_2_ units, with 91% of fold-reductions greater than or equal to 2 log_2_ units (n = 11).

#### Thailand/8/2022 and Croatia/10136RV/2023 are placed in the DA21 antigenic cluster

The Thailand/8/2022 vaccine strain is a member of the 2a.3a.1 clade, which is a descendant of the 2a clade containing the Darwin/9/2021 vaccine strain. We do not assign a new antigenic cluster for Thailand/8/2022-like viruses based on the following data:

- In the FCI VCM report from September 2024, the geometric mean fold-reduction in titer from Darwin/9/2021 to Thailand/8/2022 against anti-Darwin/9/2021 sera was 0.2 log_2_ units, with 0% of fold-reductions greater than or equal to 2 log_2_ units (n = 6).

The Croatia/10136RV/2023 vaccine strain is a member of the 2a.3a.1 clade, which is a descendant of the 2a clade containing the Darwin/9/2021 vaccine strain. We do not assign a new antigenic cluster for Croatia/10136RV/2023-like viruses based on the following data:

- In the FCI VCM report from September 2024, the geometric mean fold-reduction in titer from Darwin/9/2021 to Croatia/10136RV/2023 against anti-Darwin/9/2021 sera was 1.0 log_2_ units, with 17% of fold-reductions greater than or equal to 2 log_2_ units (n = 6).

### Detailed description of data supporting the assignment of causative cluster transition substitutions

#### FU02 to CA04, K145N

Pattinson 2019 concludes that K145N is the substitution most likely to be responsible for the FU02-CA04 cluster transition. In Neher et al. 2016 (*54*), K145N has the largest estimated antigenic effect of any of the substitutions differing between FU02 and CA04 identified in Pattinson 2019 (*53*) (K145N, Y159F, V226I, S227P) in the 1995-2005 and the 2000-2010 time periods.

#### CA04 to WI05, S193F (excluded from retrospective evaluation as it requires two nucleotide mutations in the same codon)

Pattinson 2019 concludes that S193F is the substitution most likely to be responsible for the CA04-WI05 cluster transition. In Neher et al. 2016, S193F is confounded with the only other substitution differing between these clusters, D225N, so it is not possible to separate their effects to corroborate the Pattinson 2019 conclusion. However, because the S193F substitution requires two nucleotide mutations in the same codon, it is not possible to estimate an LCR for this substitution anyway, so it is excluded from the retrospective evaluation.

#### WI05 to PE09, K158N & N189K

In 2009, two clades independently emerged containing both the K158N and N189K substitutions: the Victoria/208/2009-like clade, which additionally contained the T212A substitution, and the Perth/16/2009-like clade, which additionally contained the E62K and N144K substitutions. The two clades have been shown to have very similar antigenic properties on numerous occasions, variously described as antigenically “indistinguishable” (*57*), “similar” (*56*), or “related” (*55*). The most parsimonious explanation is that these antigenic properties are caused by the shared K158N and N189K mutations (*53*).

In Neher et al. 2016, the K158N+N189K combination has a larger estimated antigenic effect than any of E62K, N144K, or T212A in the 2000-2010 and 2005-2016 time periods.

#### PE09 to SW13, F159S

The causative antigenic substitution for the PE09 to SW13 cluster transition was identified using reverse genetics in Chambers et al. 2015 (*51*), who find that F159S is principally responsible for the antigenic change. Pattinson 2019 also identifies the F159S substitution as responsible for this cluster transition. In Neher et al. 2016, F159S has the largest estimated antigenic effect out of the substitutions differing between these clusters identified in Pattinson 2019 (A138S, F159S, N225D) in the 2005-2016 time period.

#### PE09 to HK14, F159Y

Pattinson 2019 concludes that F159Y is the substitution most likely to be responsible for the PE09-HK14 cluster transition.

In Neher et al. 2016, F159Y has the largest antigenic effect size of any of the substitutions differing between PE09 and HK14 identified in Pattinson 2019 (A128T, G142R, N144S, F159Y, K160T, N225D) in the 2005-2016 time period.

#### SW13 to KA17, F193S

Jorquera et al. 2019 (*59*) characterised 3C.3a viruses which contain only the F193S mutation on the background of the 3C.3a clade consensus, as well as KA17 sequences containing all of L3I, S91N, N144K, and F193S. They found that 3C.3a sequences with only the F193S substitution have a very similar antigenic phenotype to the sequences with the full complement of KA17 mutations, so F193S can be inferred as responsible for the antigenic change at this cluster transition.

#### HK14 to HK19, F193S

Jorquera et al. 2019 characterised 3C.2a1 (Singapore/Infimh-16-0019/2016-like) viruses which contain only the F193S mutation on the background of the 3C.2a1 clade consensus. They also characterised 3C.2a1 viruses with all of E62G, K92R, N121K, T135K, R142G, H311Q. This represents the majority of the substitutions differing between the 3C.2a1 and 3C.2a1b.1b (HK19) clades, which further adds only the F193S, S137F, and A138S substitutions. The F193S-only viruses show a substantially larger (∼3-fold) reduction in titer compared to the 3C.2a1 clade consensus than the viruses containing all of E62G+K92R+N121K+T135K+R142G+H311Q. It is possible that the S137F or A138S present alongside F193S in 3C.2a1b.1b (HK19) viruses cause further antigenic change; however, the 137 and 138 positions have not typically been associated with large antigenic changes: in Neher et al. 2016, the largest antigenic effect size associated with any mutation at either position 137 or 138 in any of the five time windows between 1985 and 2016 was 0.55 antigenic units, compared to 1.55 antigenic units for position 193.

#### HK14 to CA20, F193S (lower confidence, and excluded from retrospective evaluation as F193S previously caused a cluster transition from the HK14 cluster to the HK19 cluster)

There is relatively little data which can be used to directly estimate the substitution causing the HK14 to CA20 cluster transition. However, CA20 and HK19 exhibit similar levels of antigenic escape from HK14—of 1.6 and 1.9 antigenic units respectively in FCI VCM reports (Fig. S6) (*60*). As the CA20 and HK19 clusters share the F193S substitution, a parsimonious explanation is that F193S is also responsible for the HK14 to CA20 cluster transition. However, because CA20 also contains multiple additional substitutions for which we cannot estimate an effect (T131K, T135K, Y94N, Y195F, G186S, and S198P), the estimate of F193S as the causative antigenic substitution is made with lower confidence.

Because F193S had already caused a cluster transition away from the HK14 cluster (to HK19), we do not include the HK14 to CA20 cluster transition in the retrospective evaluation.

#### CA20 to DA21, Y159N (lower confidence)

There is also relatively little data available to determine the basis of the CA20-DA21 transition. However, out of the substitutions which differ between CA20 and DA21, only Y159N is at a site identified to have contributed substantially to antigenic evolution in Koel et al. 2013 (*11*), and has been shown to cause escape from neutralization by human sera in a root virus from the HK19 antigenic cluster (*58*). In any case, there are few sequences available from the CA20 antigenic cluster, making it challenging to determine the LCR for any substitution in the CA20 antigenic cluster. We nonetheless present Y159N in the retrospective evaluation, as it is informative because it was one of the two cases since 1987 where the cluster transition substitution was *not* successfully identified using convergence data.

## Supplementary Figures

**Fig. S1.**
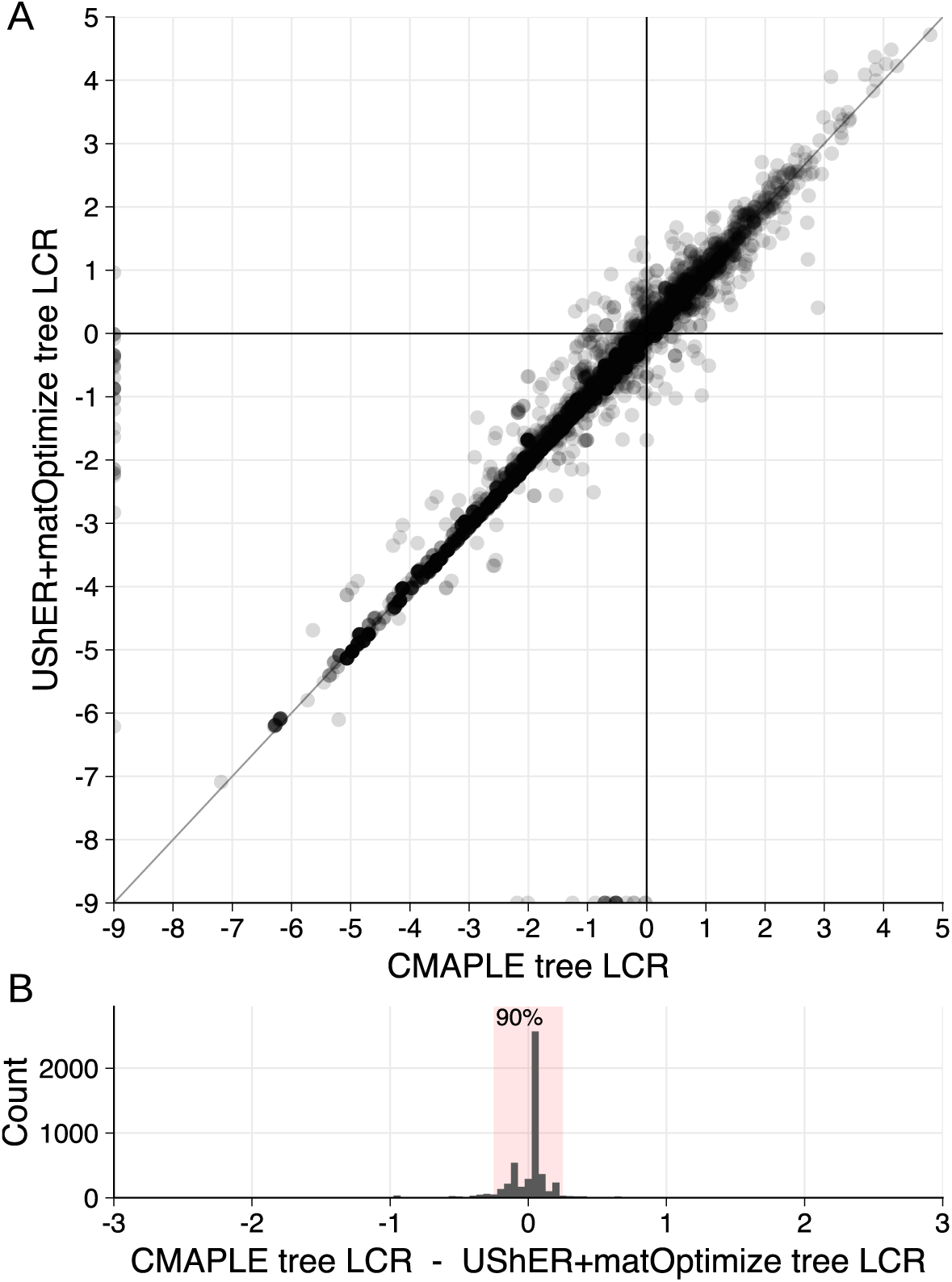
Log convergence ratio (LCR) estimates are highly concordant between trees produced using CMAPLE and using UShER + matOptimize. (**A**) Scatter plot of LCR for all substitutions with a neutral expected number of occurrences greater than one in either tree, and at least one observed occurrence. (**B**) Histogram of the difference in LCR estimates between the two trees for all substitutions from **A** with a neutral expected number of occurrences greater than one in either tree, and at least one observed occurrence in either tree. In 90% of cases, LCR estimates differ by less than 0.25 between the two trees (red shaded area). See Methods section *Tree construction* for a description of the methods used to construct each tree.

**Fig. S2.**
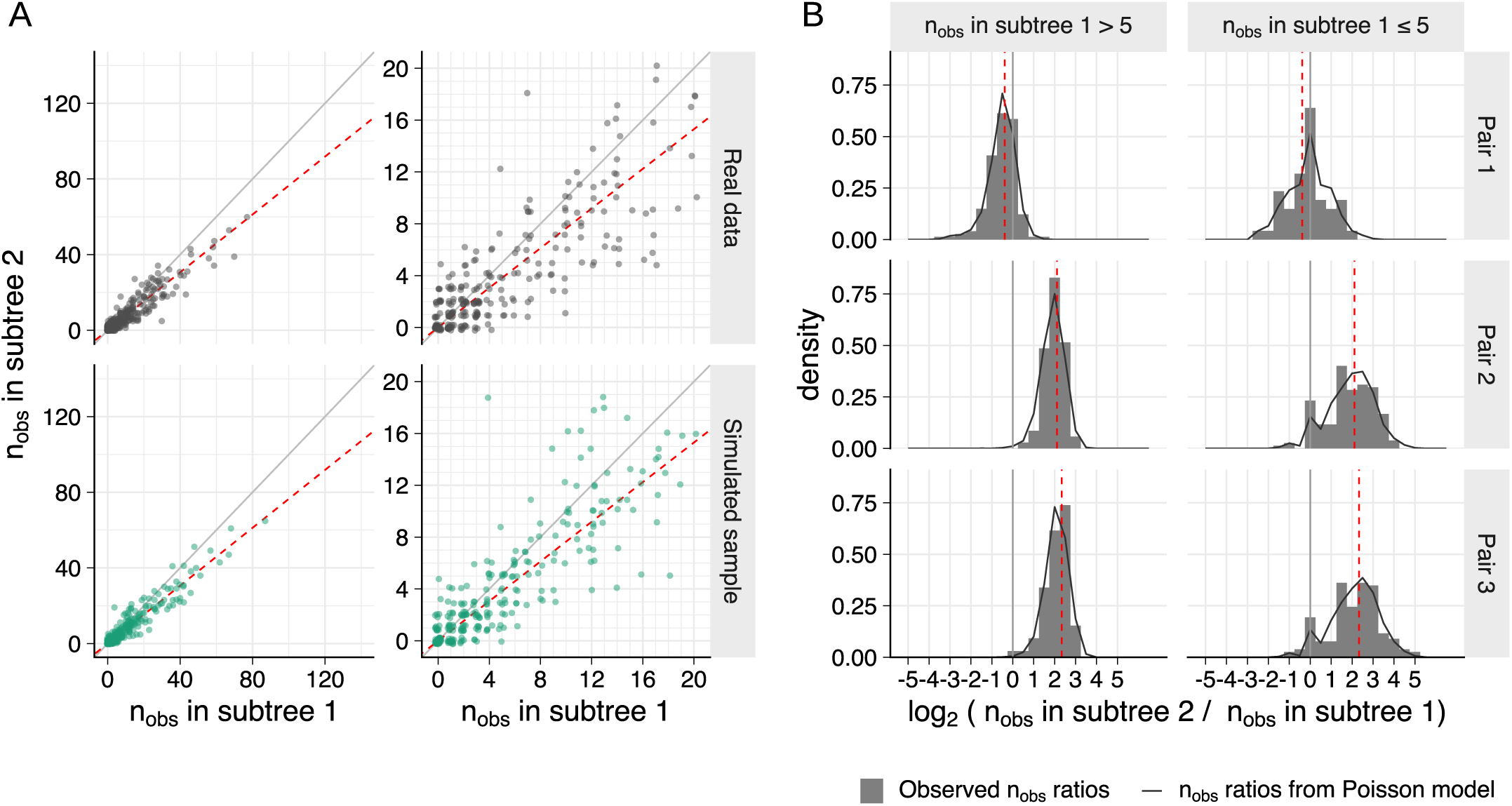
Variation in the observed number of occurrences of a nucleotide mutation at a single position is well described by a Poisson process. (**A**) The observed number of occurrences of each synonymous nucleotide mutation is plotted for two approximately evenly sized independent subtrees (top row); and for a single random draw from a Poisson process for each mutation in each subtree (bottom row), with rates equal to the expected mutation counts for the mutation in each subtree. The red dotted line indicates these expected counts, and is not at 1:1 because the subtrees differ slightly in size. Variation around this line is similar for the real observed data compared to the sample simulated using a Poisson process. The left-hand plots show the full range of the data; right-hand plots zoom in on the region of the plot near to the origin. (**B**) The grey histogram shows the distribution of the ratio of observed mutation counts between the two subtrees; the black line is the distribution of ratios obtained from 100 replicate draws of the counts from a Poisson process. The three rows show three different pairs of subtrees, obtained by splitting the tree at different branches. See Methods section *Testing the Poisson model of sampling variation* and Supplementary text section *Synonymous mutation rate variation* for more detail.

**Fig. S3.**
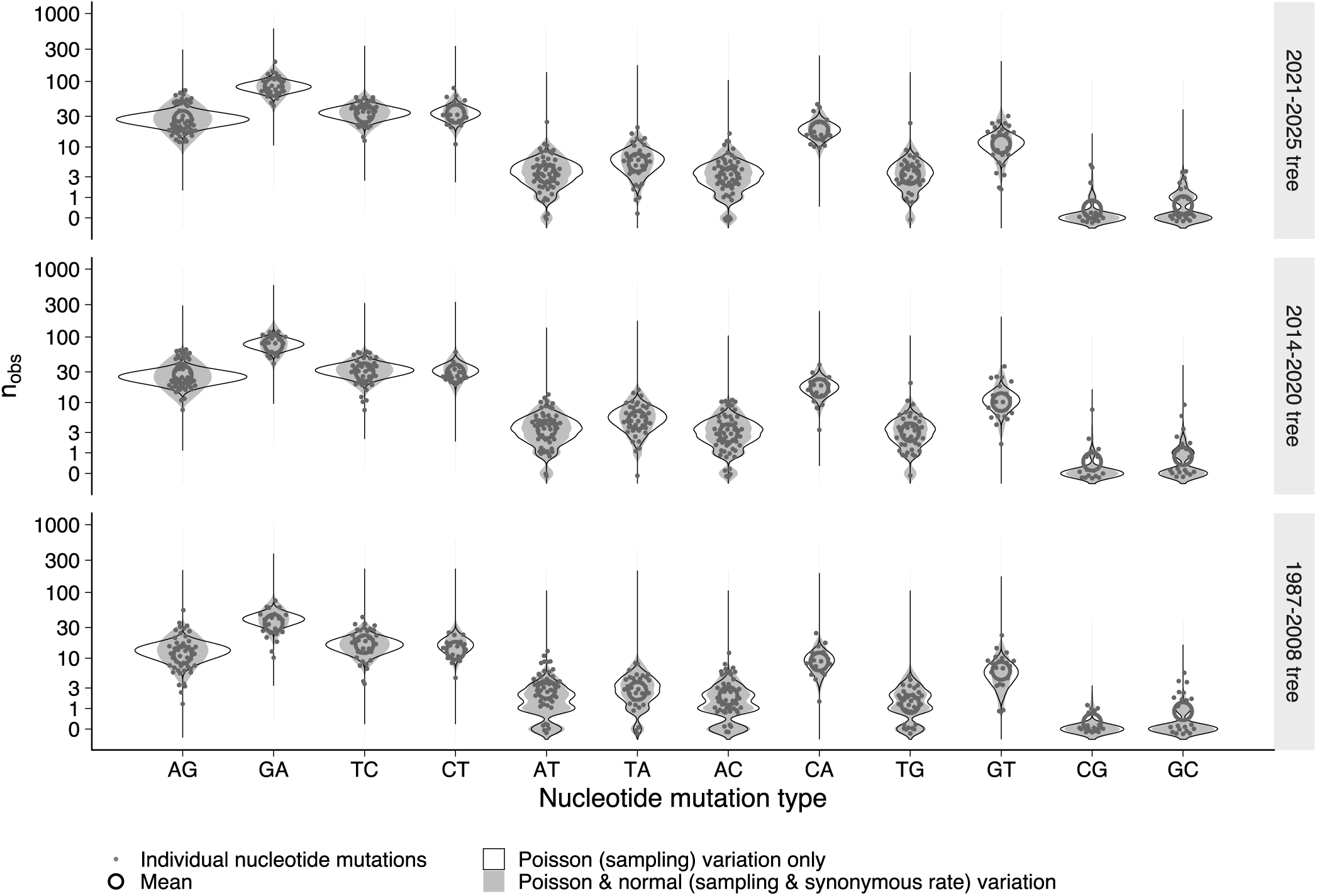
Simulated null distributions of occurrence counts closely match observed occurrence counts at four-fold synonymous positions. A null model of evolution containing Poisson distributed sampling variation and log-normally distributed variation in synonymous mutation rate (described in Methods section *Calculating the expected number of occurrences*) was fit to a tree containing sequences from approximately 2021-2025 (i.e. the Darwin 2021 antigenic cluster). Neutral occurrence counts were simulated from this model for the 2021-2025 tree, and for trees containing sequences from approximately 2014-2020 (i.e. the Hong Kong 2014 antigenic cluster) or from approximately 1987-2008 (i.e. all antigenic clusters between the Sichuan 1987 and Wisconsin 2005 antigenic clusters). Simulated distributions use the model parameters from the fit to the 2021-2025 tree for all three trees (grey shading), with the mutation rate scaled according to the relative tree size. Hollow black violins show the simulated distributions if only Poisson distributed sampling variation is included (i.e. variation in synonymous mutation rate is set to zero).

**Fig. S4.**
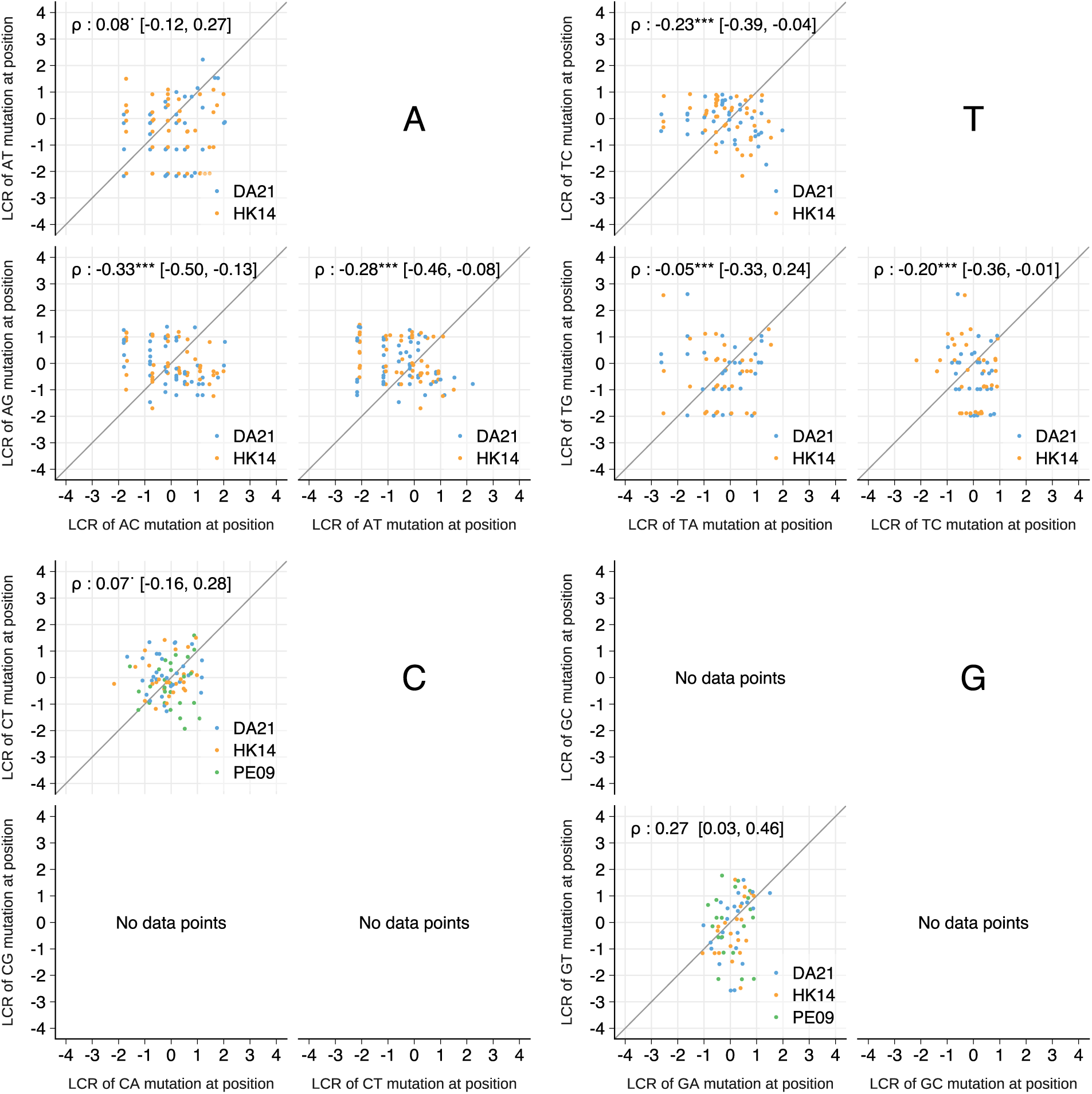
Log convergence ratios (LCR) for different synonymous mutations at the same position do not correlate. Each point represents a pair of synonymous nucleotide mutations at a single position in one of DA21, HK14, or PE09, with at least three neutral expected occurrences for both mutations in the particular cluster. The rates of different nucleotide mutations at a four-fold synonymous position do not correlate. Spearman’s rank correlation is shown with a 95% bootstrap confidence interval. ^.^ = p<0.1, * = p<0.05, ** = p<0.01, *** = p<0.001.

**Fig. S5.**
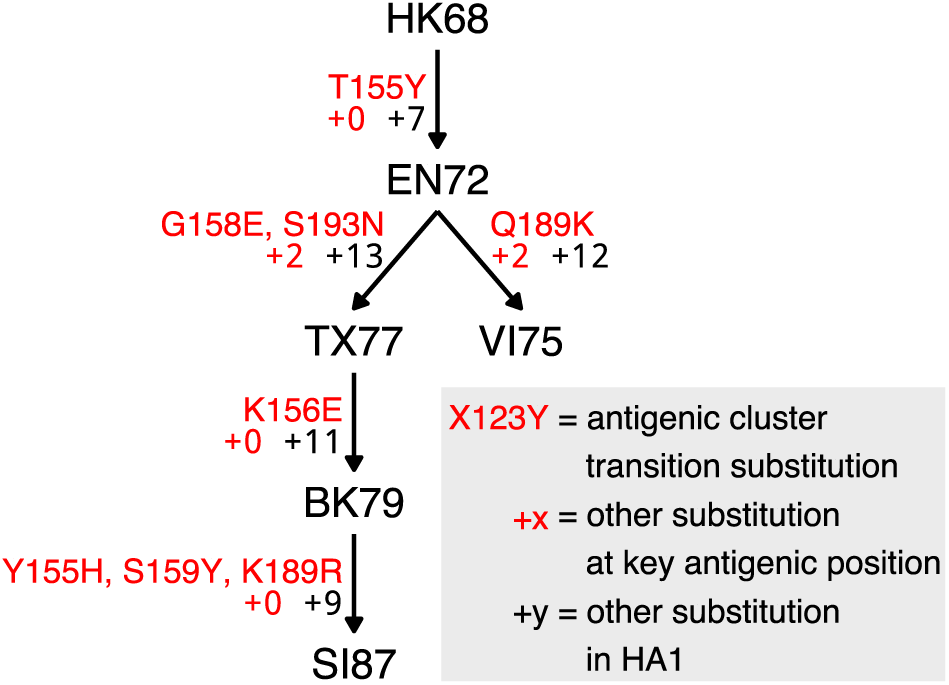
Substitutions estimated to be responsible for antigenic cluster transitions between 1968 and 1987. See Fig. 2C for antigenic cluster transitions since 1987, and Supplementary text section *Assignment of antigenic clusters and causative cluster transition substitutions* for more detail.

**Fig. S6.**
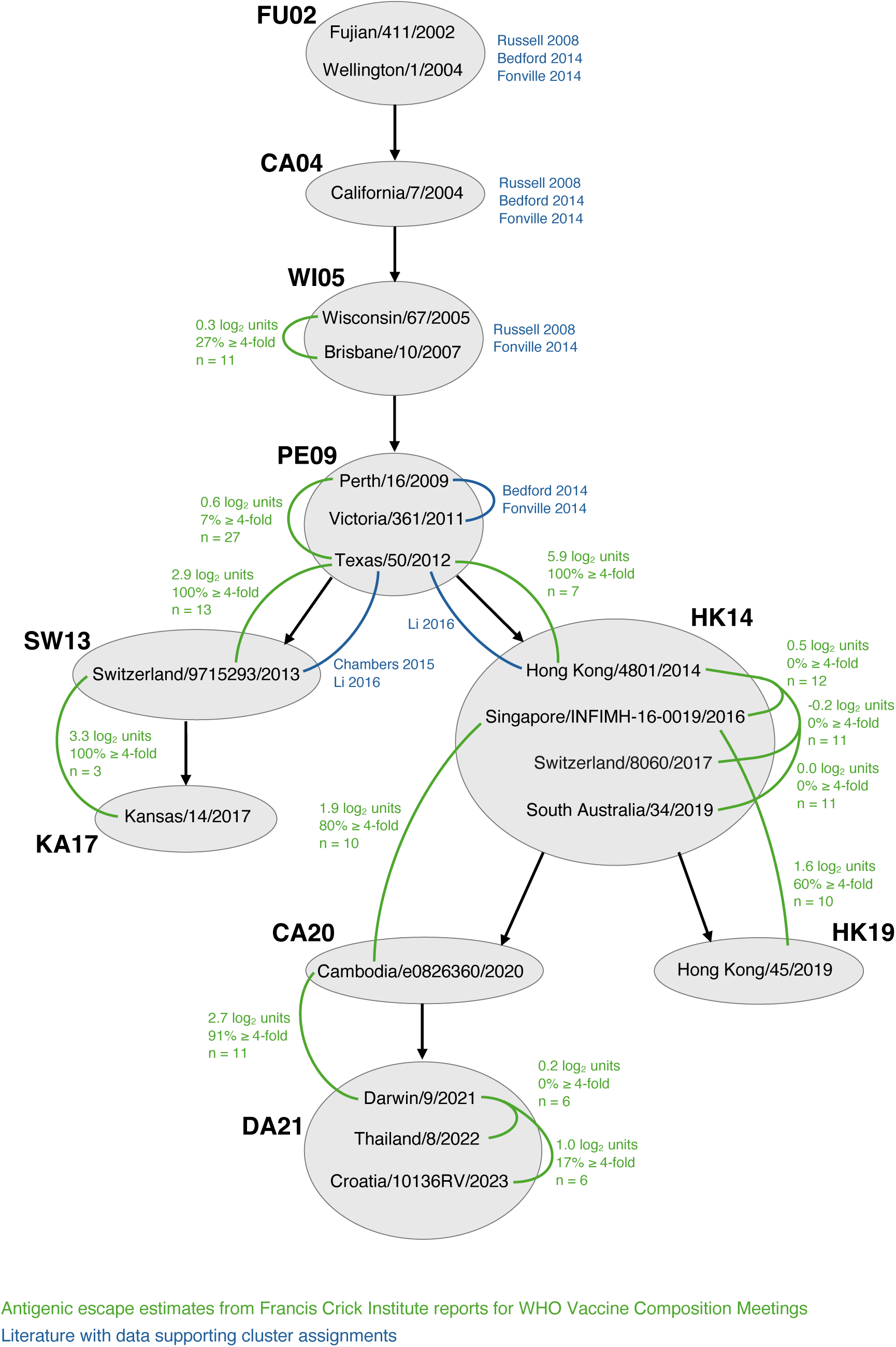
Summary of the evidence supporting the separation of post-2002 vaccine strains in antigenic clusters. Clusters up to 2002 were assigned in accordance with Smith et al. 2004, and up to 2012 in accordance with Russell et al. 2008, Bedford et al. 2014, and Fonville et al. 2014. For clusters since 2012, clusters are assigned using data from Chambers et al. 2015, Li et al. 2016, and HI data from reports prepared by the Francis Crick Institute for WHO Vaccine Composition Meetings (mean log_2_ fold-change and proportion of fold changes ≥4-fold are shown). These were used to determine whether each update to the WHO vaccine strain recommendation represented a large antigenic change and therefore a new antigenic cluster. A full description of this data is available in Supplementary text sections *Assignment of antigenic clusters* and *Detailed description of data supporting the assignment of antigenic clusters*.

**Fig. S7.**
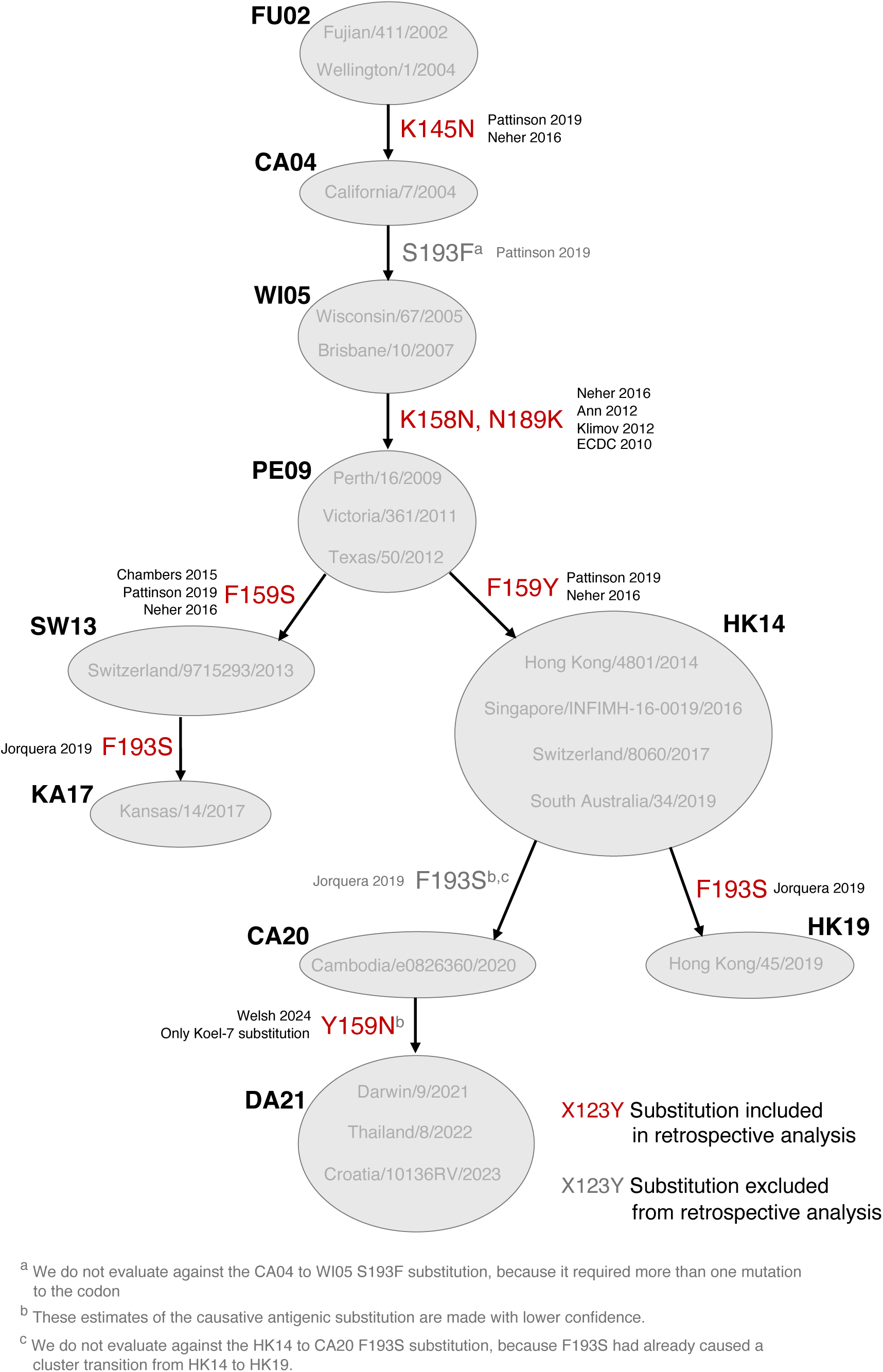
Summary of the evidence supporting the assignment of causative cluster transition substitutions for cluster transitions after FU02. Cluster transition substitutions up to the SY97 to FU02 cluster transition were assigned in accordance with Koel et al. 2013. For later clusters, we use numerous sources which estimate the effect of individual substitutions by either characterizing mutant viruses or naturally circulating viruses. A full description of this data is available in Supplementary text sections *Estimation of causative antigenic cluster transition substitutions* and *Detailed description of data supporting the assignment of causative cluster transition substitutions*.

**Fig. S8.**
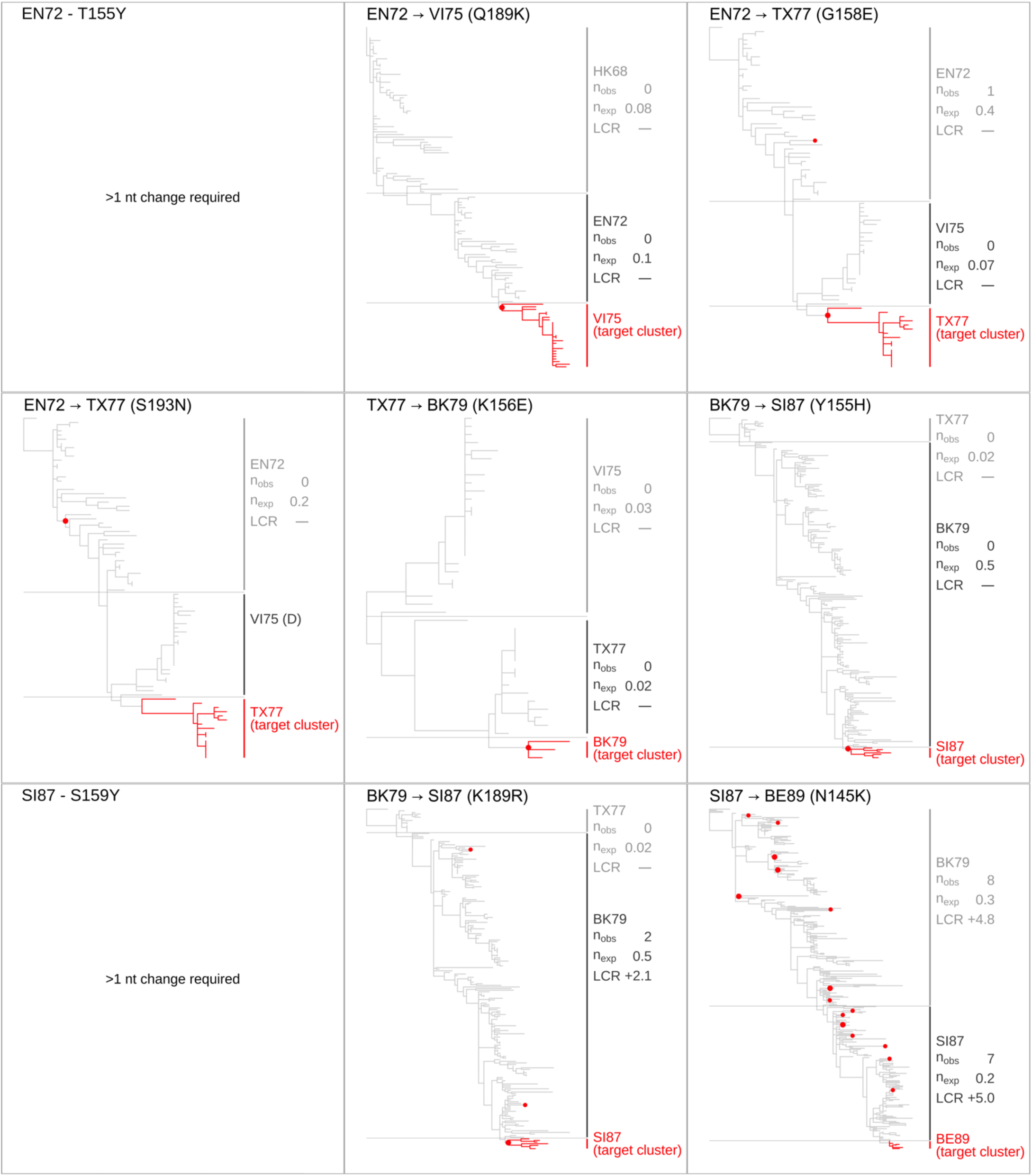
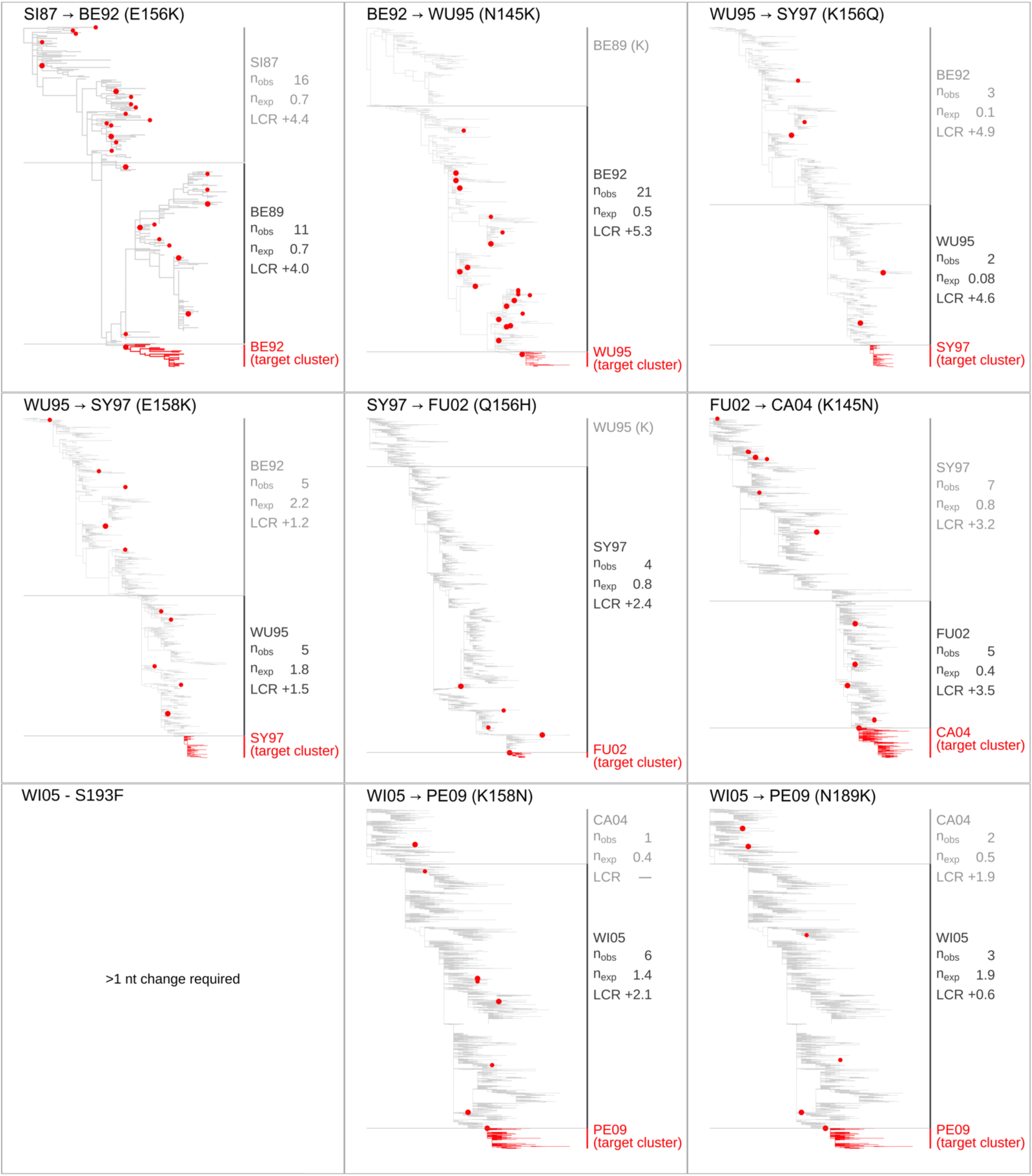
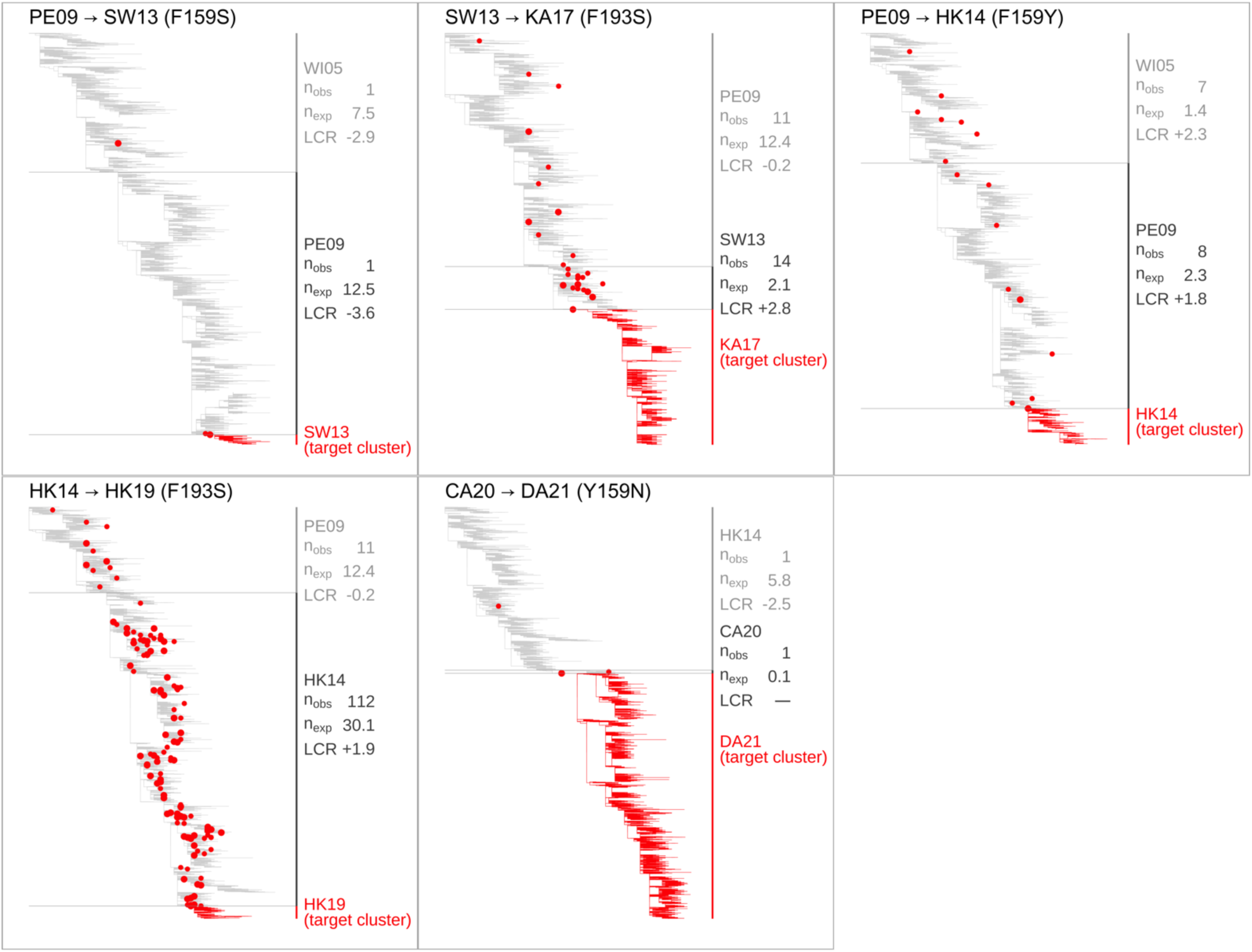
Trees marked with early occurrences of each cluster transition substitution, equivalent to Fig. 3A. For each antigenic cluster transition substitution, phylogenetic trees showing the previous two antigenic clusters are marked with occurrences of the cluster transition substitution. Larger points show occurrences on internal branches, and smaller points show occurrences on tips. Fig. 3A shows the F159Y substitution which caused the PE09 to HK14 antigenic cluster transition.

**Fig. S9.**
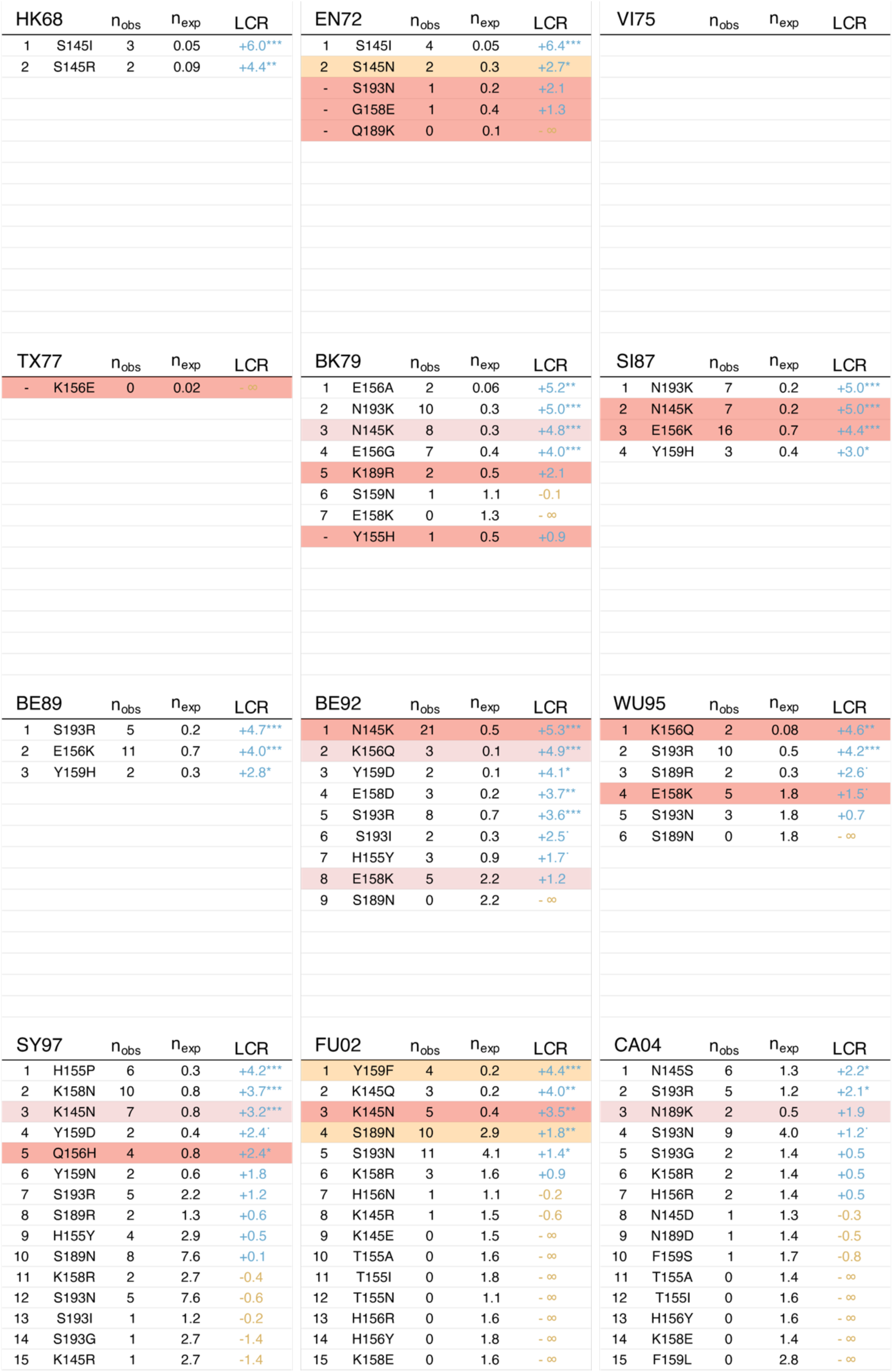
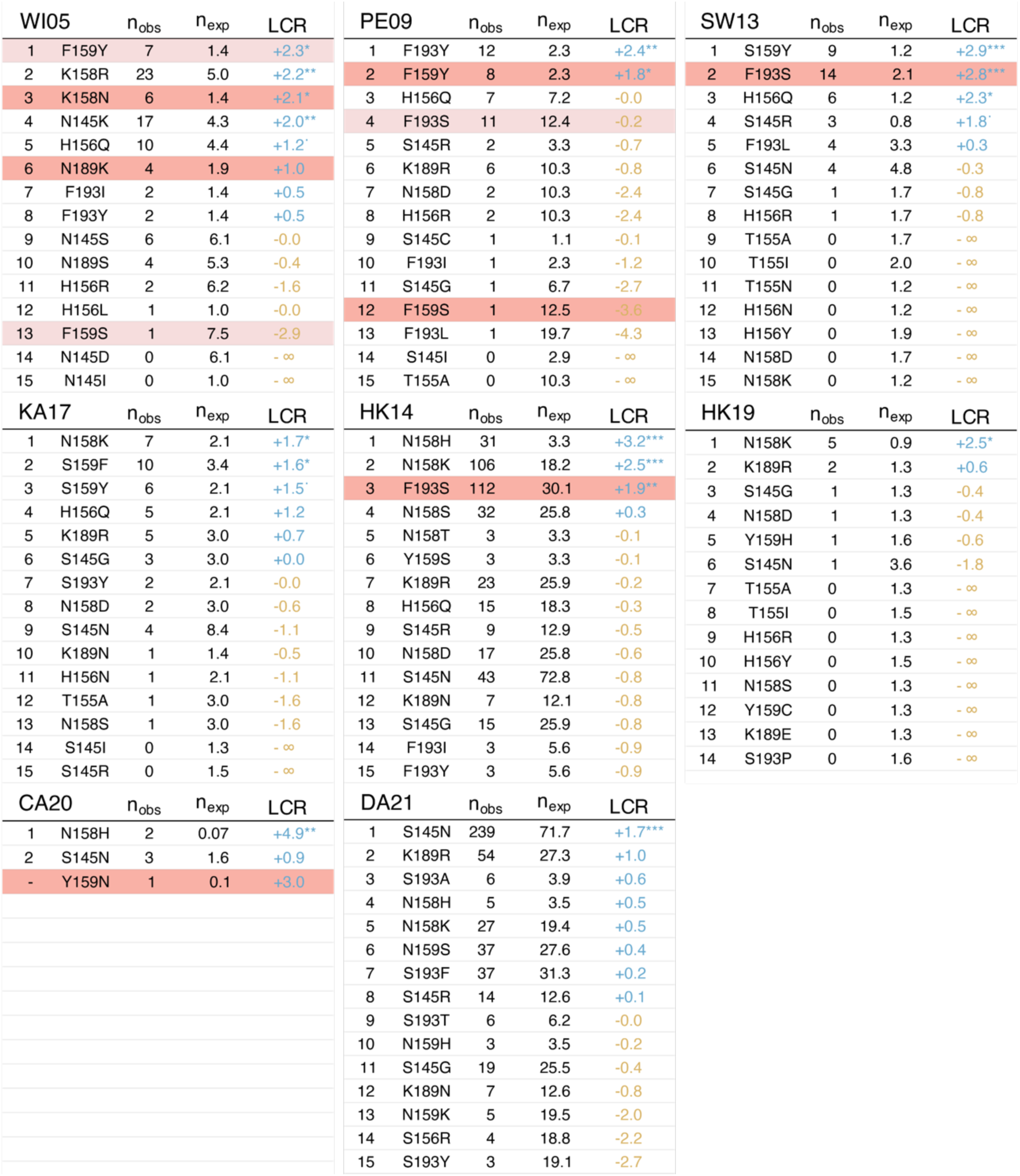
Rankings of substitutions at the seven key antigenic positions for each cluster, equivalent to Fig. 3C. Red highlighting indicates that the substitution causes an antigenic cluster transition from the current antigenic cluster; pale red highlighting indicates that the substitution causes an antigenic cluster transition from a descendant antigenic cluster; and orange highlighting indicates that the substitution occurs between the current antigenic cluster and a descendant antigenic cluster, but is not one of the causative antigenic cluster transition substitutions. ^.^ = p<0.1, * = p<0.05, ** = p<0.01, *** = p<0.001.

**Fig. S10.**
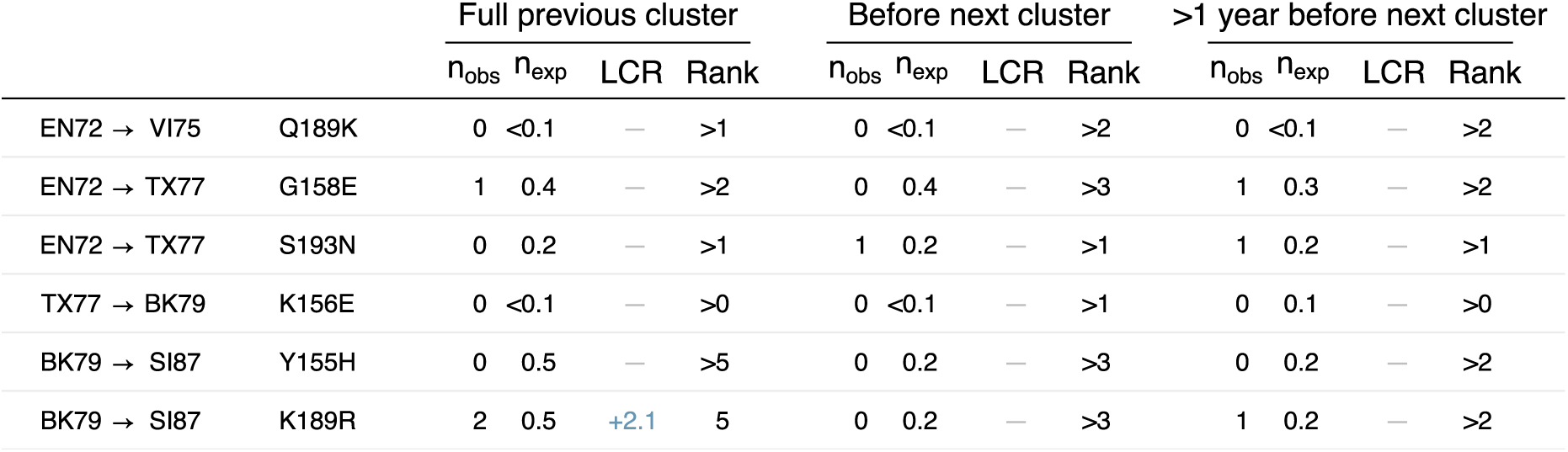
Log convergence ratio (LCR) and ranking of each cluster transition substitution from 1968 to 1987, equivalent to Fig. 4A. The “Full previous cluster” column uses all sequences in the ancestral cluster, regardless of isolation date; the “Before next cluster” and “>1 year before next cluster” columns use all sequences isolated 0-36 months or 12-48 months before the establishment of the descendant cluster respectively. n_obs_ and n_exp_ are the observed and neutral expected number of occurrences respectively. The HK68 to EN72 (T155Y) and BK79 to SI87 (S159Y) cluster transition substitutions are excluded as they required two nucleotide changes in the same codon. Fig. 4A shows antigenic cluster transition substitutions since 1987.

**Fig. S11.**
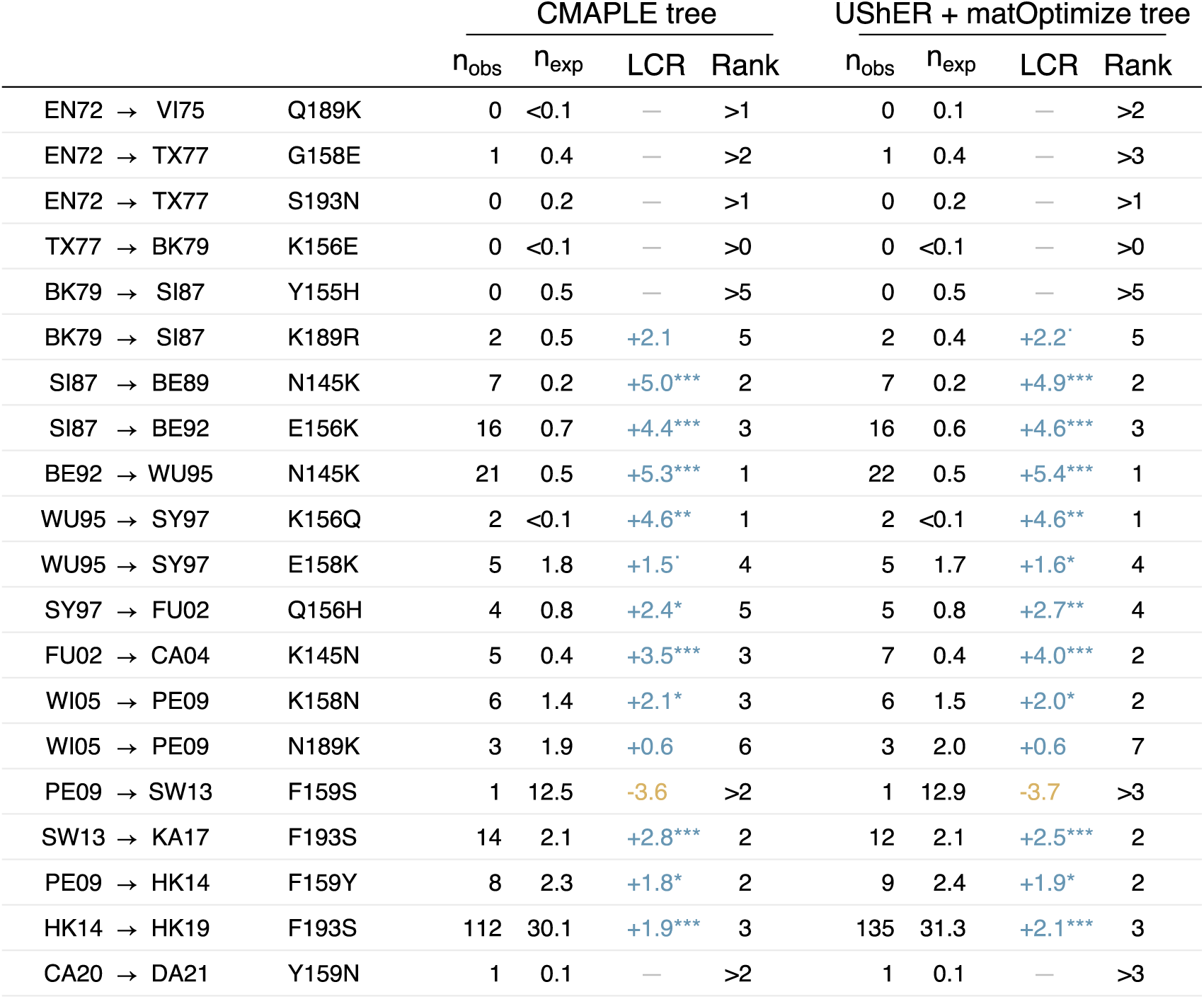
Rankings of cluster transition substitutions are concordant between trees produced using CMAPLE and UShER + matOptimize. The CMAPLE tree column shows the same data as Fig. 4A; equivalent ranks are also shown from a tree produced using UShER + matOptimize. The ranks differ by at most one position. Four cluster transition substitutions are excluded: HK68 to EN72 (T155Y), BK79 to SI87 (S159Y), and CA04 to WI05 (S193F) as each required two nucleotide changes in the same codon; and HK14 to CA20 (F193S), as F193S previously caused the transition from HK14 to HK19. ^.^ = p<0.1, * = p<0.05, ** = p<0.01, *** = p<0.001.

**Fig. S12.**
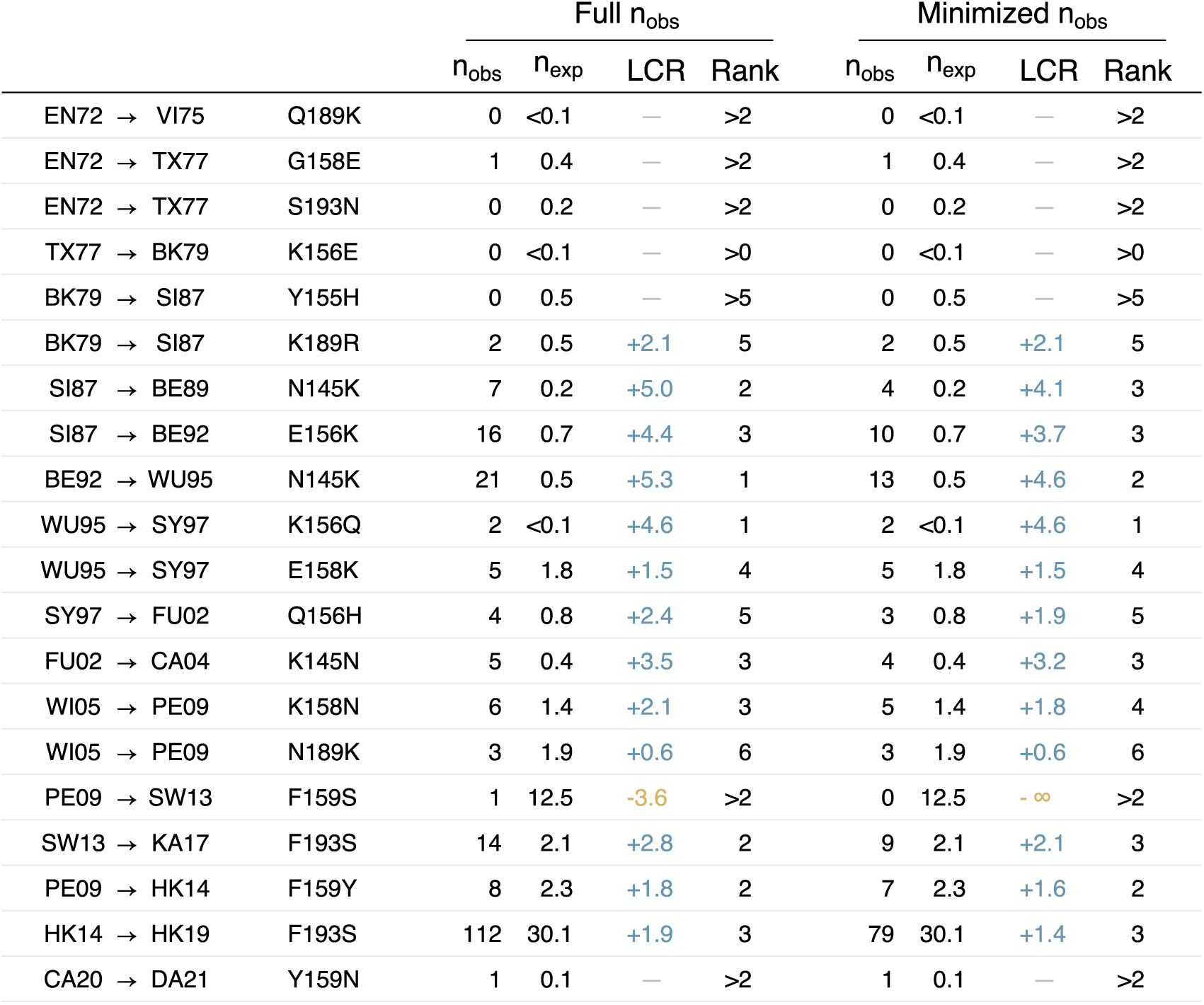
Antigenic cluster transition substitutions are highly ranked even when ambiguously placed occurrences are removed. The “Full n_obs_” column shows the same data as Fig. 4A; the “Minimized n_obs_” column shows ranks calculated using minimized values for n_obs_, where the minimum number of branches necessary to accommodate early occurrences of cluster transition substitutions was estimated by removing early occurrences with near-optimal placements outside of the ancestral cluster, and combining those with overlapping sets of near-optimal placements (see Methods section *Minimized counts of early occurrences* for more details). Four cluster transition substitutions are excluded: HK68 to EN72 (T155Y), BK79 to SI87 (S159Y), and CA04 to WI05 (S193F) as they required two nucleotide changes in the same codon; and HK14 to CA20 (F193S), as F193S previously caused the transition from HK14 to HK19.

**Fig. S13.**
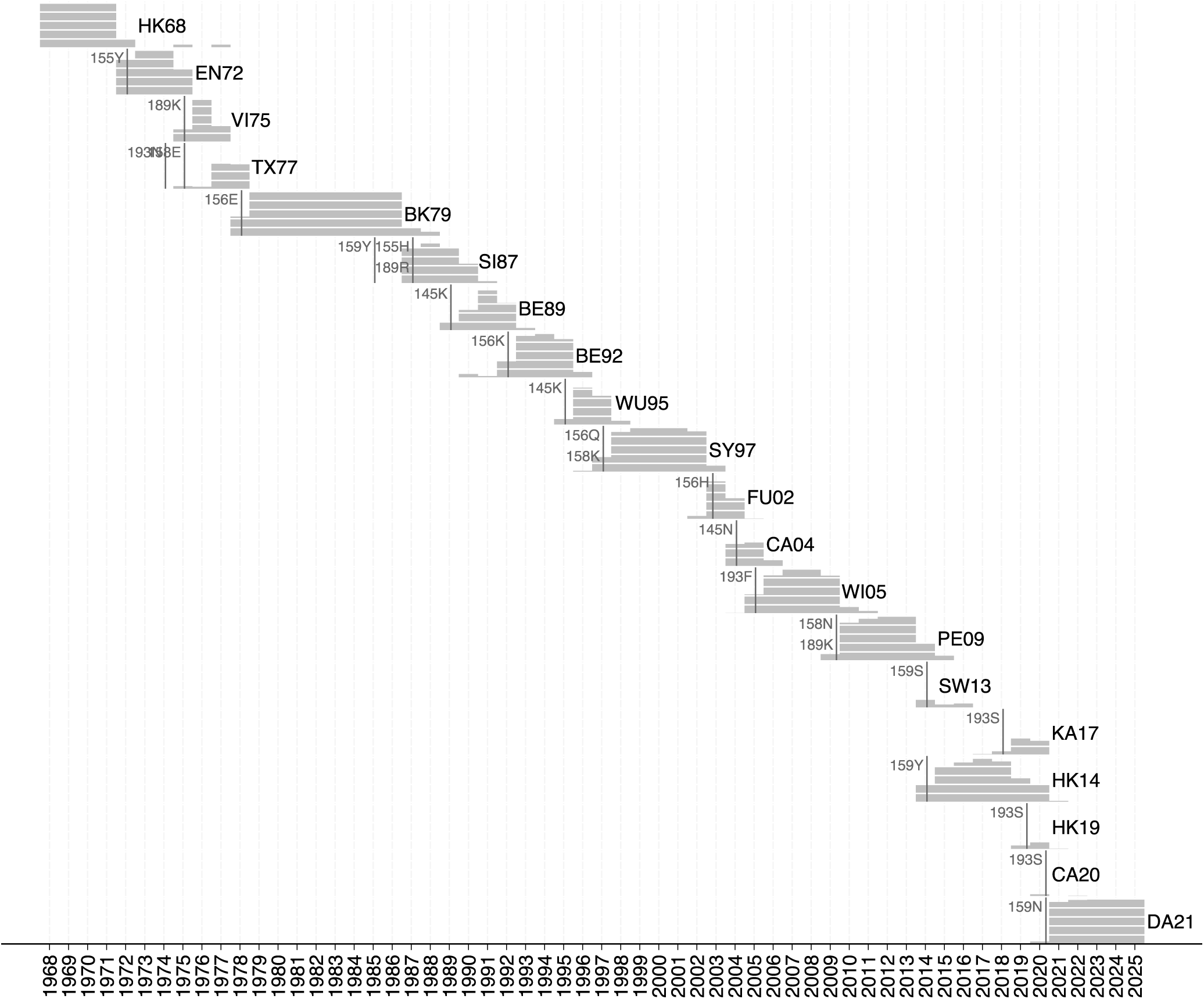
Relative circulation of antigenic clusters in each year from 1968 to 2025 in processed GISAID dataset. Each full grey bar delimits 20% of annual circulation. The establishment dates for each cluster transition substitution, which are used to assess whether positive selection for the cluster transition substitution was detectable before there was substantial circulation of the substitution, are shown with vertical grey lines. See Methods section *Constructing temporally early trees* for more detail, and Fig. S14.

**Fig. S14.**
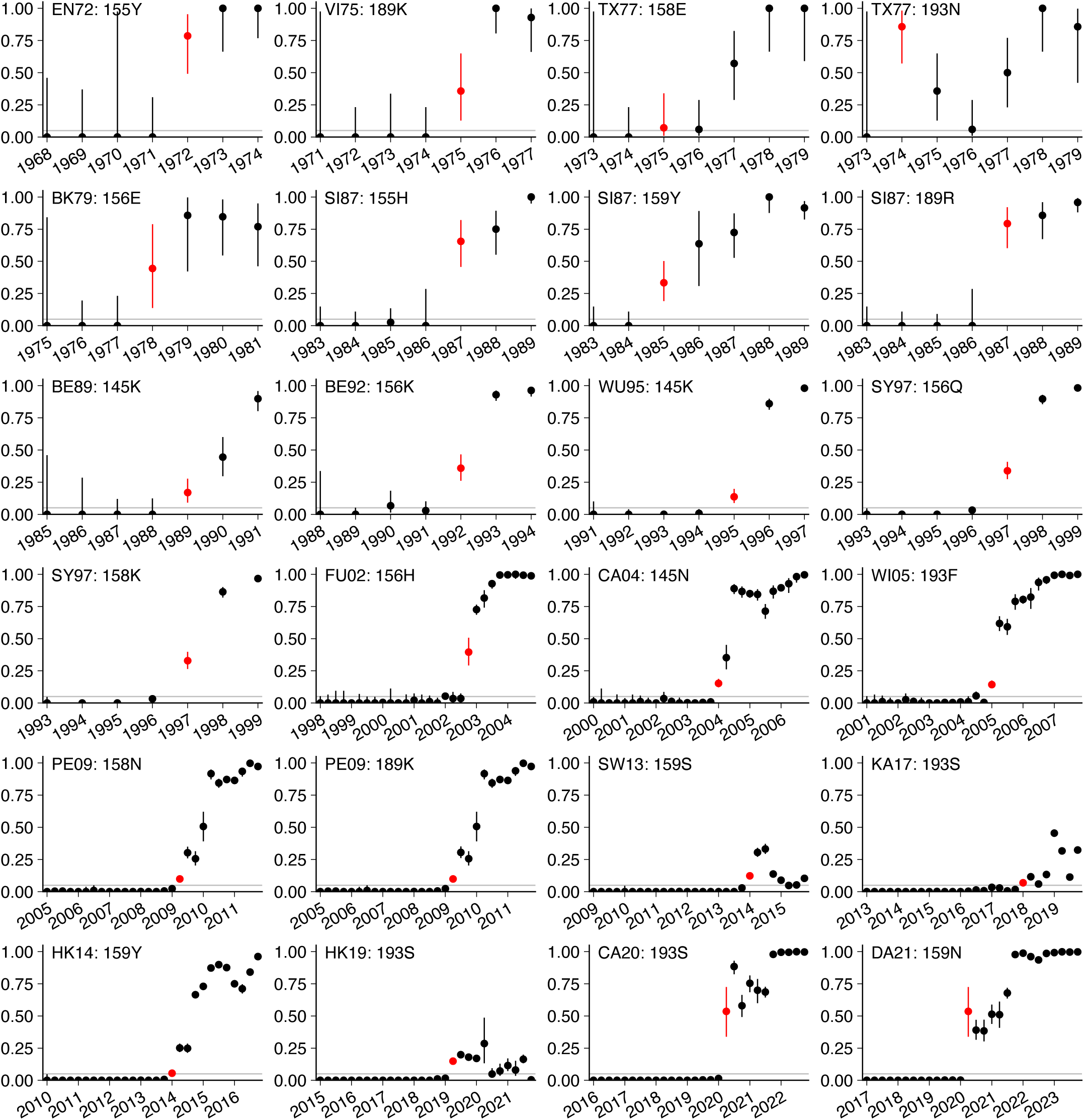
Estimated establishment dates for antigenic cluster transition substitution. The establishment date (indicated by a red point) for each antigenic cluster transition substitution is the beginning of the first sequence of two consecutive timepoints (yearly for earlier clusters, quarterly for later clusters) where the cluster transition substitution is at >5% frequency (grey horizontal line). See Methods section *Constructing temporally early trees* for more detail.

**Fig. S15.**
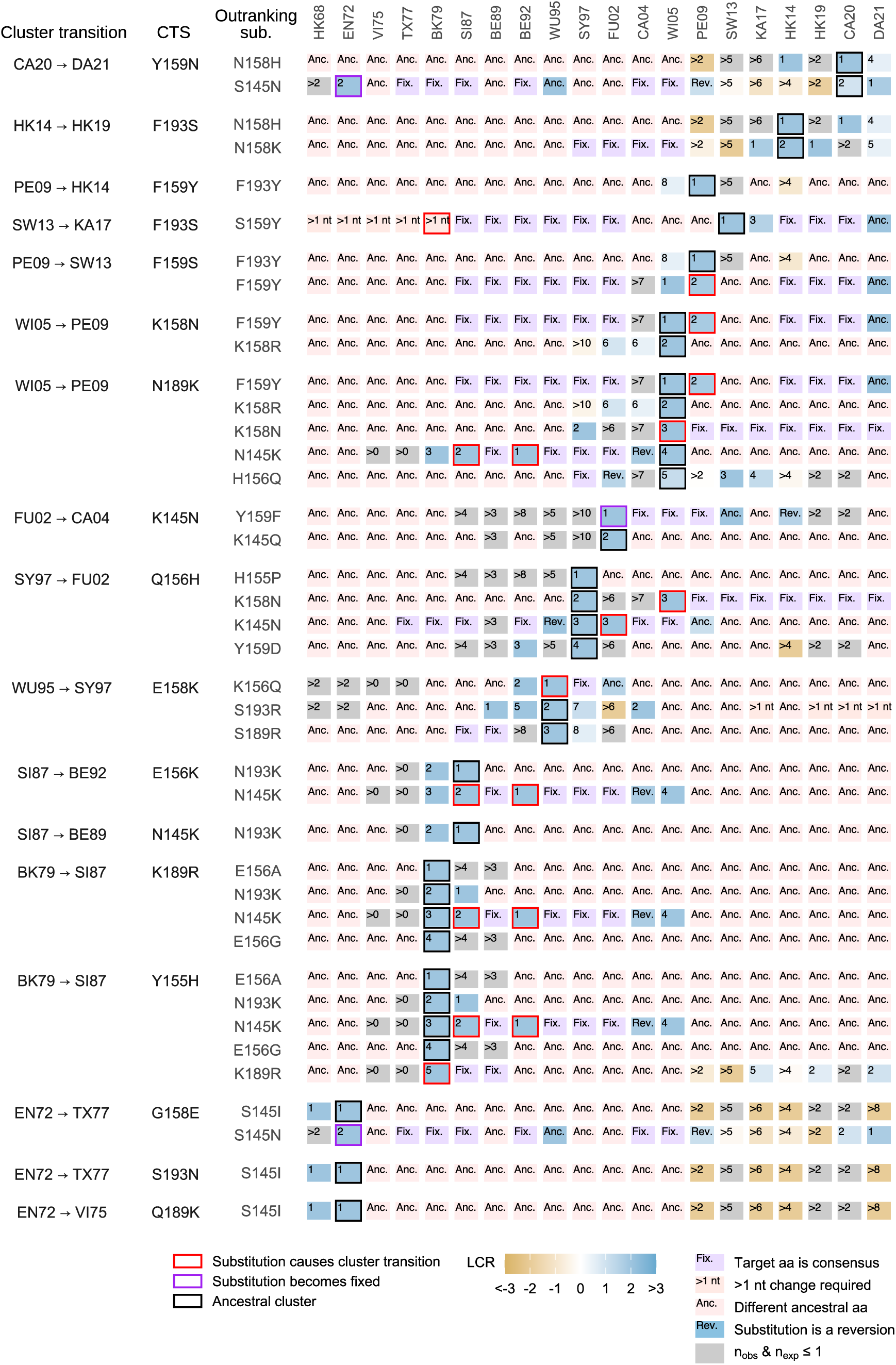
History and fate of substitutions which “outrank” antigenic cluster transition substitutions. For each cluster transition substitution (CTS), the log convergence ratio (LCR) and ranking of the substitutions which are ranked above it in the cluster before it caused a cluster transition (“outranking” substitutions) are shown. Clusters where the outranking substitution is in the consensus sequence, where an amino acid other than the ancestral amino acid of the outranking substitution is in the consensus sequence, where the outranking substitution is not possible via a single nucleotide mutation, or where the outranking substitution is positively selected but is a reversion to the previous cluster are indicated.

**Fig. S16.**
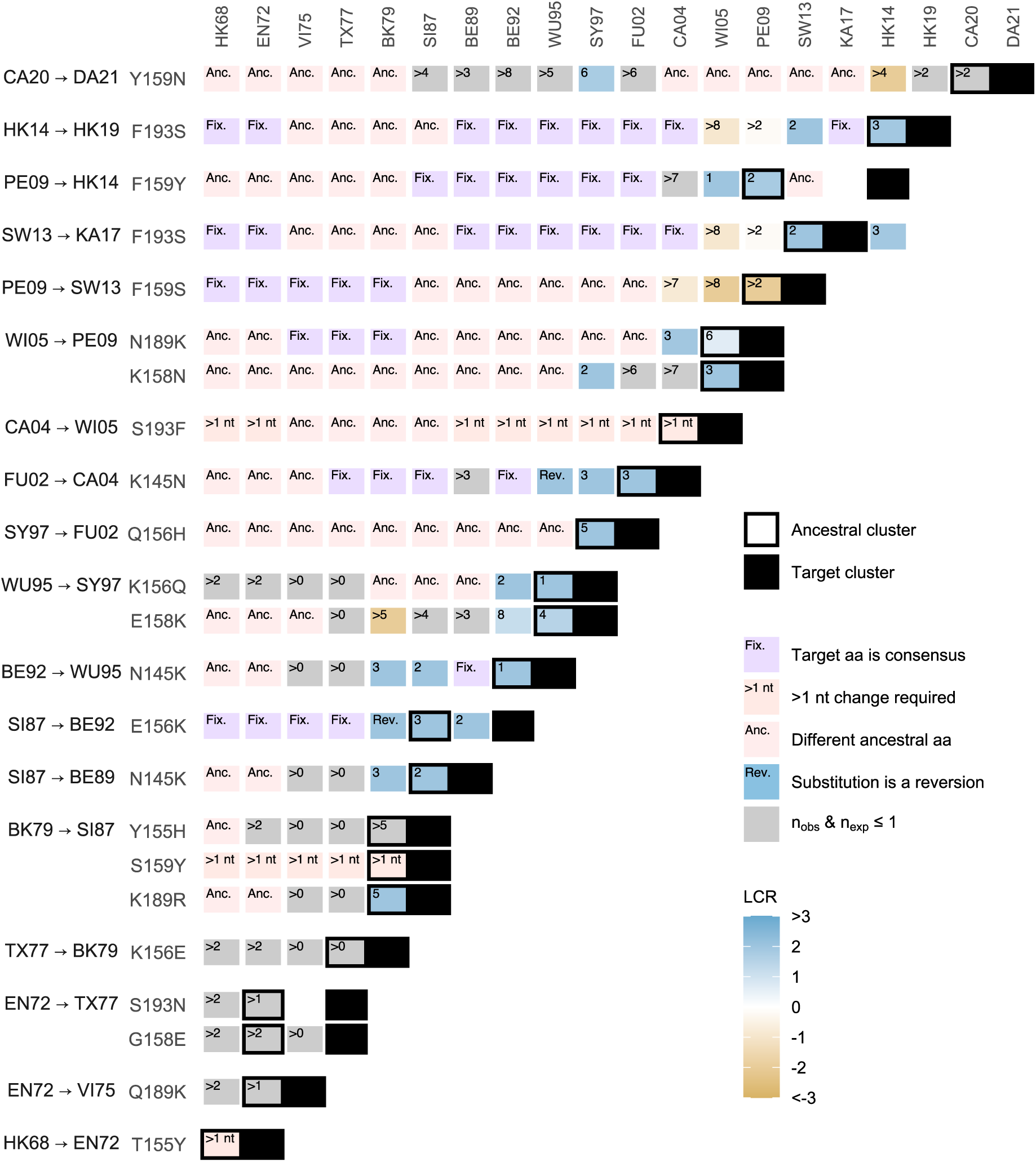
Log convergence ratios (LCR) and rankings for all cluster transition substitutions in all previous clusters. Numbers indicate the rank of each substitution in each cluster. The phylogenetically ancestral cluster is indicated with a black outline. Cases where the cluster transition substitution is already in the consensus sequence for a cluster, where an amino acid other than the ancestral amino acid of the cluster transition substitution is in the consensus sequence, where the cluster transition substitution is not possible as a single nucleotide change, and where the cluster transition substitution is positively selected but is a reversion to the previous cluster are indicated.

**Fig. S17.**
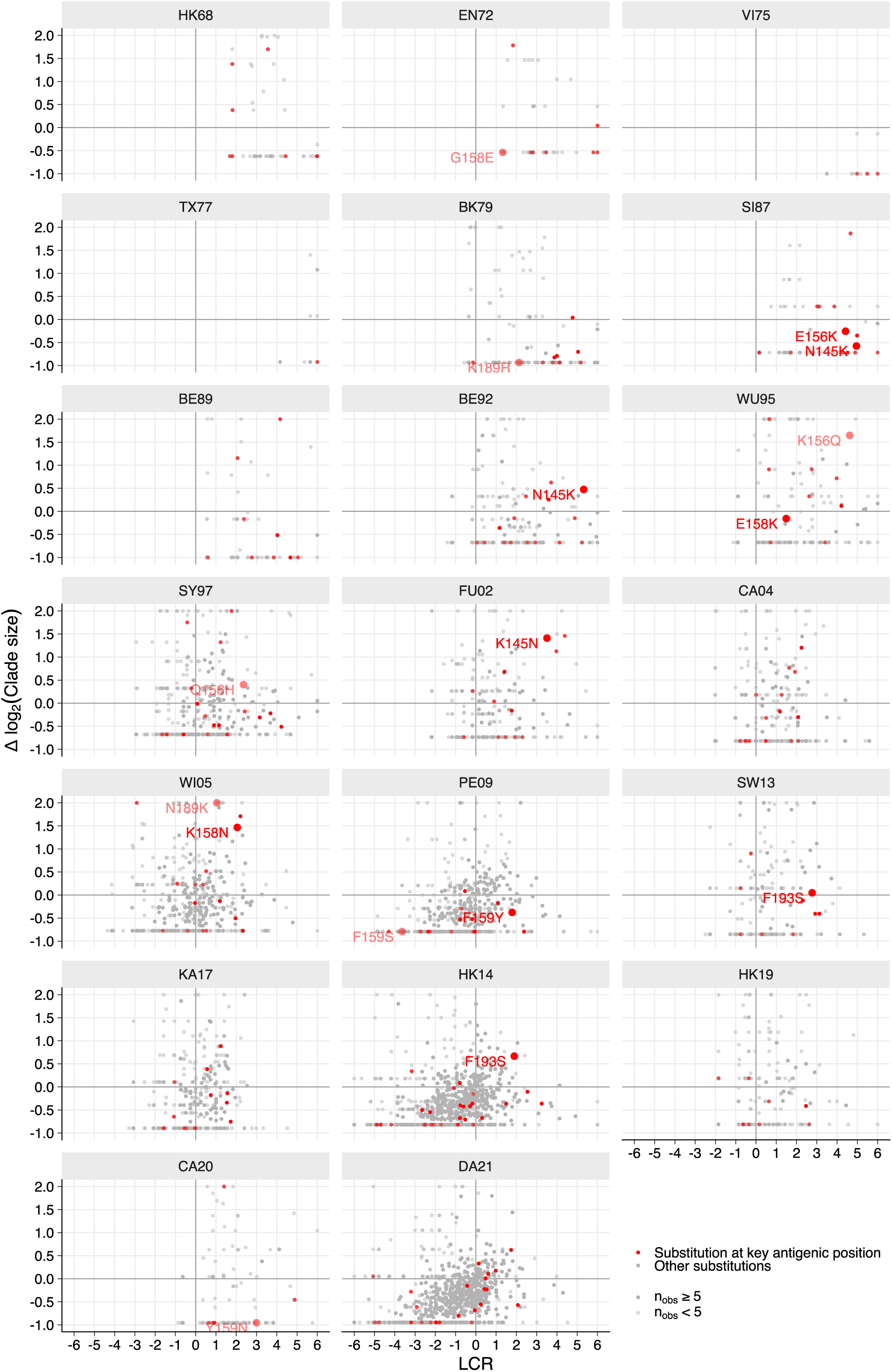
Mean clade size is correlated with the log convergence ratio (LCR), but is a poorer predictor of future antigenic substitutions. For each amino acid substitution, the y-axis position shows the difference between the mean log_2_ number of descendants for occurrences of the substitution compared to occurrences of synonymous nucleotide mutations. Antigenic cluster transition substitutions from each cluster are labelled. While clade size is significantly correlated with LCR (Fig. 4C), there are seven occasions where a cluster transition substitution has a positive LCR, indicating positive selection, but the mean clade size is smaller than that for synonymous mutations, indicating negative selection (with n_obs_ ≥ 5: SI87 to BE89, N145K; SI87 to BE92, E156K; WU95 to SY97, E158K; PE09 to HK14, F159Y; with n_obs_ < 5: EN72 to TX77, G158E; BK79 to SI87, K189R; CA20 to DA21, Y159N).

**Fig. S18.**
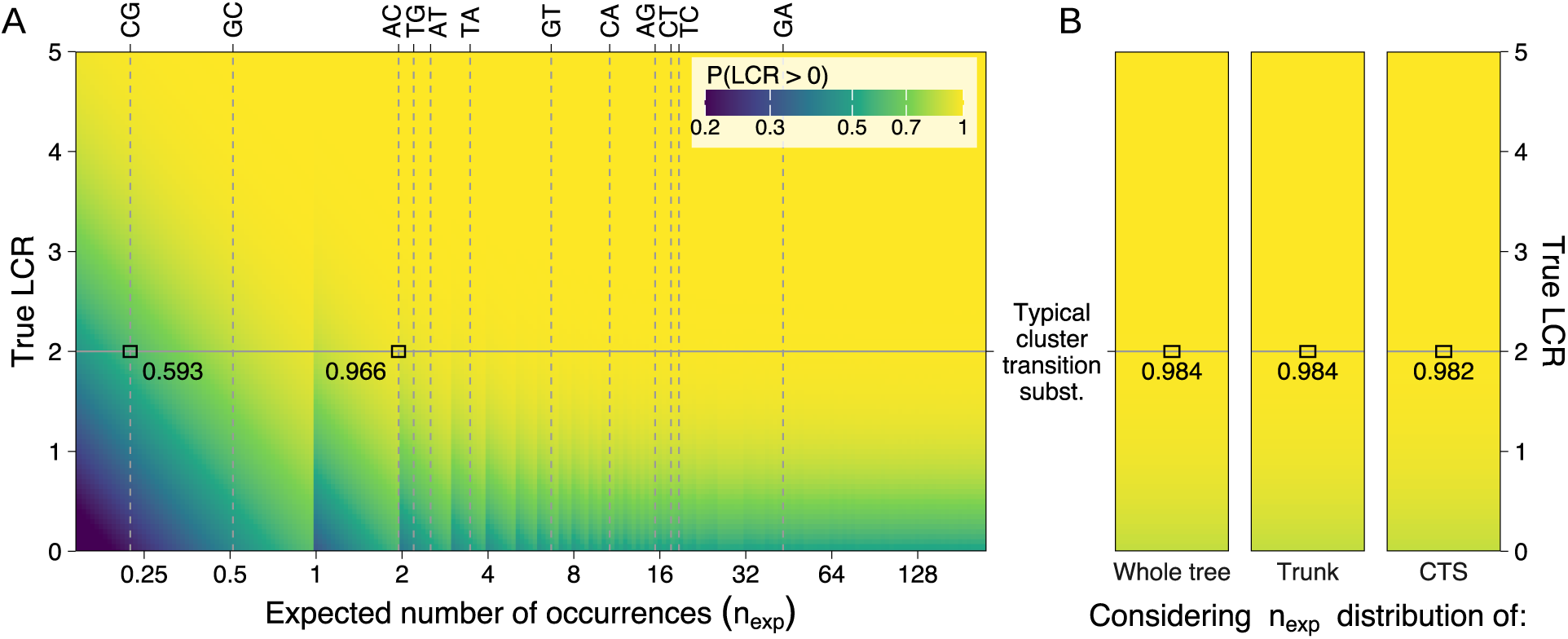
Equivalent to Fig. 5B, C, except coloring represents the probability of obtaining a positive value for the LCR, rather than the median p-value. (**A**) The probability of obtaining a positive LCR as a function of the neutral expected number of occurrences and true log convergence ratio (LCR). Vertical dashed lines show neutral expected occurrences for each mutation in 40,000 sequences. Values referenced in the text are highlighted. (**B**) As **A**, but considering the n_exp_ distribution (Fig. S19) for substitution observed anywhere on the tree, those observed on the trunk, and cluster transition substitutions. See Methods section *Simulating LCRs for a hypothetical cluster transition substitution*.

**Fig. S19.**
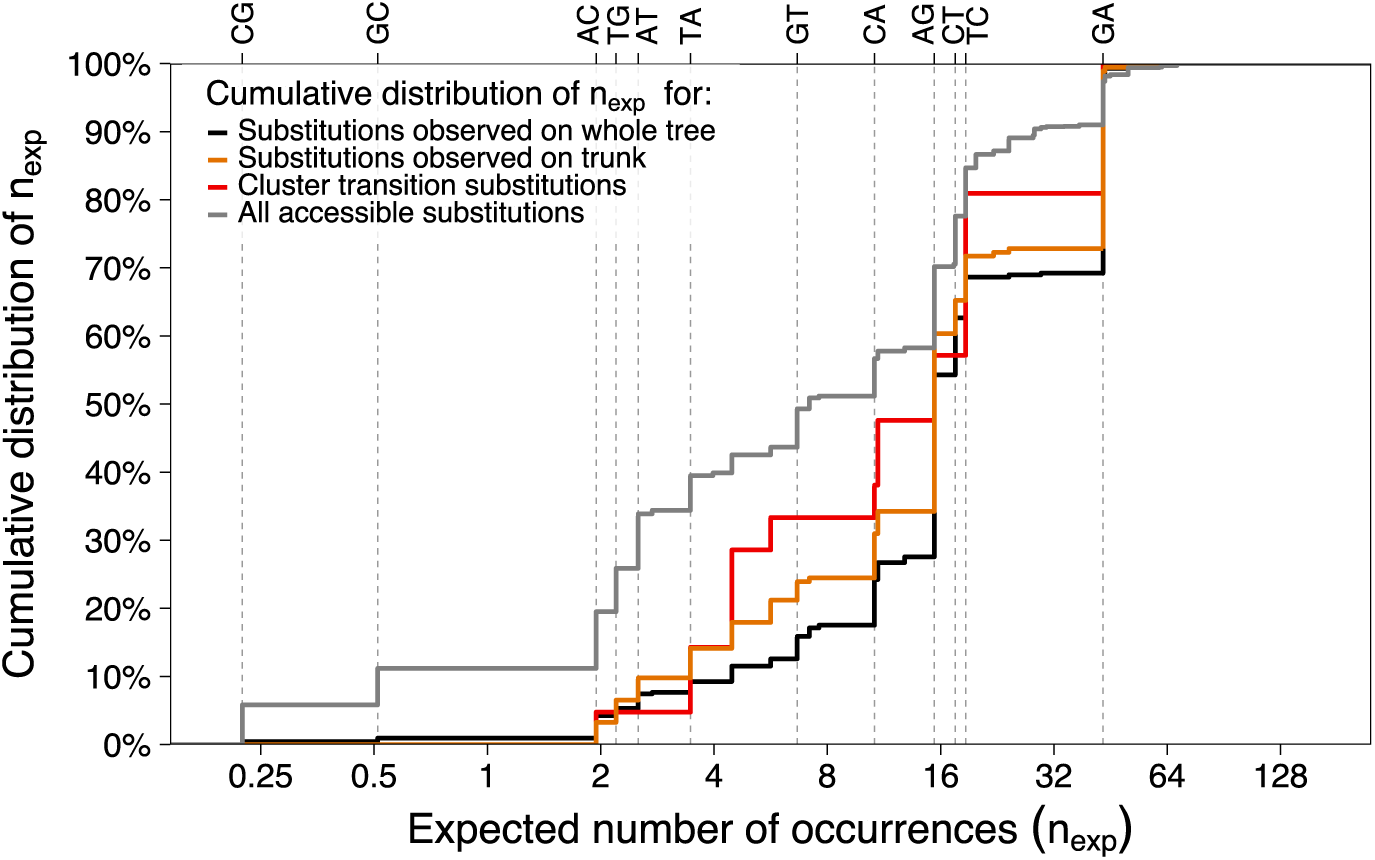
Distributions of neutral expected occurrence counts used in Fig. 5C & S18B. Empirical cumulative distribution functions for the expected number of occurrences in a hypothetical 40,000 sequence cluster, for substitutions observed anywhere on the tree (black), on the trunk of the tree (orange), or causing antigenic cluster transitions (red); and for all substitutions accessible by a single nucleotide change in HA1 (grey line, not used in Fig. 5C or S18B). Vertical dashed lines indicate the expected number of occurrences for a substitution caused by each nucleotide mutation. See Methods section *n_exp_ distributions for different substitution types* for details.

**Fig. S20.**
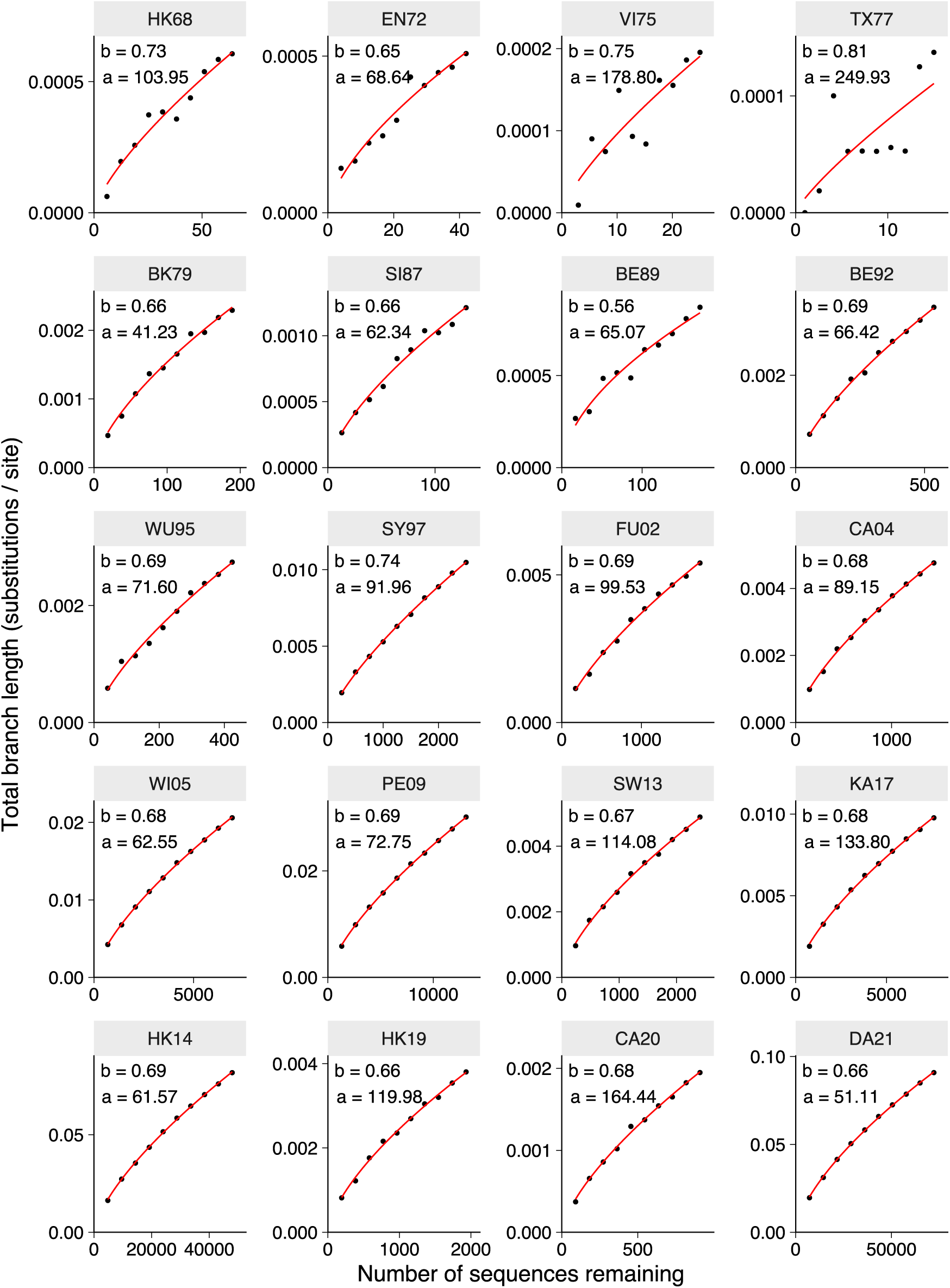
Increasing sequencing intensity gives diminishing returns on the resolution of LCR estimates. The effect of sequencing intensity on the total branch length of the tree (which is closely related to the expected number of occurrences) was determined by artificially varying the sequencing intensity by discarding between 0% and 90% of the sequences in each cluster. *a* and *b* represent parameters of a power-law model of the effect of sequencing intensity on the total branch length of the tree, which was fit to the data from each cluster (branch length = number of sequences^b^ / a). See Methods section *Modelling the effect of sequencing intensity on n_exp_* for details.

**Fig. S21.**
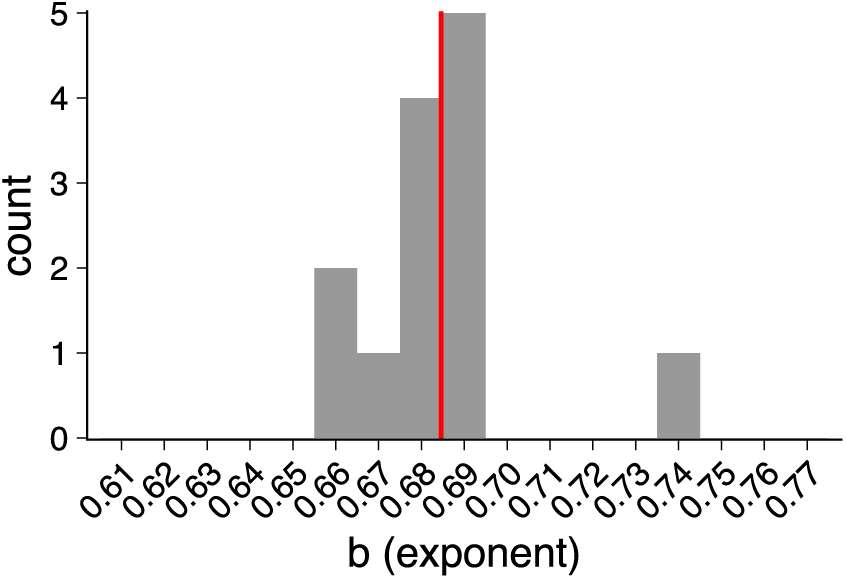
Branch length increases with the number of sequences raised to the power ∼0.69. Distribution of the exponent *b* of the sequencing intensity model (branch length = number of sequences^b^ / a) for clusters containing at least 200 sequences. The vertical red line shows the mean value over these clusters. See Methods section *Modelling the effect of sequencing intensity on n_exp_* for details, and Fig. S20 for model fits.

**Fig. S22.**
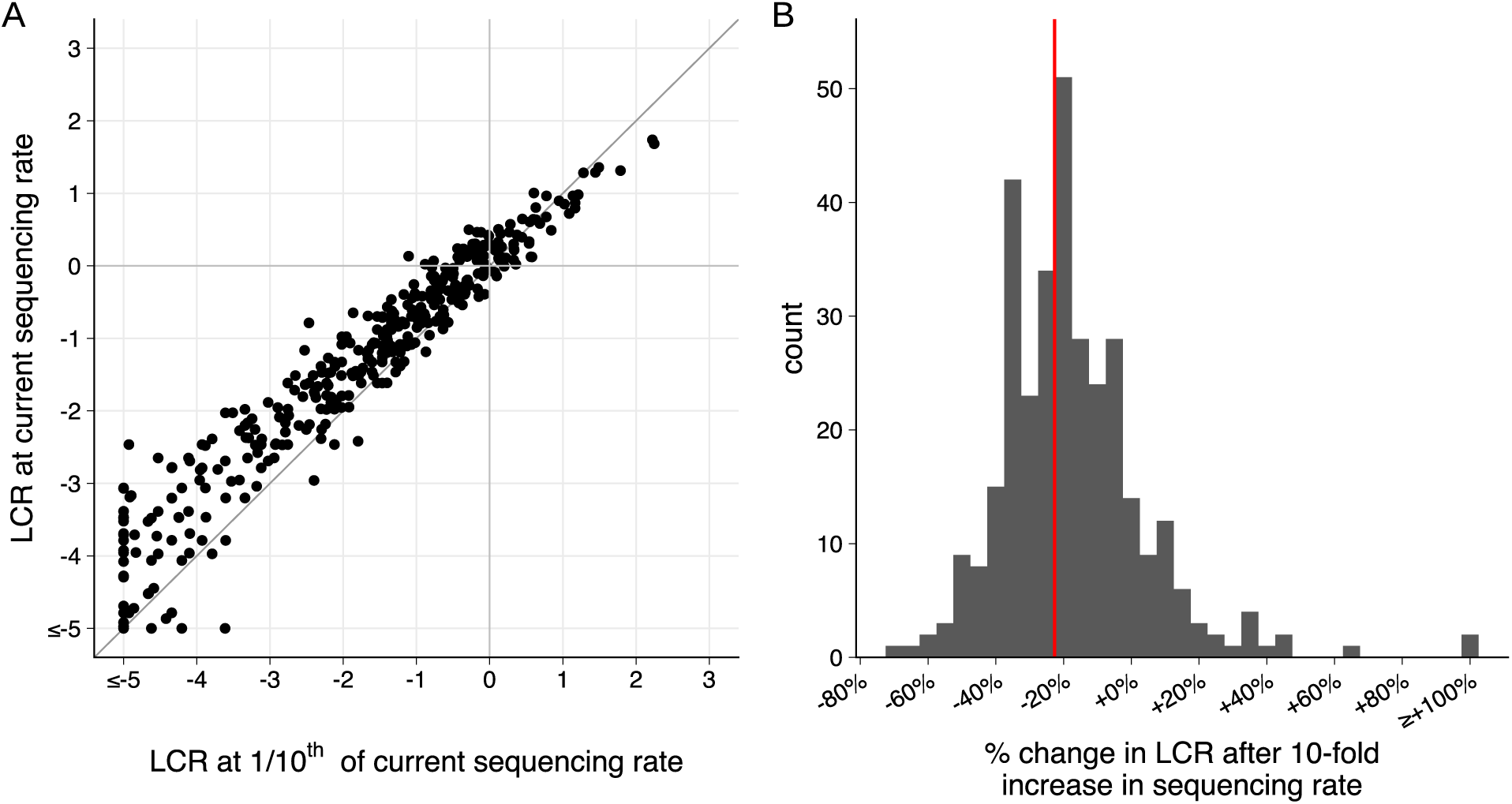
A 10-fold change in sequencing intensity changes the magnitude of log convergence ratio (LCR) estimates by ∼20%. We calculated LCRs for a subtree containing all DA21 sequences, and for 10 subtrees containing randomly selected 10% subsets of the DA21 sequences. Observed and expected mutation counts are averaged over the 10 subsetted trees for each substitution. Substitutions are shown in the scatter plot (**A**) if the average expected number of occurrences is at least five across the subsetted trees, and there is at least one observed occurrence of the substitution in any of the subsetted trees. The histogram (**B**) additionally filters to substitutions with LCR > 0.5 in the subsetted data. The median change in LCR after the 10-fold increase in sequencing intensity between the subsetted and the full sequence sets is shown with the red vertical line.

**Fig. S23.**
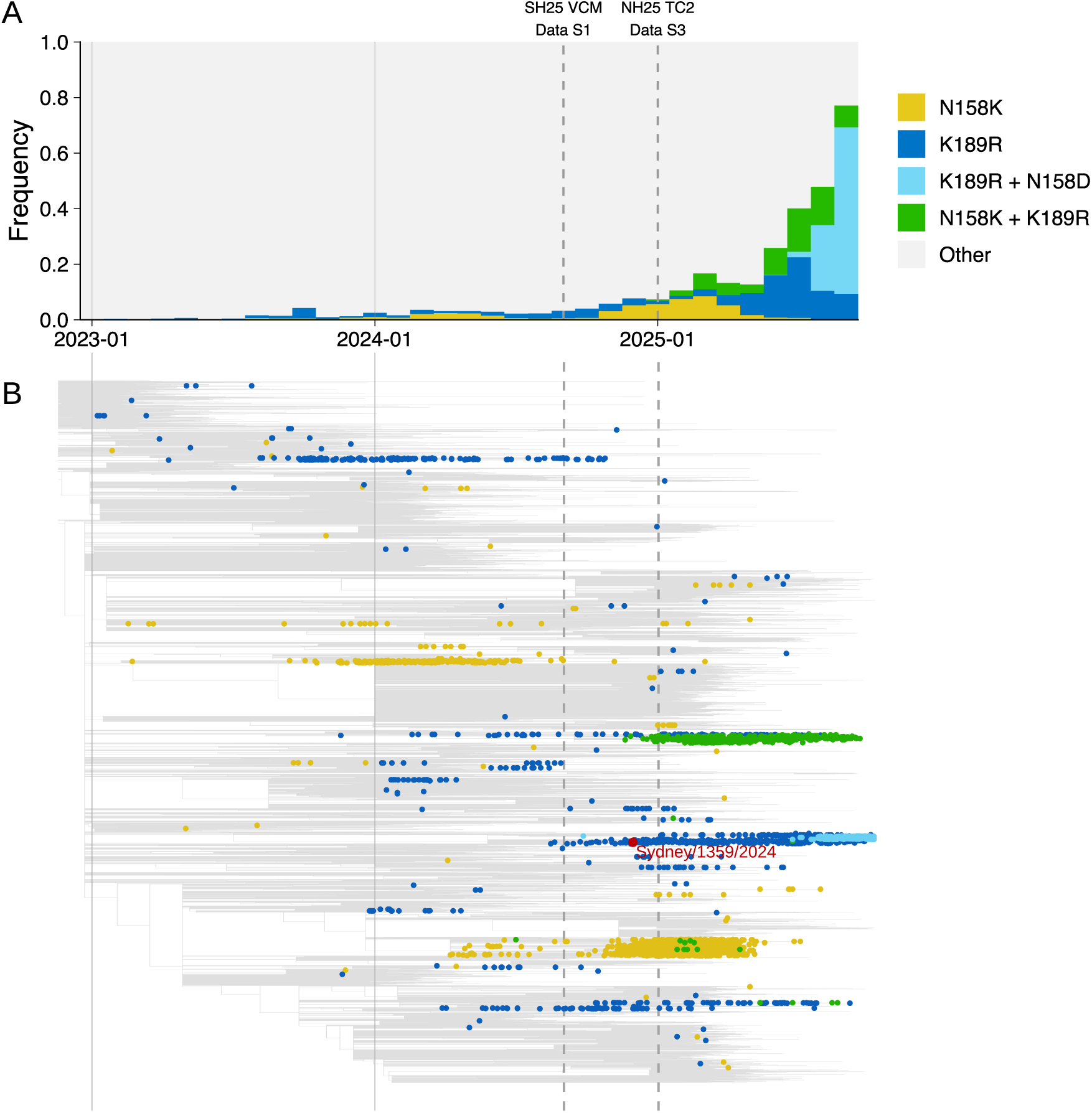
Clades containing N158K and K189R have emerged and grown since their identification based on convergent evolution in September 2024. (**A**) Global proportion of sequences containing N158K, K189R, N158K+K189R, or N158D+K189R in GISAID. (**B**) Time-resolved phylogenetic tree of sequences in GISAID collected since 1^st^ January 2023, with those containing substitution combinations from **A** marked. Approximate dates are indicated for the WHO Vaccine Composition Meeting for the Southern Hemisphere 2025, where N158K and K189R were first identified, and for Teleconference 2 prior to the WHO Vaccine Composition Meeting for the Northern Hemisphere 2025/26, where the N158K+K189R double mutant clade was identified. The vaccine virus recommended following the WHO Vaccine Composition Meeting for the Southern Hemisphere 2026 (A/Sydney/1358/2024 virus) is marked on the phylogenetic tree.

**Fig. S24.**
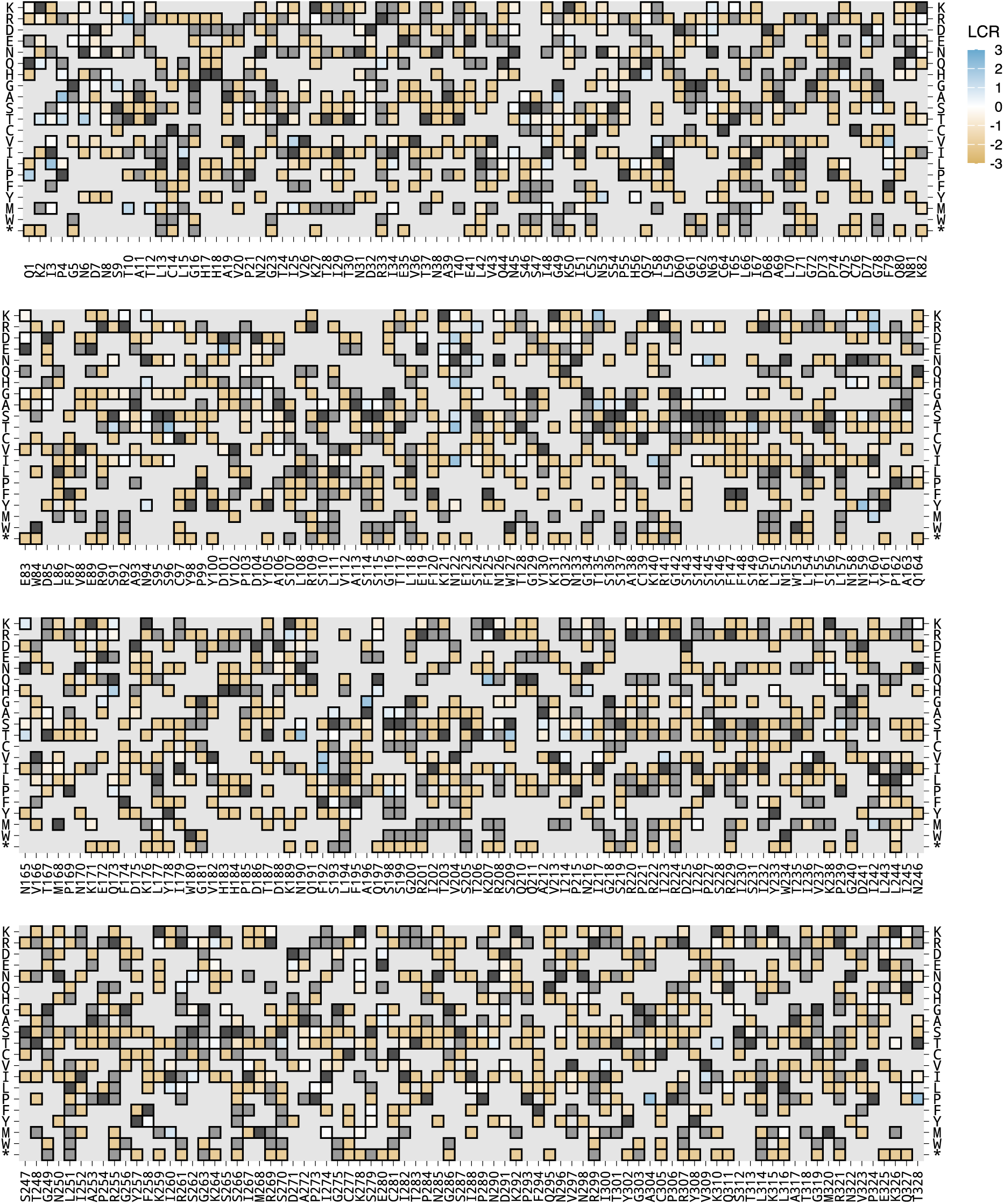
Estimates of the log convergence ratio (LCR) for all amino acid substitutions at HA positions 1-328 in the DA21 antigenic cluster. Dark grey squares identify the ancestral amino acid at the position, and light grey squares show substitutions where neither the expected nor observed number of occurrences exceed one.

**Data S1 (separate file)**

Convergence data compiled in September 2024 for the WHO Consultation on the Composition of Influenza Vaccines for the Southern Hemisphere 2025.

**Data S2 (separate file)**

Convergence data compiled in December 2024 for the first Teleconference prior to the WHO Consultation on the Composition of Influenza Vaccines for the Northern Hemisphere 2025/26.

**Data S3 (separate file)**

Convergence data compiled in January 2025 for the second Teleconference prior to the WHO Consultation on the Composition of Influenza Vaccines for the Northern Hemisphere 2025/26.

