## Supplementary material for "Predicting the antigenic evolution of seasonal influenza viruses using phylogenetic convergence": Data S1

### Information for the WHO Consultation on the Composition of Influenza Vaccines for the Southern Hemisphere 2025

Addendum 3 (H3 convergent evolution)

23 September 2024

Center for Pathogen Evolution

University of Cambridge, United Kingdom

### Convergent evolution of F193S

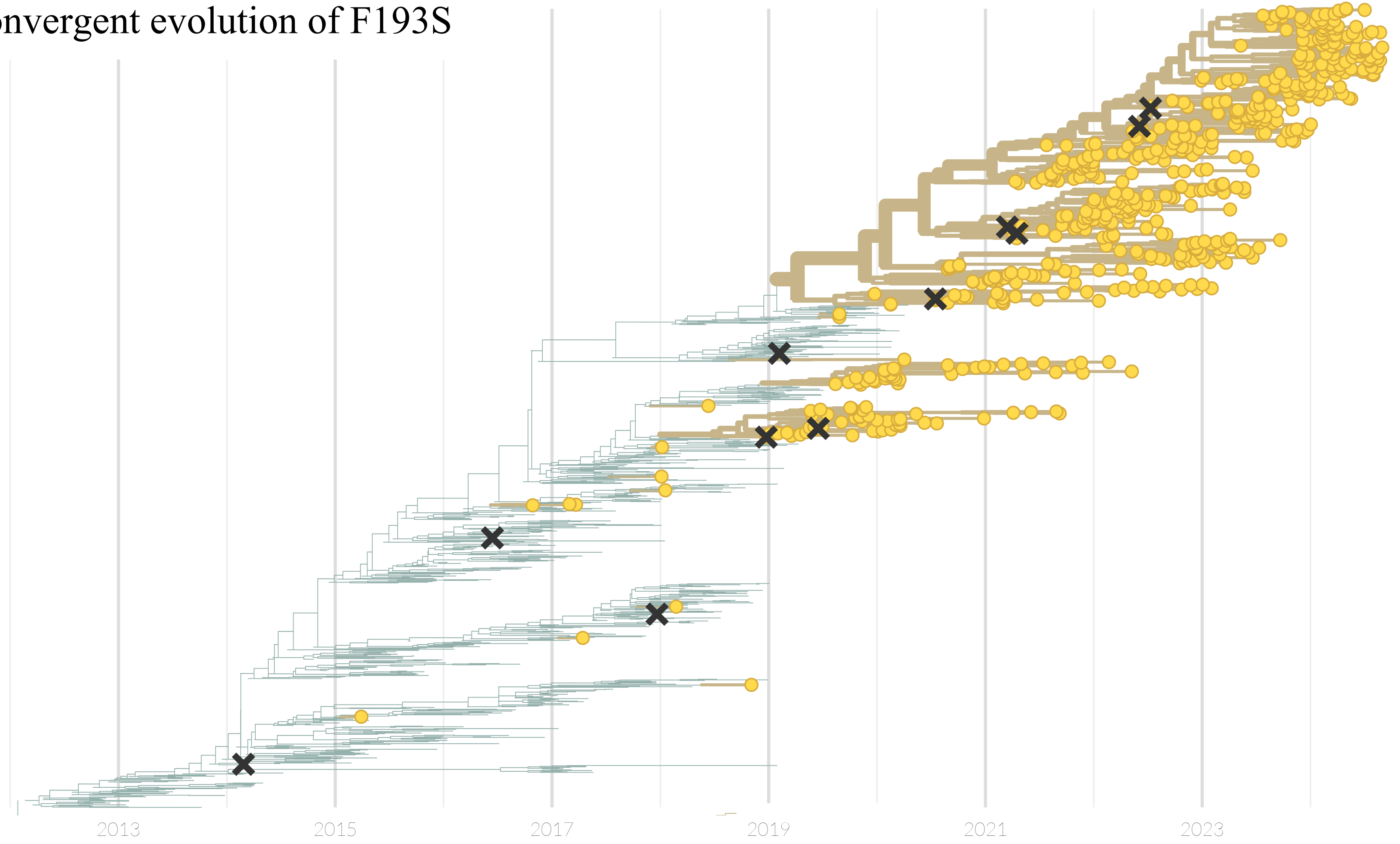

### Convergent evolution of F193S

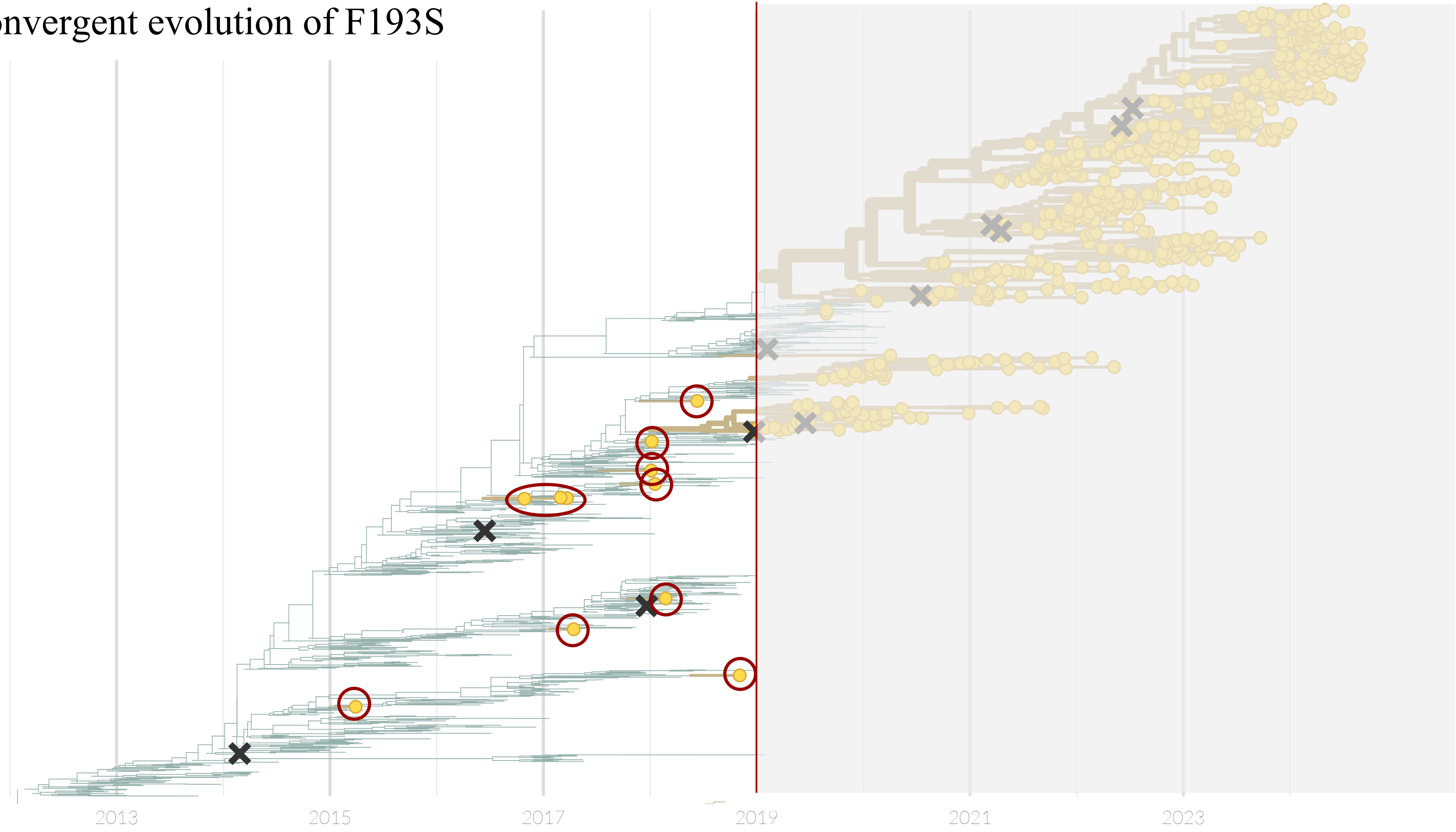

### Convergent evolution of F193S

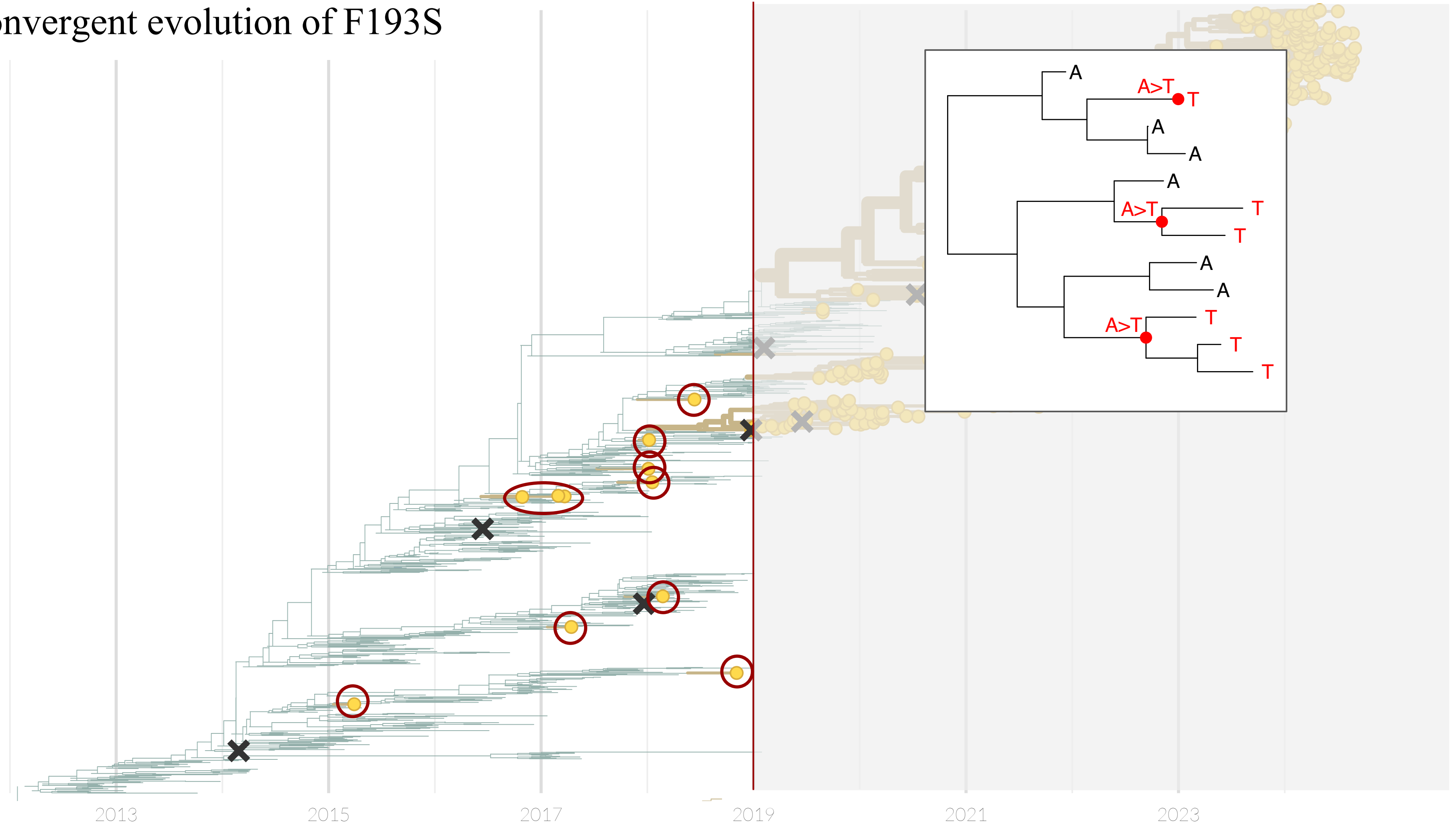

### Convergent evolution of F193S

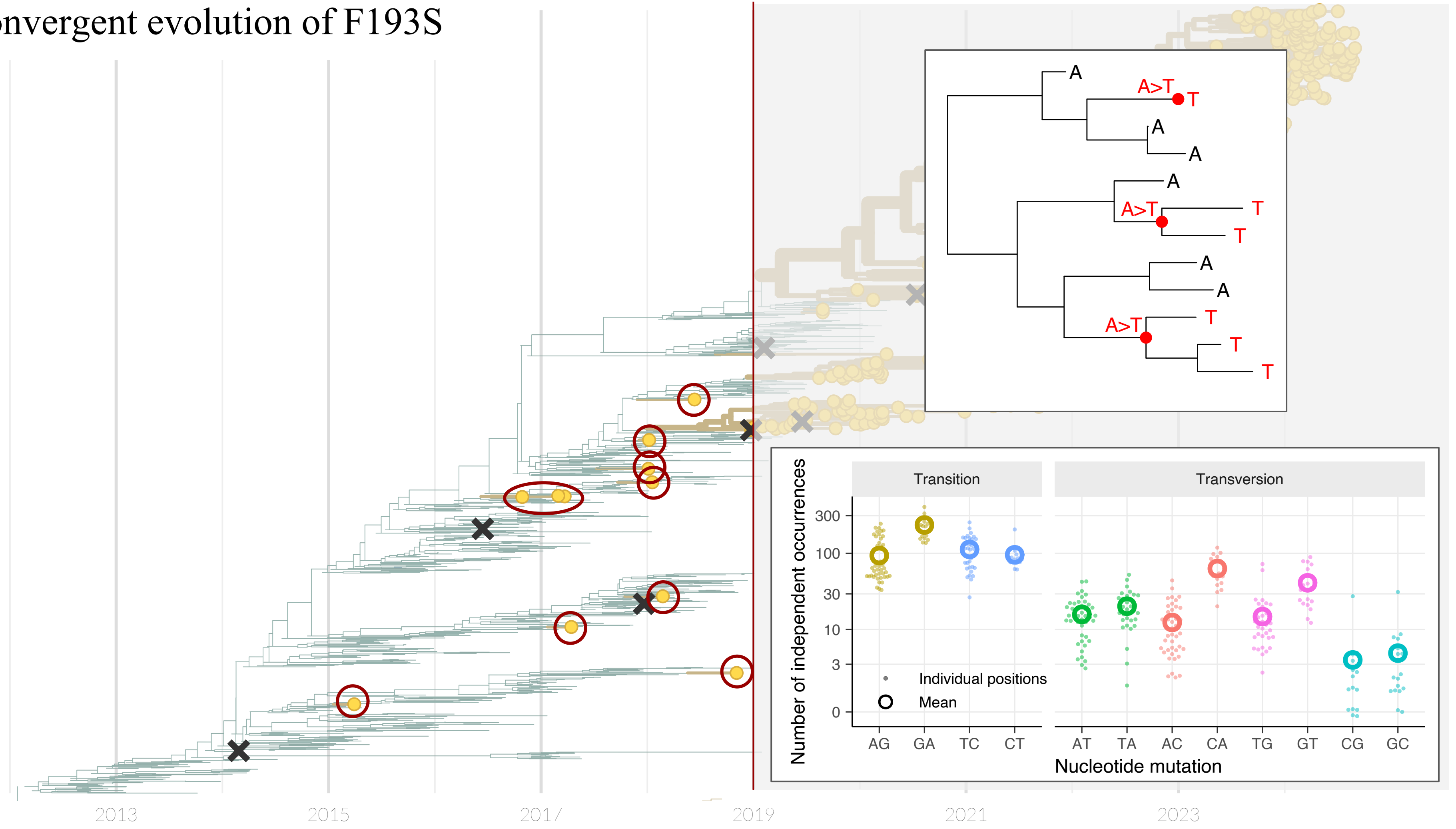

### Fitness effect (FE) measurements

(here in Hong Kong/4801/2014-like viruses)

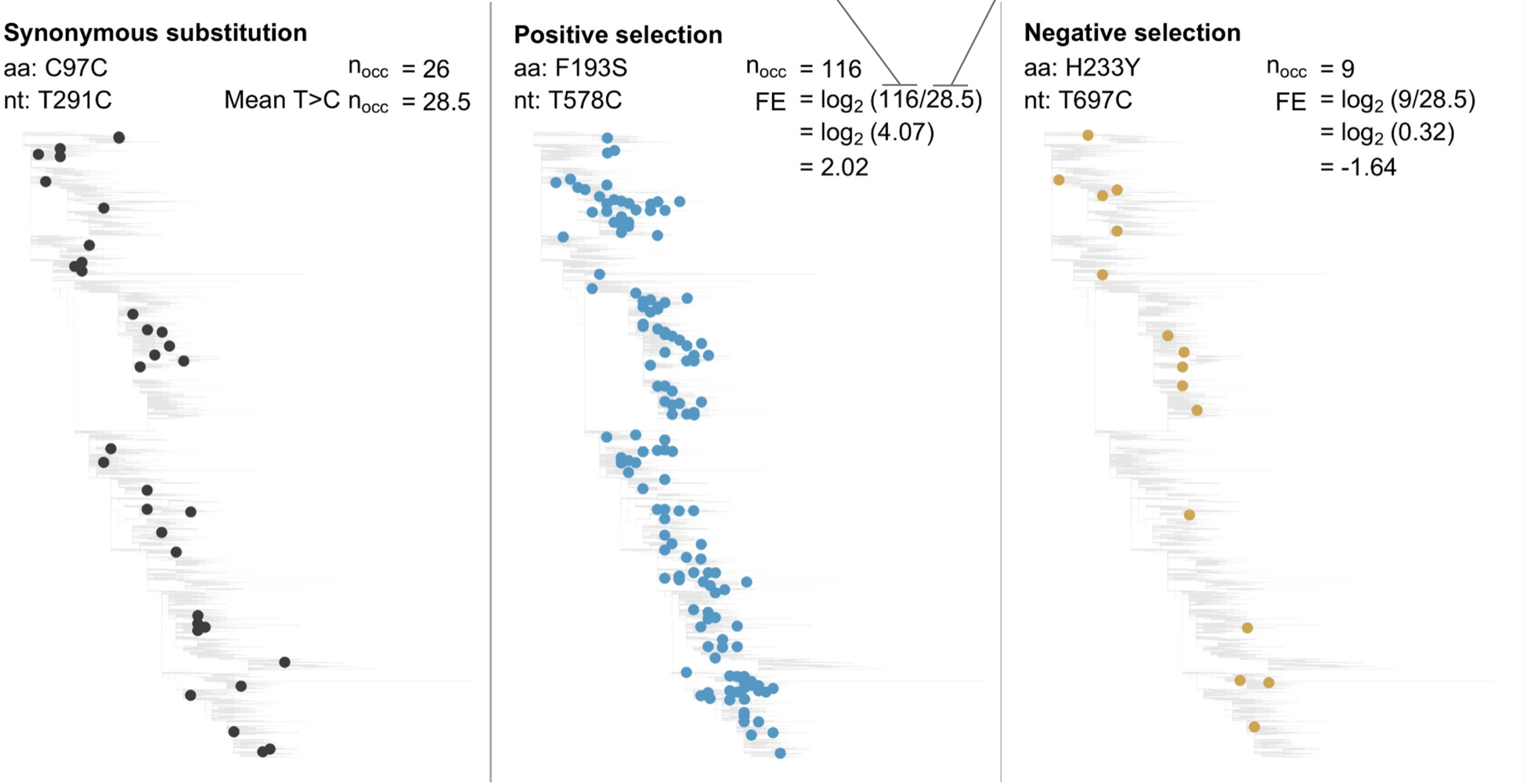


Similar to Bloom & Neher 2023 on SARS-CoV-2 https://doi.org/10.1093/ve/vead055

### Fitness effect (FE) measurements

(here in Hong Kong/4801/2014-like viruses)

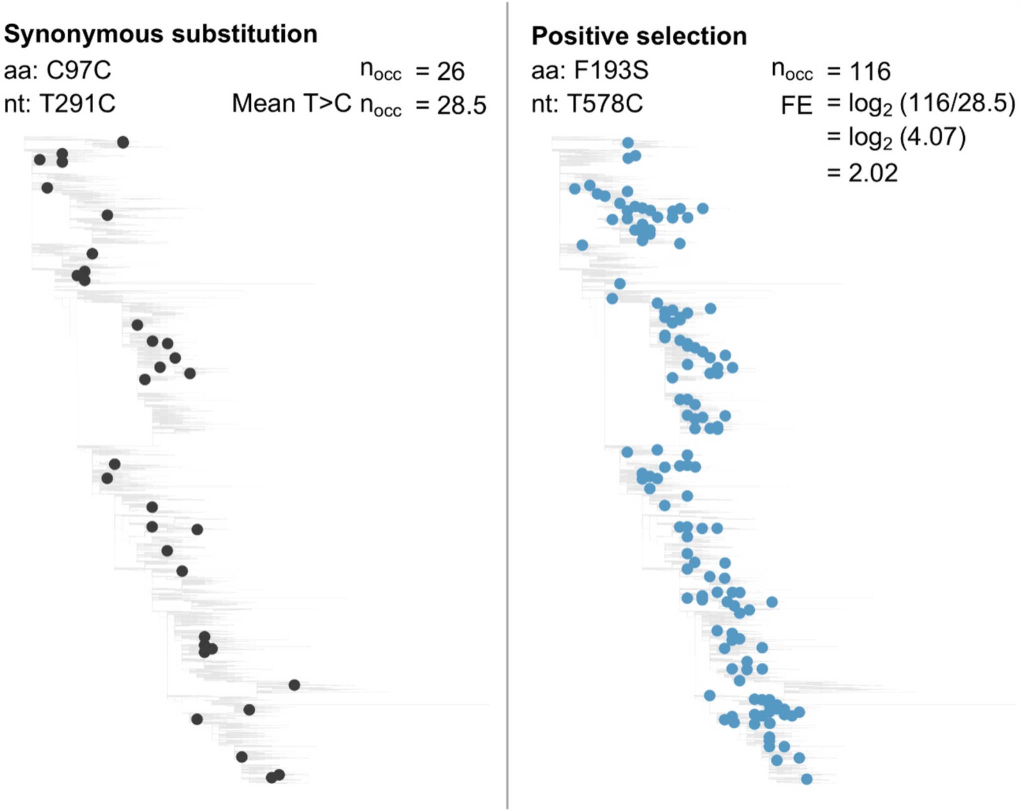


Similar to Bloom & Neher 2023 on SARS-CoV-2 https://doi.org/10.1093/ve/vead055

#### Top 15 substitutions at "Koel-7" positions:

| | | n / E(n) | FE |
|---|---|---|---|
| 1 | N 158 H | 35 / 4.1 | +3.1 |
| 2 | N 158 K | 129 / 22.0 | +2.6 |
| 3 | F 193 S | 147 / 35.3 | +2.1 |
| 4 | Y 159 F | 17 / 5.3 | +1.7 |
| 5 | N 158 S | 31 / 30.9 | +0.0 |
| 6 | N 158 T | 3 / 4.1 | -0.5 |
| 7 | K 189 R | 21 / 31.0 | -0.6 |
| 8 | H 156 Q | 15 / 22.2 | -0.6 |
| 9 | F 193 Y | 4 / 6.5 | -0.7 |
| 10 | K 189 M | 3 / 5.3 | -0.8 |
| 11 | S 145 R | 9 / 16.1 | -0.8 |
| 12 | N 158 D | 17 / 30.9 | -0.9 |
| 13 | K 189 N | 8 / 15.3 | -0.9 |
| 14 | S 145 N | 39 / 75.6 | -1.0 |
| 15 | S 145 G | 15 / 30.9 | -1.0 |

(Koel-7 = 145, 155, 156, 158, 159, 189, 193)

### All antigenic cluster transitions since 1987

| | | FE | rank |
|---|---|---|---|
| SI87 → BE89 | N 145 K | +5.0 | 1 |
| SI87 → BE92 | E 156 K | +4.5 | 3 |
| BE92 → WU95 | N 145 K | +5.5 | 1 |
| WU95 → SY97 | K 156 Q | +4.8 | 1 |
| WU95 → SY97 | E 158 K | +1.6 | 4 |
| SY97 → FU02 | Q 156 H | +2.3 | 6 |
| FU02 → CA04 | K 145 N | +3.6 | 2 |
| WI05 → PE09 | K 158 N | +1.9 | 3 |
| WI05 → PE09 | N 189 K | +0.7 | 6 |
| PE09 → SW13 | F 159 S | -2.9 | >4 |
| SW13 → KA17 | F 193 S | +2.5 | 2 |
| PE09 → HK14 | F 159 Y | +2.0 | 2 |
| HK14 → HK19 | F 193 S | +2.1 | 3 |
| HK14 → CA20 | F 193 S | +2.1 | 3 |
| CA20 → DA21 | Y 159 N | +2.8 | >3 |

Median rank of 3rd among Koel-7 substitutions

Too few CA20 seqs. to rank

### Convergent substitutions in current viruses

| | Overall | 2021/2 | 2022/3 | 2023/4 |
|---|---|---|---|---|
| S 145 N | +1.4<br>160/62 | +0.8<br>18/10.1 | +1.0<br>66/34 | +2.1<br>76/18 |
| N 158 K | +0.1<br>20/18.8 | -1.6<br>1/3 | -1.0<br>5/10.1 | +1.3<br>14/5.8 |
| K 189 R | +0.4<br>35/26.2 | -0.5<br>3/4.2 | -0.0<br>14/14 | +1.2<br>18/8 |
| S 145 R | -0.5<br>9/13.1 | -1.1<br>1/2.1 | -1.8<br>2/7.2 | +0.7<br>6/3.8 |
| N 159 S | -0.1<br>24/26.2 | +0.5<br>6/4.1 | -0.6<br>9/14.1 | +0.2<br>9/8.1 |
| S 193 A | +0.5<br>6/4.2 | +1.6<br>2/0.7 | +0.4<br>3/2.2 | -0.4<br>1/1.3 |

Fitness effect

observed # occurrences / expected # occurrences (neutral)

- Fitness effects (FE) calculated for Darwin/2021 antigenic cluster & descendants
  1. For April to April years
  2. Overall
- Showing: Koel-7 substitutions with +ve FE in 2023/4 or Overall

### Convergent substitutions in current viruses

| | Overall | 2021/2 | 2022/3 | 2023/4 |
|---|---|---|---|---|
| S 145 N | +1.4<br>160/62 | +0.8<br>18/10.1 | +1.0<br>66/34 | +2.1<br>76/18 |
| N 158 K | +0.1<br>20/18.8 | -1.6<br>1/3 | -1.0<br>5/10.1 | +1.3<br>14/5.8 |
| K 189 R | +0.4<br>35/26.2 | -0.5<br>3/4.2 | -0.0<br>14/14 | +1.2<br>18/8 |
| S 145 R | -0.5<br>9/13.1 | -1.1<br>1/2.1 | -1.8<br>2/7.2 | +0.7<br>6/3.8 |
| N 159 S | -0.1<br>24/26.2 | +0.5<br>6/4.1 | -0.6<br>9/14.1 | +0.2<br>9/8.1 |
| S 193 A | +0.5<br>6/4.2 | +1.6<br>2/0.7 | +0.4<br>3/2.2 | -0.4<br>1/1.3 |

- Fitness effects (FE) calculated for Darwin/2021 antigenic cluster & descendants
  1. For April to April years
  2. Overall
- Showing: Koel-7 substitutions with +ve FE in 2023/4 or Overall

### Convergent substitutions in current viruses

| | Overall | 2021/2 | 2022/3 | 2023/4 |
|---|---|---|---|---|
| S 145 N | +1.4<br>160/62 | +0.8<br>18/10.1 | +1.0<br>66/34 | +2.1<br>76/18 |
| N 158 K | +0.1<br>20/18.8 | -1.6<br>1/3 | -1.0<br>5/10.1 | +1.3<br>14/5.8 |
| K 189 R | +0.4<br>35/26.2 | -0.5<br>3/4.2 | -0.0<br>14/14 | +1.2<br>18/8 |
| S 145 R | -0.5<br>9/13.1 | -1.1<br>1/2.1 | -1.8<br>2/7.2 | +0.7<br>6/3.8 |
| N 159 S | -0.1<br>24/26.2 | +0.5<br>6/4.1 | -0.6<br>9/14.1 | +0.2<br>9/8.1 |
| S 193 A | +0.5<br>6/4.2 | +1.6<br>2/0.7 | +0.4<br>3/2.2 | -0.4<br>1/1.3 |

- Fitness effects (FE) calculated for Darwin/2021 antigenic cluster & descendants
  1. For April to April years
  2. Overall
- Showing: Koel-7 substitutions with +ve FE in 2023/4 or Overall

### Convergent substitutions in current viruses

| | Overall | 2021/2 | 2022/3 | 2023/4 |
|---|---|---|---|---|
| S 145 N | +1.4<br>160/62 | +0.8<br>18/10.1 | +1.0<br>66/34 | +2.1<br>76/18 |
| N 158 K | +0.1<br>20/18.8 | -1.6<br>1/3 | -1.0<br>5/10.1 | +1.3<br>14/5.8 |
| K 189 R | +0.4<br>35/26.2 | -0.5<br>3/4.2 | -0.0<br>14/14 | +1.2<br>18/8 |
| S 145 R | -0.5<br>9/13.1 | -1.1<br>1/2.1 | -1.8<br>2/7.2 | +0.7<br>6/3.8 |
| N 159 S | -0.1<br>24/26.2 | +0.5<br>6/4.1 | -0.6<br>9/14.1 | +0.2<br>9/8.1 |
| S 193 A | +0.5<br>6/4.2 | +1.6<br>2/0.7 | +0.4<br>3/2.2 | -0.4<br>1/1.3 |

- Fitness effects (FE) calculated for Darwin/2021 antigenic cluster & descendants
  1. For April to April years
  2. Overall
- Showing: Koel-7 substitutions with +ve FE in 2023/4 or Overall

### Convergent substitutions in current viruses

| | Overall | 2021/2 | 2022/3 | 2023/4 |
|---|---|---|---|---|
| S 145 N | +1.4<br>160/62 | +0.8<br>18/10.1 | +1.0<br>66/34 | +2.1<br>76/18 |
| N 158 K | +0.1<br>20/18.8 | -1.6<br>1/3 | -1.0<br>5/10.1 | +1.3<br>14/5.8 |
| K 189 R | +0.4<br>35/26.2 | -0.5<br>3/4.2 | -0.0<br>14/14 | +1.2<br>18/8 |
| S 145 R | -0.5<br>9/13.1 | -1.1<br>1/2.1 | -1.8<br>2/7.2 | +0.7<br>6/3.8 |
| N 159 S | -0.1<br>24/26.2 | +0.5<br>6/4.1 | -0.6<br>9/14.1 | +0.2<br>9/8.1 |
| S 193 A | +0.5<br>6/4.2 | +1.6<br>2/0.7 | +0.4<br>3/2.2 | -0.4<br>1/1.3 |

- Fitness effects (FE) calculated for Darwin/2021 antigenic cluster & descendants
  1. For April to April years
  2. Overall
- Showing: Koel-7 substitutions with +ve FE in 2023/4 or Overall

Additional slides

### Convergent substitutions in current viruses at all HA1 positions

(2023/4 data)

| | | n / E(n) | FE |
|---|---|---|---|
| 1 | K 207 Q | 18 / 1.1 | +4.1 |
| 2 | F 79 V | 5 / 1.2 | +2.1 |
| 3 | S 145 N | 76 / 18.0 | +2.1 |
| 4 | N 63 D | 25 / 8.0 | +1.7 |
| 5 | I 48 R | 4 / 1.3 | +1.6 |
| 6 | T 10 M | 24 / 8.0 | +1.6 |
| 7 | I 25 V | 20 / 6.7 | +1.6 |
| 8 | K 278 M | 4 / 1.4 | +1.5 |
| 9 | T 135 A | 23 / 7.9 | +1.5 |
| 10 | Q 173 H | 7 / 2.4 | +1.5 |
| 11 | Q 197 H | 7 / 2.5 | +1.5 |
| 12 | K 207 R | 20 / 7.9 | +1.3 |
| 13 | N 158 K | 14 / 5.8 | +1.3 |
| 14 | I 242 M | 19 / 7.9 | +1.3 |
| 15 | I 160 M | 19 / 8.0 | +1.2 |
| 16 | S 198 A | 3 / 1.3 | +1.2 |
| 17 | S 137 A | 3 / 1.3 | +1.2 |
| 18 | I 160 R | 3 / 1.3 | +1.2 |
| 19 | K 189 R | 18 / 8.0 | +1.2 |
| 20 | I 260 M | 18 / 8.0 | +1.2 |
| 21 | N 45 I | 3 / 1.4 | +1.1 |
| 22 | T 135 K | 11 / 5.4 | +1.0 |
| 23 | K 238 N | 5 / 2.5 | +1.0 |
| 24 | D 271 E | 11 / 5.6 | +1.0 |
| 25 | S 124 R | 13 / 6.7 | +1.0 |
| 26 | N 94 H | 2 / 1.1 | +0.9 |
| 27 | N 165 T | 2 / 1.1 | +0.9 |
| 28 | N 81 D | 15 / 8.1 | +0.9 |
| 29 | K 264 T | 2 / 1.1 | +0.9 |
| 30 | S 279 F | 15 / 8.1 | +0.9 |
| 31 | L 15 I | 10 / 5.5 | +0.9 |
| 32 | P 239 S | 14 / 7.8 | +0.9 |
| 33 | I 214 T | 17 / 9.5 | +0.8 |
| 34 | K 278 N | 7 / 3.9 | +0.8 |
| 35 | R 208 I | 6 / 3.5 | +0.8 |
| 36 | R 261 L | 6 / 3.6 | +0.8 |
| 37 | L 177 M | 3 / 1.8 | +0.7 |
| 38 | A 212 S | 6 / 3.6 | +0.7 |
| 39 | E 280 G | 13 / 8.1 | +0.7 |
| 40 | S 146 G | 13 / 8.1 | +0.7 |
| 41 | S 144 N | 31 / 19.5 | +0.7 |
| 42 | S 145 R | 6 / 3.8 | +0.7 |
| 43 | S 115 A | 2 / 1.3 | +0.7 |
| 44 | N 165 K | 9 / 5.8 | +0.6 |
| 45 | S 114 A | 2 / 1.3 | +0.6 |
| 46 | S 199 A | 2 / 1.3 | +0.6 |
| 47 | I 274 M | 2 / 1.3 | +0.6 |
| 48 | N 53 Y | 2 / 1.3 | +0.6 |
| 49 | R 33 Q | 29 / 19.3 | +0.6 |
| 50 | I 25 M | 10 / 6.7 | +0.6 |

Multiple convergent substitutions at same position:

| | FE | n / E(n) | Gly. |
|---|---|---|---|
| I 25 V | +1.6 | 20 / 6.7 | |
| I 25 M | +0.6 | 10 / 6.7 | |
| F 79 V | +2.1 | 5 / 1.2 | |
| F 79 L | +0.5 | 16 / 11.5 | |
| S 124 R | +1.0 | 13 / 6.7 | |
| S 124 N | +0.4 | 26 / 19.2 | |
| T 135 A | +1.5 | 23 / 7.9 | - |
| T 135 K | +1.0 | 11 / 5.4 | - |
| I 160 M | +1.2 | 19 / 8.0 | |
| I 160 R | +1.2 | 3 / 1.3 | |
| K 207 Q | +4.1 | 18 / 1.1 | |
| K 207 R | +1.3 | 20 / 7.9 | |
| K 278 M | +1.5 | 4 / 1.4 | |
| K 278 N | +0.8 | 7 / 3.9 | |
