## Supplementary material for "Predicting the antigenic evolution of seasonal influenza viruses using phylogenetic convergence": Data S2

### Information for TC1 prior to the WHO February 2025 NH Influenza Vaccines Consultation Meeting (VCM)

H3 convergent evolution

17th December 2024

Center for Pathogen Evolution

University of Cambridge, United Kingdom

### Fitness effect (FE) measurements

(here F193S in Hong Kong/4801/2014-like viruses)

observed # occurrences

expected # occurrences

**Synonymous substitution**

aa: C97C

nt: T291C

$n_{occ} = 26$

Mean T>C $n_{occ} = 28.5$

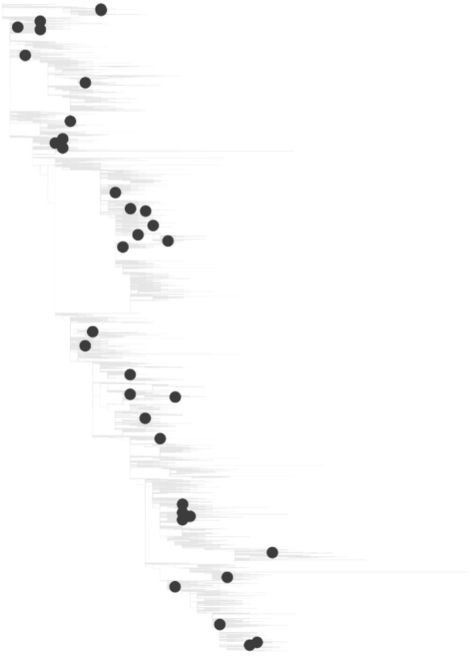

**Positive selection**

aa: F193S

nt: T578C

$n_{occ} = 116$

$FE = \log_2(116/28.5) = \log_2(4.07) = 2.02$

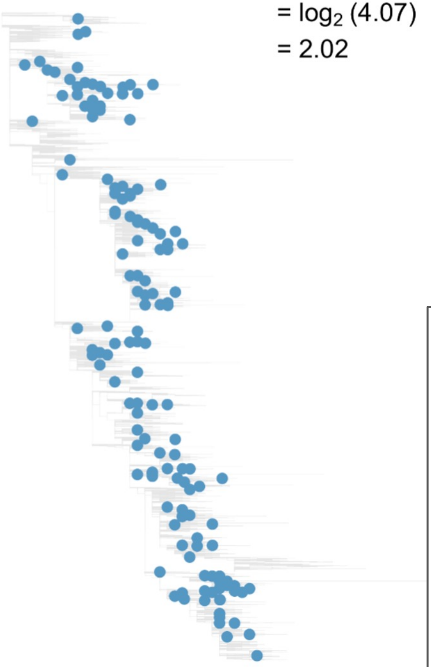

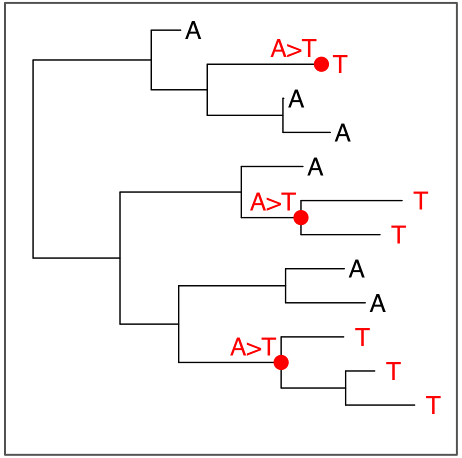


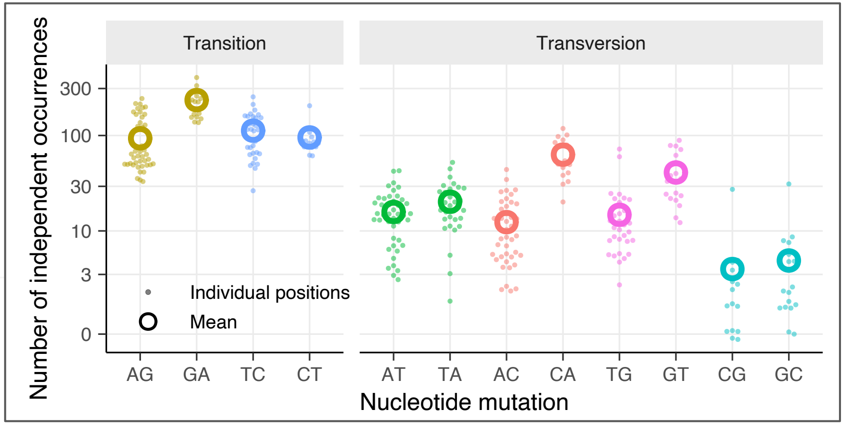


Similar to Bloom & Neher 2023 on SARS-CoV-2 https://doi.org/10.1093/ve/vead055

### Convergent substitutions in recent viruses

| | Overall | 2021/2 | 2022/3 | 2023/4 |
|---|---|---|---|---|
| S 145 N | +1.4<br>160/62 | +0.8<br>18/10.1 | +1.0<br>66/34 | +2.1<br>76/18 |
| N 158 K | +0.1<br>20/18.8 | -1.6<br>1/3 | -1.0<br>5/10.1 | +1.3<br>14/5.8 |
| K 189 R | +0.4<br>35/26.2 | -0.5<br>3/4.2 | -0.0<br>14/14 | +1.2<br>18/8 |
| S 145 R | -0.5<br>9/13.1 | -1.1<br>1/2.1 | -1.8<br>2/7.2 | +0.7<br>6/3.8 |
| N 159 S | -0.1<br>24/26.2 | +0.5<br>6/4.1 | -0.6<br>9/14.1 | +0.2<br>9/8.1 |
| S 193 A | +0.5<br>6/4.2 | +1.6<br>2/0.7 | +0.4<br>3/2.2 | -0.4<br>1/1.3 |

Fitness effect

observed # occurrences / expected # occurrences

Fitness effects calculated for Darwin/2021 antigenic cluster & descendants

1. For April to April years
2. Overall

Showing Koel-7 substitutions with +ve FE in **2023/4 or Overall**

Phylogenetic tree constructed with **FastTree** using all sequences from GISAID up to **1st April 2024**.

### Convergent substitutions in recent viruses

| | Overall | 2021/2 | 2022/3 | 2023/4 | 2024/5 | Frequency since April 2024 |
|---|---|---|---|---|---|---|
| S 145 N | +1.6<br>287/92.2 | +0.7<br>20/12.5 | +1.0<br>77/38 | +2.2<br>115/25.2 | +2.8<br>70/10.2 | 21% |
| N 158 K | +1.0<br>45/22 | -1.5<br>1/2.8 | -0.3<br>7/8.7 | +1.4<br>16/6.1 | +1.8<br>10/2.9 | 1.4% |
| K 189 R | +0.7<br>55/34.4 | -0.6<br>3/4.5 | -0.1<br>13/13.6 | +1.2<br>22/9.6 | +1.7<br>15/4.5 | 0.8% |
| S 145 R | -0.1<br>14/14.6 | -1.0<br>1/2 | -1.0<br>3/6 | +0.8<br>7/4 | +0.9<br>3/1.6 | 0.3% |
| N 159 S | +0.2<br>36/31.5 | +0.3<br>5/4.1 | -0.5<br>9/13.1 | +0.7<br>16/9.7 | +0.4<br>6/4.6 | 0.1% |
| S 193 T | -0.8<br>4/6.9 | +0.1<br>1/0.9 | -∞<br>0/2.8 | +0.6<br>3/2 | -∞<br>0/0.9 | 0% |

Fitness effects calculated for Darwin/2021 antigenic cluster & descendants

1. For April to April years
2. Overall

Showing Koel-7 substitutions with +ve FE in **2023/4 or 2024/5**

Phylogenetic tree constructed with **CMAPLE** using all sequences from GISAID up to **3rd December 2024**.
