## Supplementary material for "Predicting the antigenic evolution of seasonal influenza viruses using phylogenetic convergence": Data S3

### Information for TC2 prior to the WHO February 2025 NH Influenza Vaccines Consultation Meeting (VCM)

H3 convergent evolution

30th January 2025

Center for Pathogen Evolution

University of Cambridge, United Kingdom

### Fitness effect (FE) measurements

(here F193S in Hong Kong/4801/2014-like viruses)

observed # occurrences

expected # occurrences

**Synonymous mutation**

aa: C97C — $n_{occ}$ = 26

nt: T291C — Mean T>C $n_{occ}$ = 28.5

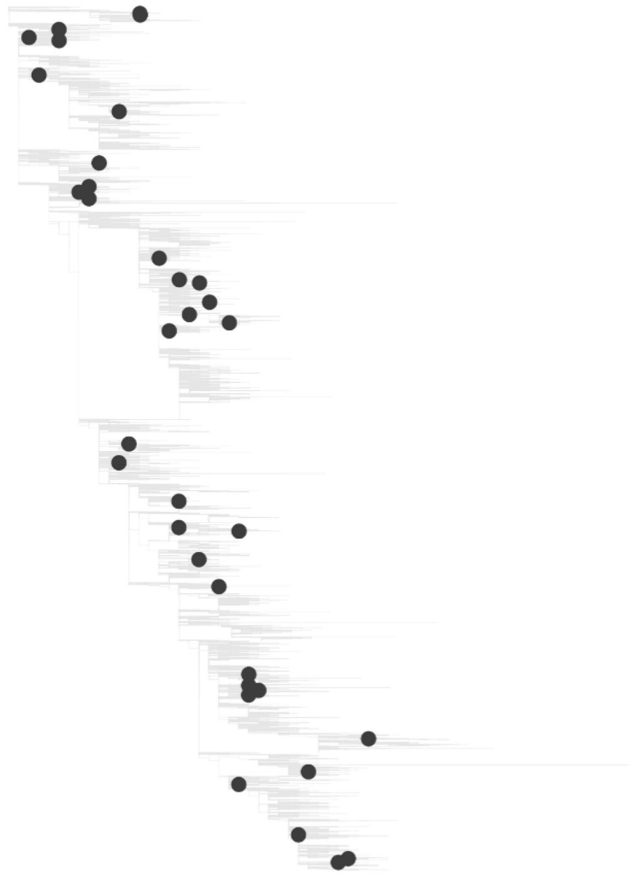

**Positive selection**

aa: F193S — $n_{occ}$ = 116

nt: T578C

$$FE = \log_2 (116/28.5) = \log_2 (4.07) = 2.02$$

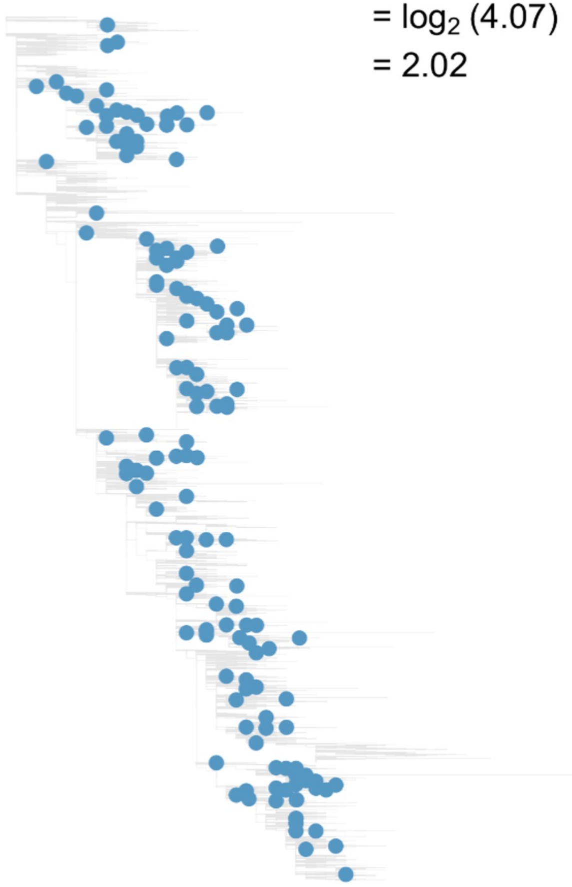

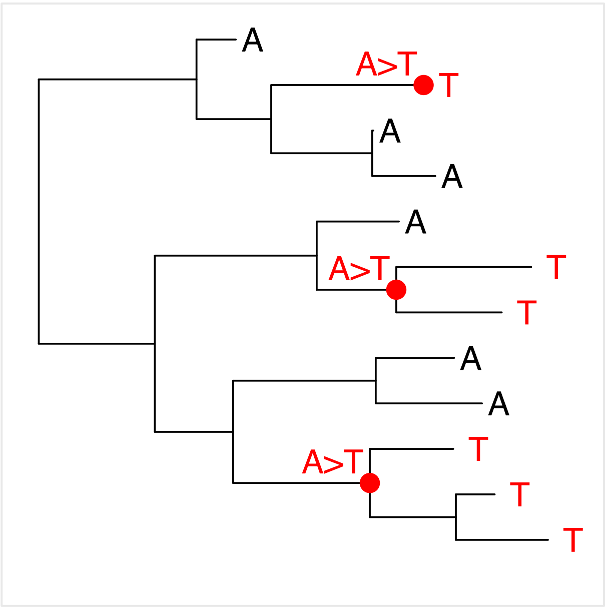


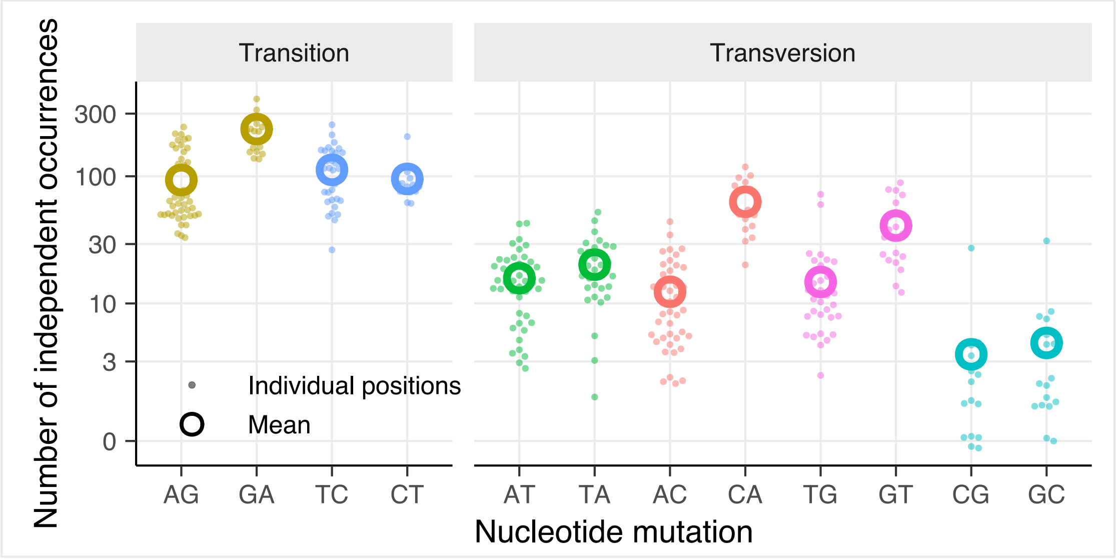


Similar to Bloom & Neher 2023 on SARS-CoV-2 https://doi.org/10.1093/ve/vead055

### Convergent substitutions in recent viruses

| | Overall | 2021/2 | 2022/3 | 2023/4 | 2024/5 | |
|---|---|---|---|---|---|---|
| S 145 N | +1.7<br>276/86.76 | +0.6<br>19/12.14 | +1.1<br>76/36.49 | +2.1<br>103/24.74 | +2.5<br>78/13.39 | 24% |
| N 158 K | +0.7<br>33/20.84 | -1.5<br>1/2.74 | -0.7<br>5/8.28 | +1.3<br>15/5.99 | +1.7<br>12/3.82 | 2.2% |
| K 189 R | +0.8<br>59/32.95 | -0.5<br>3/4.35 | -0.0<br>13/13.14 | +1.4<br>25/9.51 | +1.6<br>18/5.94 | 2.2% |
| N 158 H | +1.3<br>9/3.63 | -∞<br>0/0.48 | +1.8<br>5/1.44 | +0.9<br>2/1.04 | +1.6<br>2/0.67 | 0.02% |
| S 145 R | +0.0<br>14/13.61 | -0.9<br>1/1.91 | -2.5<br>1/5.73 | +0.9<br>7/3.88 | +1.3<br>5/2.1 | 0.02% |
| N 159 S | +0.2<br>38/32.32 | +0.3<br>5/3.98 | -0.7<br>8/12.64 | +0.6<br>15/9.59 | +0.7<br>10/6.11 | 0.2% |

Frequency since October 2024

Fitness effects calculated for Darwin/2021 antigenic cluster & descendants for:

1. April to April years
2. 1st April 2021 to present

Showing Koel-7 substitutions with +ve FE in **2024/5**

Phylogenetic tree constructed with CMAPLE using all sequences from GISAID up to **28th January 2025**.

### Convergent substitutions in recent viruses

| | Overall | 2021/2 | 2022/3 | 2023/4 | 2024/5 | |
|---|---|---|---|---|---|---|
| S 145 N | +1.7<br>276/86.76 | +0.6<br>19/12.14 | +1.1<br>76/36.49 | +2.1<br>103/24.74 | +2.5<br>78/13.39 | 24% |
| N 158 K | +0.7<br>33/20.84 | -1.5<br>1/2.74 | -0.7<br>5/8.28 | +1.3<br>15/5.99 | +1.7<br>12/3.82 | 2.2% |
| K 189 R | +0.8<br>59/32.95 | -0.5<br>3/4.35 | -0.0<br>13/13.14 | +1.4<br>25/9.51 | +1.6<br>18/5.94 | 2.2% |

Frequency since October 2024

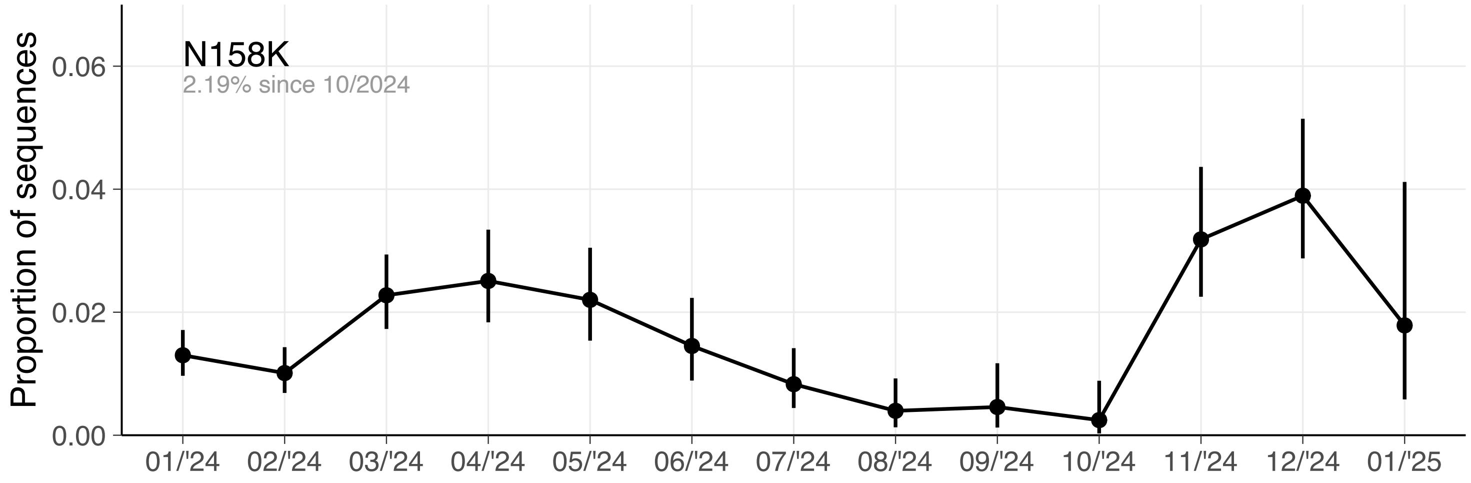


Fitness effects calculated for Darwin/2021 antigenic cluster & descendants for:

1. April to April years
2. 1st April 2021 to present

Showing Koel-7 substitutions with +ve FE in **2024/5**

Phylogenetic tree constructed with CMAPLE using all sequences from GISAID up to **28th January 2025**.

### Convergent substitutions in *very* recent viruses

| | Overall | 01/2024 | 04/2024 | 07/2024 | 10/2024 | Frequency since October 2024 |
|---|---|---|---|---|---|---|
| S 145 N | +2.4<br>135/24.9 | +2.3<br>57/11.56 | +2.3<br>30/6.12 | +3.1<br>34/3.97 | +2.1<br>14/3.25 | 24% |
| N 158 K | +1.6<br>20/6.71 | +1.5<br>8/2.9 | +2.2<br>7/1.56 | +1.2<br>3/1.28 | +1.1<br>2/0.96 | 2.2% |
| K 189 R | +1.6<br>33/10.55 | +1.7<br>15/4.62 | +1.5<br>7/2.47 | +1.6<br>6/1.93 | +1.7<br>5/1.52 | 2.2% |
| N 158 H | +0.8<br>2/1.17 | -∞<br>0/0.51 | +1.9<br>1/0.27 | -∞<br>0/0.22 | +2.6<br>1/0.17 | 0.02% |
| S 145 R | +1.6<br>12/3.9 | +2.0<br>7/1.81 | +1.6<br>3/0.96 | +0.7<br>1/0.62 | +1.0<br>1/0.51 | 0.02% |
| N 159 S | +1.0<br>21/10.76 | +1.2<br>11/4.67 | +1.3<br>6/2.5 | -1.0<br>1/2 | +0.9<br>3/1.59 | 0.2% |

Fitness effects calculated for Darwin/2021 antigenic cluster & descendants for:

1. Quarters since 1st Jan 2024
2. 1st Jan 2024 to present

Showing Koel-7 substitutions with +ve FE in **2024/5**

Phylogenetic tree constructed with CMAPLE using all sequences from GISAID up to **28th January 2025**.

### New occurrences of N158K and K189R (since July 2024)

**N158K (5x new occurrences)**

N158K + N145S **K189R** S378N
Dec. 2024 to Jan. 2025
12x NETHERLANDS

Relative to J.2+S145N (recent SH vaccine recommendation)

N158K + I535T
1x NORWAY

N158K + N145S S312N A530V
1x CAMBODIA

N158K + T65K S124N N145S L532V
1x AUSTRALIA

N158K + T65K S124N N145S
1x AUSTRALIA

Separated by 4x synonymous mutations

**K189R (11x new occurrences)**

K189R + T135K N145S
Jul. 2024 to Dec. 2024
7x USA
5x UK
3x BHUTAN
2x CANADA
1x CROATIA
1x MALAYSIA
1x SINGAPORE
1x THAILAND

N145S K189R S378N
21x BRAZIL

K189R + N145S
12x NETHERLANDS
7x BRAZIL
1x GUATEMALA
1x PORTUGAL

K189R + N63D N145S V309I
11x AUSTRALIA

K189R + N63D N145S V347M
9x BRAZIL

K189R + N145S D291N T301A S378N
9x PERU

K189R + I67V V112I N145S I529V
8x CANADA
1x UK

K189R + N145S
3x PERU

K189R + N145S V223I
1x USA

K189R + F79L N145S P239S V347M
1x NETHERLANDS

K189R + N145S V223I
1x USA

### Additional slides

# N158K + N145S K189R S378N

| Isolate name | Collection date | AA diff. | Syn diff. |
|---|---|---|---|
| NETHERLANDS/10685/2024 | 12/12/2024 | | C495T |
| NETHERLANDS/2093/2024 | 17/12/2024 | | G555A |
| NETHERLANDS/10706/2024 | 23/12/2024 | N126K | G555A |
| NETHERLANDS/2182/2024 | 25/12/2024 | G504E | |
| NETHERLANDS/2121/2024 | 28/12/2024 | N126K | |
| NETHERLANDS/2175/2024 | 29/12/2024 | P4H | G555A |
| NETHERLANDS/10734/2024 | 31/12/2024 | G5R | |
| NETHERLANDS/24/2025 | 01/01/2025 | D408N | G555A |
| NETHERLANDS/10008/2025 | 03/01/2025 | | T165C |
| NETHERLANDS/77/2025 | 06/01/2025 | | |
| NETHERLANDS/10080/2025 | 08/01/2025 | S262N | G555A |
| NETHERLANDS/10024/2025 | 09/01/2025 | | G555A |

# K189R + T135K N145S

| Isolate name | Collection date | AA diff. | Syn diff. |
|---|---|---|---|
| PHUKET/P3184/2024 | 15/08/2024 | | |
| ENGLAND/3940090/2024 | 05/09/2024 | | |
| BHUTAN/1657/2024 | 13/09/2024 | | |
| BHUTAN/1699/2024 | 30/09/2024 | | |
| BHUTAN/1478/2024 | 30/09/2024 | | |
| WASHINGTON/284/2024 | 02/11/2024 | | |
| MALAYSIA/IMRSARI2716/2024 | 06/11/2024 | | G1530A |
| WASHINGTON/286/2024 | 18/11/2024 | | |
| BRITISH_COLUMBIA/RV05533/2024 | 26/11/2024 | | |
| BRITISH_COLUMBIA/RV05532/2024 | 26/11/2024 | | |
| CROATIA/HZJZ8199/2024 | 27/11/2024 | | |
| ENGLAND/4961005/2024 | 30/11/2024 | | G72A |
| ENGLAND/4921120/2024 | 02/12/2024 | | G72A |
| UTAH/2051439/2024 | 12/12/2024 | | G126A C684T |
| COLORADO/ISC1492/2024 | 13/12/2024 | | |
| COLORADO/ISC1479/2024 | 13/12/2024 | | |
| COLORADO/ISC1493/2024 | 13/12/2024 | | |
| COLORADO/ISC1494/2024 | 14/12/2024 | | |
| ENGLAND/5141055/2024 | 17/12/2024 | | G72A T1146C |
| ENGLAND/5160227/2024 | 17/12/2024 | | |
| SINGAPORE/GP20238/2024 | 26/12/2024 | | C318A G1077A |

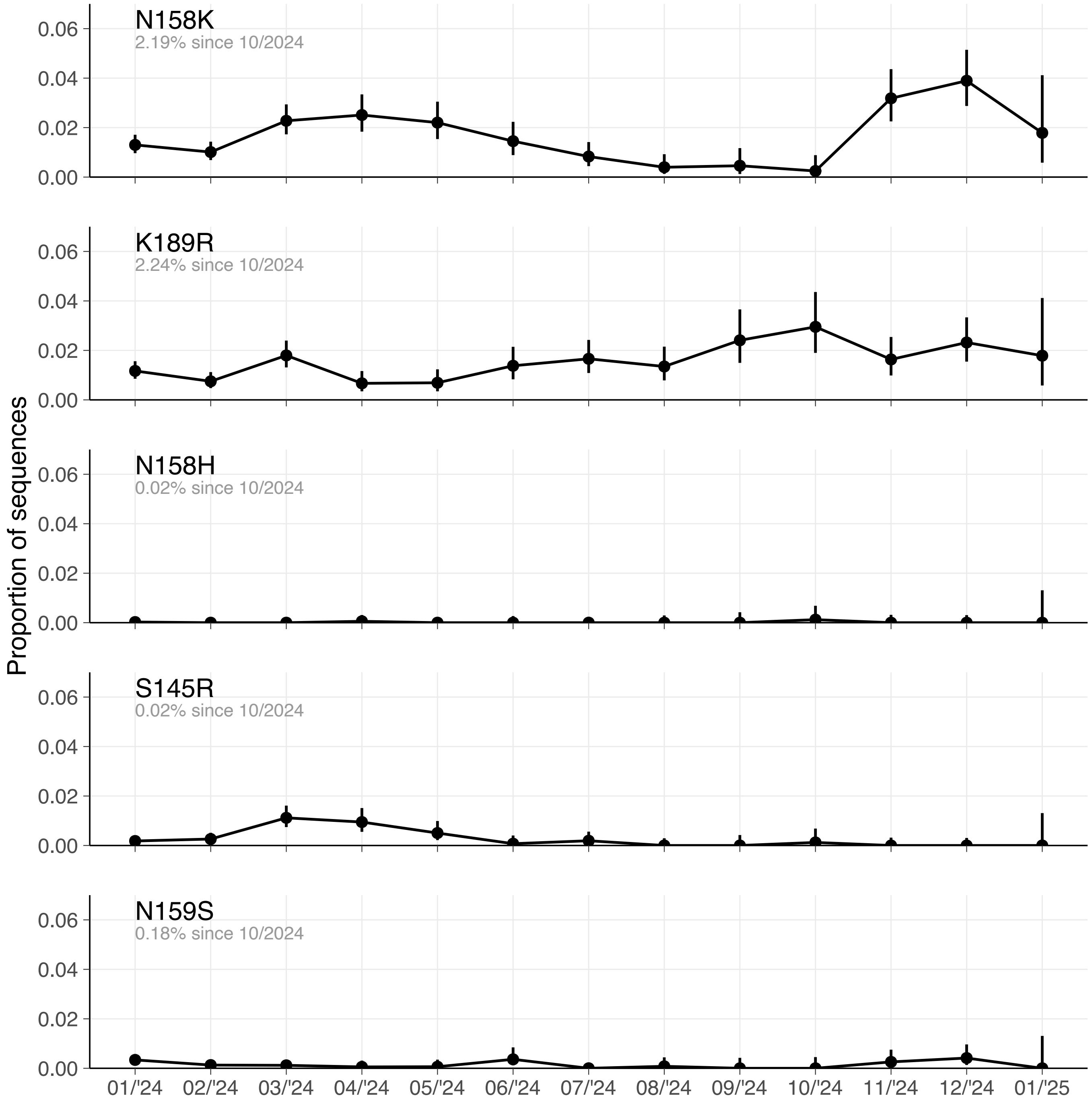

Proportion of sequences
N158K
2.19% since 10/2024
K189R
2.24% since 10/2024
N158H
0.02% since 10/2024
S145R
0.02% since 10/2024
N159S
0.18% since 10/2024
0.06
0.04
0.02
0.00
01/'24
02/'24
03/'24
04/'24
05/'24
06/'24
07/'24
08/'24
09/'24
10/'24
11/'24
12/'24
01/'25
